# Hijacking a bacterial membrane transporter for efficient genetic code expansion

**DOI:** 10.1101/2025.03.06.641855

**Authors:** Tarun Iype, Maximilian Fottner, Paul Böhm, Carlos Piedrafita, Kathrin Lang

## Abstract

The site-specific encoding of non-canonical amino acids (ncAAs) provides a powerful tool for expanding the functional repertoire of proteins. Its widespread use for basic research and biotechnological applications is, however, hampered by low efficiencies of current ncAA incorporation strategies. We uncover poor cellular ncAA uptake as a main obstacle to efficient genetic code expansion and overcome this bottleneck by hijacking a bacterial membrane transporter to actively import isopeptide-linked ncAAs within easily synthesizable tripeptide-scaffolds. Using this approach, we enable efficient encoding of eleven previously inaccessible ncAAs, decorating proteins with bioorthogonal and crosslinker moieties, posttranslational modifications, and functionalities for chemoenzymatic conjugation. To enhance scalability of protein production, we evolve the membrane transporter for preferential import of isopeptide-linked tripeptides, creating a novel *Escherichia coli* strain that facilitates single and multi-site ncAA incorporation with wild type efficiencies. Additionally, we adapt the tripeptide-scaffolds for co-transport of two different ncAAs, enabling their efficient dual incorporation. This work underscores the importance of optimizing ncAA-uptake for high-yielding production of modified proteins and will accelerate the development of generalizable transport systems, aiding incorporation of non-canonical building blocks to broaden the chemical space of proteins without the need to design for passive membrane permeability.

## INTRODUCTION

Co-translational incorporation of non-canonical amino acids (ncAAs) into proteins via genetic code expansion (GCE) has become a versatile approach to increase the chemical space of the proteome. By leveraging orthogonal aminoacyl-tRNA synthetases (aaRSs), a plethora of diverse functionalities^1^ have been incorporated into proteins of interest (POIs) across all domains of life, most commonly in response to the amber nonsense codon (amber suppression). Site-specific encoding of functionalities such as post-translational modifications (PTMs), bioorthogonal handles, crosslinking moieties, spectroscopic probes, and photocaged amino acids offers a wide range of tools for studying and modifying protein functions and generating proteins with potential therapeutic and biotechnological significance.^2,3^

While GCE holds promise in a wide range of fields, its widespread use in commercial and research applications remains limited due to several challenges, most notably low protein production yields. This stems from insufficient enzymatic activities of orthogonal aaRSs towards many ncAA substrates, as well as unfavorable competition of aminoacylated tRNAs with release factors, causing premature translational termination at introduced nonsense codons.^4,5^ In addition, many ncAAs require advanced expertise in chemical synthesis, or are prohibitively expensive, and are used at high millimolar concentrations in typical expression experiments, a factor that is exacerbated by the low incorporation yields.

Significant advances have been made in recent years towards addressing these challenges. Optimization of aaRS/tRNA expression systems combined with novel selection and evolution strategies to improve both orthogonal aaRSs and tRNAs have considerably increased suppression efficiencies^6–10^. Efforts to bypass competition with release factors have led to orthogonal ribosomes that read quadruplet codons^11,12^, release factor knockout strains^13,14^ and recoded strains with compressed genetic codes^15,16^, allowing also for sense codon reassignment^17^.

Another less investigated factor impacting ncAA incorporation efficiencies lies in their intracellular bioavailability. In typical GCE experiments, chemically synthesized ncAAs are added to expression media and enter cells via passive diffusion, resulting in intracellular concentrations at best equal to, but most likely severely below those added to media. This specifically reduces aminoacylation efficiencies of aaRS/ncAA combinations that are operating below saturating conditions^4^. Furthermore, the reliance on passive diffusion limits the full potential of introducing new-to-nature functionalities, as cell-permeability needs to be considered when designing new ncAAs. For a handful of ncAAs, efforts in introducing and engineering enzymatic pathways to biosynthesize corresponding ncAAs directly within cells have resulted in increased incorporation efficiencies compared to exogenous ncAA addition^18–23^. Despite being an attractive prospect for GCE, introducing novel metabolic pathways into new hosts may require significant development. In addition, many ncAA functionalities lack biosynthetic strategies, necessitating substantial advancements before such approaches can be widely applied for increasing ncAA bioavailability and thus genetic encoding.

Engineering membrane transport systems as an alternative strategy for boosting intracellular ncAA uptake, is relatively unexplored, but holds the potential of being widely applicable for increasing expression yields of modified proteins. Previous research in this direction has investigated mutants of a periplasmic leucine binding protein to improve uptake and incorporation of various known ncAAs ^24^. Another study has used a ‘trojan-horse’ approach, in which a sulfonic acid moiety attached to a cargo molecule serves as recognition motif for a promiscuous bacterial sulfonate importer. Attaching an impermeant ncAA to a sulfonate carrier, followed by cleavage of the carrier via an engineered enzyme, led to increased cytosolic concentrations of this ncAA, but incorporation into proteins was not shown due to lack of a specific aaRS^25^. The use of peptide transporters has also been investigated^26,27^, most notably for transporting phosphotyrosine as a lysine-linked dipeptide^28^, which was believed to be a substrate for the dipeptide transporter Dpp. Peptide transporters, especially ATP-binding cassette (ABC) transporters like the Dpp-transporter, are promising candidates for engineering ncAA uptake due to their substrate promiscuity and ability to maintain high concentration gradients across the membrane^29^. In addition, attaching ncAAs to peptide carriers is easily achievable via solid-phase peptide synthesis (SPPS) requiring minimal chemical expertise.

Here we leverage a propeptide-based strategy combined with the engineering of a bacterial ABC-transporter system to boost intracellular ncAA concentrations and expression yields of modified proteins. We find that easily synthesized isopeptide-linked tripeptides (G-XisoK) are actively transported into *E. coli* cells via the oligopeptide permease transporter Opp and are processed in the cytosol to reveal dipeptidic XisoK ncAAs. These isopeptide-linked lysine derivatives allow for the incorporation of various functionalities available for GCE, with efficiencies rivaling wild type (wt) protein expressions, providing a toolbox of eleven novel ncAAs, covering approaches from bioorthogonal labeling over chemical and photocrosslinking to native chemical ligation and chemoenzymatic conjugation. To exploit efficient G-XisoK-uptake for cheap and scalable production of modified proteins, we evolve the periplasmic binding protein of the Opp transporter via a fluorescence-activated cell sorting platform, facilitating the preferential transport of G-XisoK tripeptides over linear tripeptides present in expression media. By introducing the evolved Opp binding protein variant into the *E. coli* genome, we create a novel *E. coli* strain that allows high XisoK incorporation efficiencies across a range of target proteins for both single as well as multiple XisoK encoding. Furthermore, we adapt the tripeptide scaffold for incorporation of two different ncAAs in response to two different nonsense codons via their concomitant transport using a single tripeptide, demonstrating the significant impact that active ncAA uptake has on the efficient production of proteins with an expanded alphabet.

## RESULTS AND DISCUSSION

### Isopeptide-linked tripeptides are privileged scaffolds for efficient *E. coli* uptake

Previous work in our group has pioneered the use of transpeptidases in combination with genetic code expansion to generate defined protein-protein conjugates. Through site-specific incorporation of an ncAA bearing an azide-caged dipeptidic nucleophile acceptor (AzGGisoK, Fig. S1a) and subsequent Staudinger reduction, GGisoK-modified proteins can engage in transpeptidation with donor proteins modified at their C-terminus with an appropriate recognition sequence. We have leveraged these approaches for successfully generating ubiquitin (Ub)- and Ub-like modifier (Ubl)-POI conjugates using sortase or *Oa*AEP1 as transpeptidases (Fig. S1a)^30–32^. In our quest to diversify the linker sequence in the generated protein conjugates, we explored the site-specific incorporation of ncAAs resembling a general G-XisoK scaffold (Fig. 1a). While AzGGisoK required intensive PylRS engineering, we were not able to identify a PylRS-variant for direct GGisoK incorporation^30^. In contrast, addition of the alanine-bearing G-XisoK tripeptide (G-AisoK, Fig. 1a) to growing *E. coli* cells (K12 strain), transformed with plasmids coding for the wild-type *Methanosarcina barkeri* Pyrrolysine-tRNA synthetase/tRNA pair (wt-*Mb*PylRS/PylT) and super-folder green fluorescence protein bearing an amber codon at position 150 (sfGFP-N150TAG), led to highly efficient sfGFP expression, comparable to wt-sfGFP production and amber suppression yields using the gold-standard ncAA BocK (Fig. 1a,b). Interestingly, MS-analysis of purified sfGFP expressed in presence of G-AisoK revealed site-specific incorporation of AisoK (Fig 1c). This suggests that the N-terminal glycine is cleaved off intracellularly, either on the free ncAA, or co/post-translationally. On the other hand, supplementing K12 with AisoK instead of G-AisoK showed almost no visible amber suppression and sfGFP production (Fig. 1b). This is corroborated by live-cell sfGFP-fluorescence measurements in the presence of different ncAAs. Cells grown in the presence of AisoK showed minimal fluorescence, while fluorescence is observed at earlier timepoints and reaches higher values with G-AisoK when compared to BocK (Fig. 1d). Similarly, G-AisoK-mediated AisoK incorporation was observed for other amber codon-bearing target proteins (Fig. S1b). Intrigued by this observation, we set out to determine intracellular levels of the corresponding ncAAs, using LC-MS based uptake assays^18,33^. Upon G-AisoK addition, we could not observe any intracellular G-AisoK, but AisoK accumulated at 5-10-fold higher concentrations compared to supplementing K12 cells with AisoK itself (Fig 1e, Fig. S1c).

**Figure 1.**
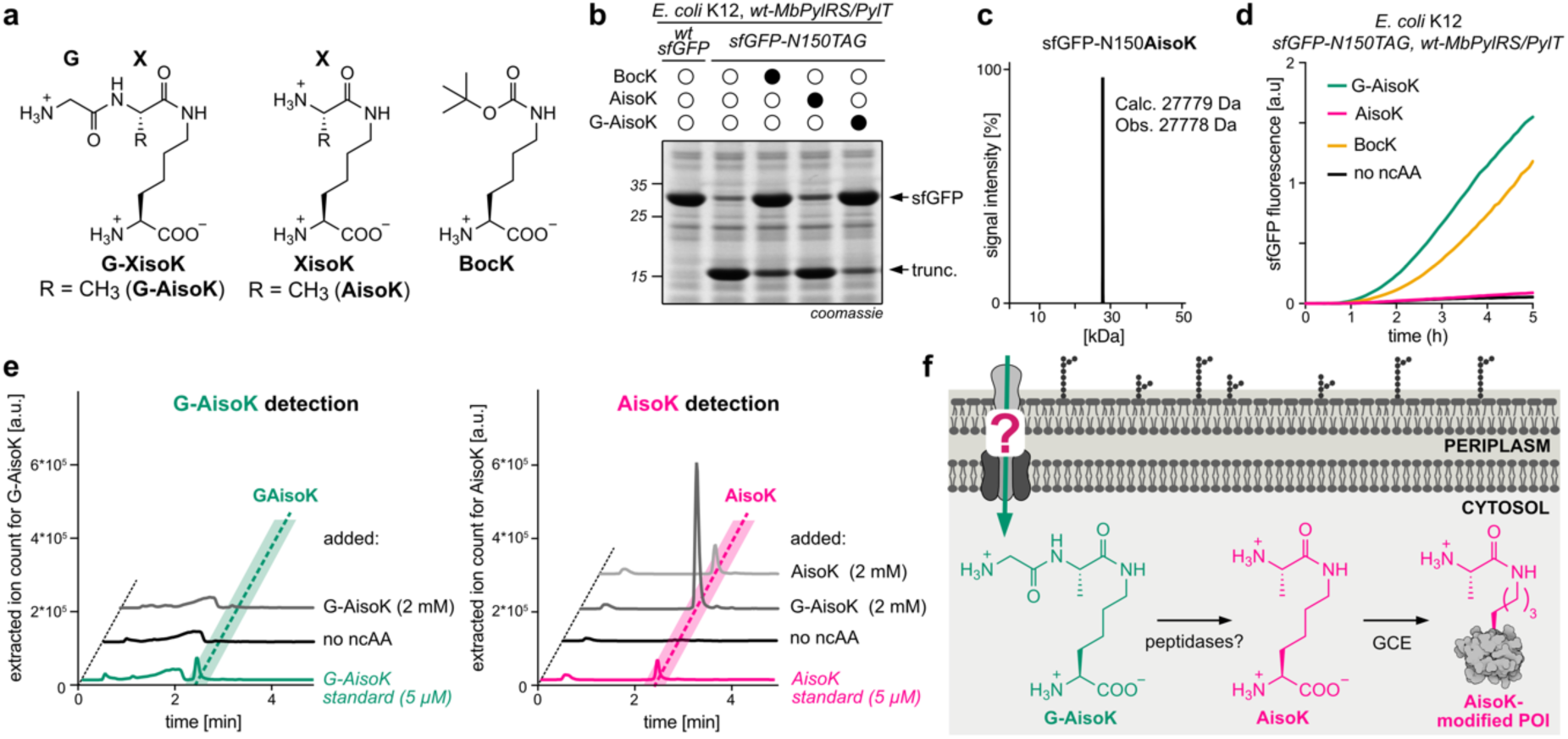
Isopeptide-linked tripeptides are privileged scaffolds for efficient *E. coli* uptake **a.** Chemical structures for G-XisoK, XisoK and BocK. X is alanine for G-AisoK and AisoK. **b.** SDS-PAGE analysis of sfGFP-N150TAG expression using wt-*Mb*PylRS/PylT in the presence of 2 mM BocK, AisoK or G-AisoK. (trunc. denotes truncated protein). Consistent results were obtained over three distinct replicate experiments. **c.** LC-MS analysis of sfGFP-N150AisoK. **d.** Time course measurements of sfGFP fluorescence from cultures expressing sfGFP-N150TAG and wt-*Mb*PylRS/PylT in the presence of G-AisoK, AisoK or BocK, or grown without ncAAs. Consistent results were obtained over three distinct replicate experiments. **e.** Extracted ion chromatograms for determining intracellular concentrations of G-AisoK (teal) and AisoK (pink) by an LC-MS assay, performed on K12 cell extracts. Intracellular G-AisoK concentrations in K12 cells grown in the presence of 2 mM G-AisoK are negligible (dark grey, left). Intracellular AisoK concentrations in K12 cells grown in the presence of 2 mM G-AisoK (dark grey, right) are 5-10-fold higher than when grown in the presence of 2 mM AisoK (light grey, right). Consistent results were obtained over three distinct replicate experiments. **f.** Proposed model for increased AisoK incorporation in the presence of G-AisoK. The tripeptide G-AisoK is actively taken up by K12 cells via a transporter. Within the cytoplasm G-AisoK is processed to AisoK, which is a substrate for wt-*Mb*PylRS/PylT and is incorporated site-specifically into a protein of interest (POI).

This observation led us to formulate the hypothesis that a specific transport mechanism may actively pump G-AisoK into cells. Within the cytosol G-AisoK is then enzymatically processed to AisoK, which accumulates in high concentrations and serves as a substrate for wt-PylRS, leading to efficient AisoK encoding (Fig. 1f).

### A bacterial ABC-transporter facilitates tripeptide uptake

In gram-negative organisms such as *E. coli,* small peptidic substrates gain access to the periplasm via diffusion through outer membrane pore-forming proteins known as porins^34^. Depending on the mode of action there are two major classes of active peptide transporters located in the inner membrane that shuffle specific peptides from the periplasm to the cytosol (Fig. S2a): (1) proton-dependent oligopeptide transporters (POTs) that leverage a proton gradient for peptide import and (2) ABC-transporters that use ATP hydrolysis as energy source for peptide import. Bacterial POTs are monomeric, multi-transmembrane (TM) helix-containing proteins that typically import various di- or tripeptides^35^. Bacterial ABC-transporters associated with peptide uptake (e.g. Dpp-, Ddp- and Opp-transporters) require a periplasmic binding protein that delivers the captured peptidic substrate to a multi-subunit transmembrane complex for ATP-dependent promiscuous di- and oligopeptide transport.

In order to identify a potential uptake system for G-AisoK, we used single gene knockouts^36^, which have individual transporter domains deleted, for amber suppression of sfGFP-N150TAG in the presence of the wt-PylRS/PylT pair and G-AisoK. We hypothesized that amber suppression of sfGFP would be abolished or diminished, if a potentially involved transporter system was absent. Deletion of the POT-family members and components of the *ddp* or *dpp* tranporters did not have any influence on efficient sfGFP production (Fig. S2b-c). In contrast, individual knockouts of genes constituting the *opp* operon led to complete abolishment of sfGFP expression in the presence of G-AisoK (Fig. 2a). The Opp transporter shares its general domain organization with other ABC-transporters^37^: it consists of the periplasmic binding protein (OppA), two transmembrane domains (TMDs, OppB and OppC) that span the periplasmic membrane and two cytoplasmic nucleotide-binding domains (NBDs, OppD and OppF, Fig. 2b) that bind and hydrolyze ATP. Docking of peptide-bound OppA to the TMDs triggers ATP binding and dimerization of the NBDs, leading to uptake of the substrate into the translocation channel. Hydrolysis of ATP results in dissociation of the NBD-dimerization interface and flips the TMDs to trigger release of the substrate into the cytosol. ADP-ATP exchange re-induces NBD-dimerization and the TMDs can again bind to OppA (Fig. 2c).

**Figure 2.**
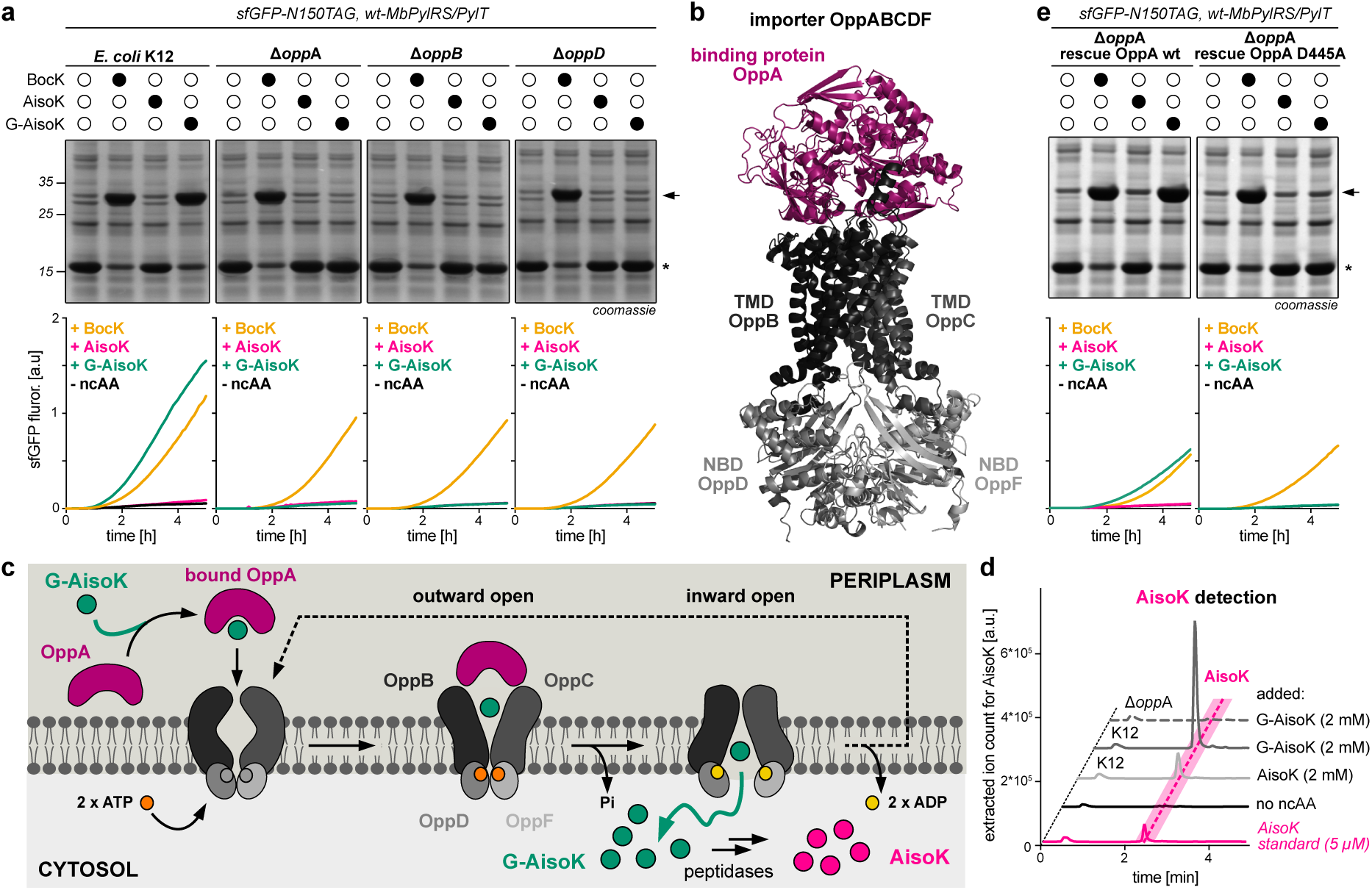
The Opp transporter is responsible for efficient G-AisoK uptake. **a.** SDS-PAGE analysis and time course fluorescence measurements of sfGFP-N150TAG expression in the presence of BocK, AisoK or G-AisoK in K12 cells and in τ1*oppA*, τ1*oppB* or τ1*oppD* knockouts indicating that the Opp transporter is responsible for G-AisoK uptake. Results for other knockouts can be found in Fig. S3a. Consistent results were obtained over three distinct replicate experiments. **b.** AlphaFold2^40^ predicted structure of the Opp transporter consisting of the periplasmic peptide binding protein OppA, two transmembrane domains (TMDs) OppB,C and two nucleotide binding domains (NBDs) OppD,F. **c.** Proposed mechanism of Opp-mediated uptake. G-AisoK binds to OppA in the periplasm and is shuttled to membrane-bound OppB,C. The tripeptide is actively transported into the cytosol in an ATP-dependent manner where it is cleaved by endogenous peptidases to AisoK. OppA in its apo-form is released from the TMDs to allow for binding of new G-AisoK. **d.** Extracted ion chromatograms of *E. coli* K12 lysates for determining intracellular AisoK concentrations in K12 versus τ1*oppA* K12 cells. Genomic deletion of *oppA* results in undetectable AisoK concentrations when growing cells in the presence of 2 mM G-AisoK (grey, dashed). Consistent results were obtained over three distinct replicate experiments. **e.** SDS-PAGE analysis and time course fluorescence measurements of sfGFP-N150TAG expression in Δ*oppA* cells, constitutively expressing OppA variants (left: wt, right: D445A). Plasmid-based constitutive expression of wt-OppA rescues full-length sfGFP expression in presence of G-AisoK, while this is not the case for OppA-D445 expression, indicating that D445 is essential for G-AisoK binding and uptake. BocK dependent sfGFP expression remains unchanged in both cases. Arrow indicates full-length sfGFP, asterisk indicates truncated protein). Consistent results were obtained over three distinct replicate experiments.

Individual deletions of either the binding protein OppA, or any of the two TMDs or NBDs led to complete loss of amber suppression and sfGFP fluorescence in the presence of G-AisoK, while no differences in sfGFP-N150BocK expression yields were observed, indicating that the Opp transporter may be specifically involved in G-AisoK uptake (Fig. 2a, Fig. S3a). Confirming these expression results, uptake assays showed unchanged intracellular BocK concentrations between K12 cells and τ1*oppA*-K12 cells, while AisoK, which accumulated in millimolar concentrations in K12 cells upon addition of G-AisoK, was not detectable when the *oppA* gene was deleted (Fig. 2d, Fig. S3b).

Remarkably, plasmid-based constitutive expression of OppA in τ1o*pp*A-K12 cells completely rescued amber suppression in the presence of G-AisoK (Fig. 2e).

OppA is known to promiscuously bind to 2-5 amino acid long peptides and shows some preference for tri- and tetrapeptides containing positively charged side chains. Previously reported crystal structures of the apo and the liganded form of OppA have elucidated its binding mechanism: OppA shows a three-domain architecture and upon ligand binding, the N- and C-terminal domains rearrange in a Venus fly-trap mechanism to adopt a closed conformation, in which the bound peptide is completely engulfed inside the protein^38^. Furthermore, available structural information of an OppA:tripeptide complex indicates that the OppA binding cavity provides large hydrated pockets for various side chains of the tripeptide (Fig. S3c). Direct interactions between the peptidic substrate and OppA target the backbone and its termini. The N-terminus of the bound tripeptide makes multiple interactions with OppA residues and is kept in place by an extensive network of hydrogen bonds and salt bridges. Specifically, the protonated α-amine of the bound tripeptide forms a salt bridge with OppA residue D445, which is kept in its deprotonated form by hydrogen-bonding to Y135 and H187 (Fig. S3c). The C-terminus of the tripeptide forms a salt bridge with R439. To experimentally verify if these interactions are also necessary for recognition of G-AisoK, we constitutively co-expressed OppA-D445A or OppA-R439A in τ1*oppA*-K12 cells. Expression of the OppA-R439A variant did not impact G-AisoK dependent amber suppression of sfGFP, implying that R439 may not be involved in interacting with the C-terminus of isopeptide-linked G-AisoK (Fig S3a). In stark contrast, overexpression of the OppA-D445A variant did not lead to any sfGFP expression in the presence of G-AisoK, indicating-that the N-terminal amine of G in G-AisoK is interacting with OppA in a similar way as the one of a linear tripeptide (Fig. 2e, Fig. S3d).

To identify a potential peptidase responsible for G-AisoK to AisoK processing within the *E. coli* cytosol, we performed amber suppression experiments in single-gene knockouts^36^ that had individual peptidases or proteases deleted in the presence of G-AisoK. We reasoned that lack of N-terminal glycine processing should lead to diminished protein expression yields. We could however not identify one specific peptidase solely responsible for G-AisoK to AisoK processing (Fig. S4) and speculate that G-AisoK may be a substrate for several promiscuous aminopeptidases^39^.

### A versatile toolbox for efficient encoding of diverse chemical functionalities into POIs

As OppA shows little sequence specificity for bound peptidic substrates, we envisioned to leverage our propeptide strategy for the efficient incorporation of a range of functionalities typically available for GCE, including moieties for site-specific protein conjugation and crosslinking (Fig. 3a). To test the scope of a potential G-XisoK-based toolbox for protein modification, we first synthesized a panel of G-XisoK derivatives bearing natural amino acids serine, threonine, cysteine, valine, leucine, proline or histidine at position X. Supplementing K12 cells with these G-XisoK derivatives, led to efficient suppression of sfGFP-N150TAG and Ub-K63TAG using wt-*Mb*PylRS/PylT or appropriate variants thereof that were identified through screening of a large panel of PylRS-variants available in the lab, with almost no truncation product visible. Importantly, in all cases, MS-analysis confirmed site-specific incorporation of the corresponding XisoK dipeptides and severely reduced protein expression yields were observed when cells were supplemented with XisoK derivatives (Fig. 3b, Fig. S5, S6). Efficient encoding of such XisoK derivatives is interesting, as lysine aminoacylation, in which the ε-amino group of lysine is covalently linked to the α-carboxyl of any of the 20 naturally occurring amino acids, was recently identified as a reversible PTM^41,42^. As previous attempts at directly encoding S/T/P/CisoK derivatives via GCE proved very inefficient^19,41,43–45^ (Fig. 4b), functional studies on these PTMs are elusive.

**Figure 3.**
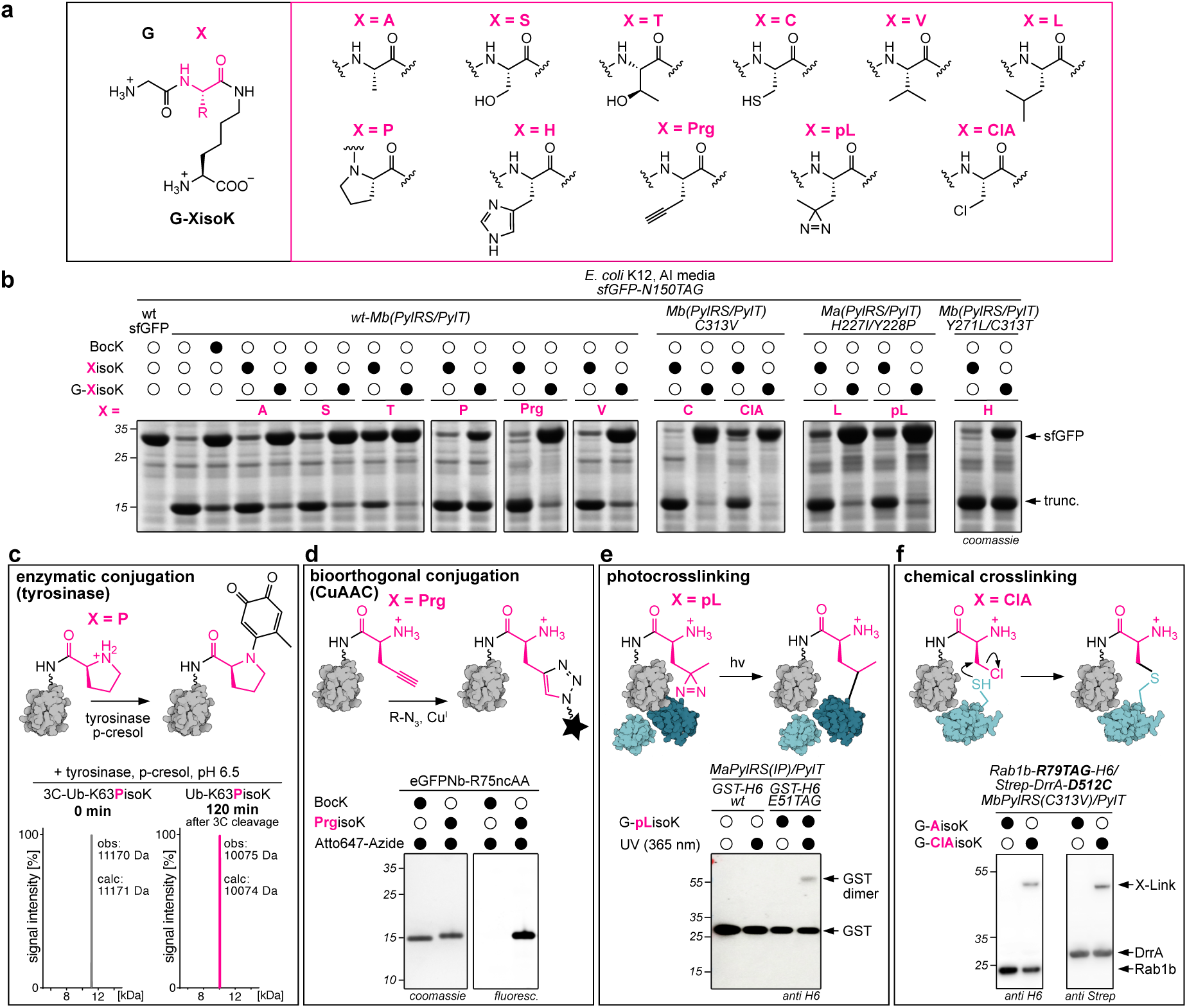
A versatile G-XisoK toolbox. **a.** Structures of all functionalities incorporated into the G-XisoK scaffold. **b.** SDS-PAGE analysis of sfGFP-N150TAG expression in the presence of either 2 mM XisoK or G-XisoK. All G-XisoK derivatives show high levels of full-length sfGFP expression using the corresponding PylRS/PylT pairs. Cells grown in the presence of XisoK derivatives, on the contrary, show much lower or no expression of full-length sfGFP. LC-MS analyses of purified sfGFP and Ub variants, confirming incorporation of XisoK derivatives are shown in Figs. S5, S6 and S9. **c.** LC-MS analysis of tyrosinase-mediated labelling of 3C-Ub bearing PisoK at K63 with p-cresol shows quantitative conversion. Conditions: 20 µM 3C-UbK63PisoK, 100 µM p-cresol, 0.4 µM tyrosinase at pH 6.5, 120 min (for details and full data see Fig. S8) **d.** SDS-PAGE analysis of CuAAC labeling of purified eGFPNb-R75PrgisoK with an Atto647-Azide fluorophore. No labeling is observed for eGFPNb-R75BocK, as expected (for details and full data see Fig. S10a). **e.** Western blot analysis of GST-dimer crosslinking for GST-E51pLisoK upon UV_365 nm_ illumination. No crosslink is observed for GST-wt, as expected (for details and full data see Fig. S10b). **f.** Western blot analysis of proximity-induced chemical crosslinking between Rab1b-R79ClAisoK and its interactor DrrA-D512C_339-522 55_. Cells expressing both binding partners in the presence of G-ClAisoK show a higher molecular weight band corresponding to the crosslinked complex in both anti-H6 and anti-Strep blots. No crosslinking was observed for cells grown in the presence of G-AisoK. Full data and further experiments can be found in Fig. S11. **b-f**) Consistent results were obtained over three distinct replicate experiments.

**Figure 4.**
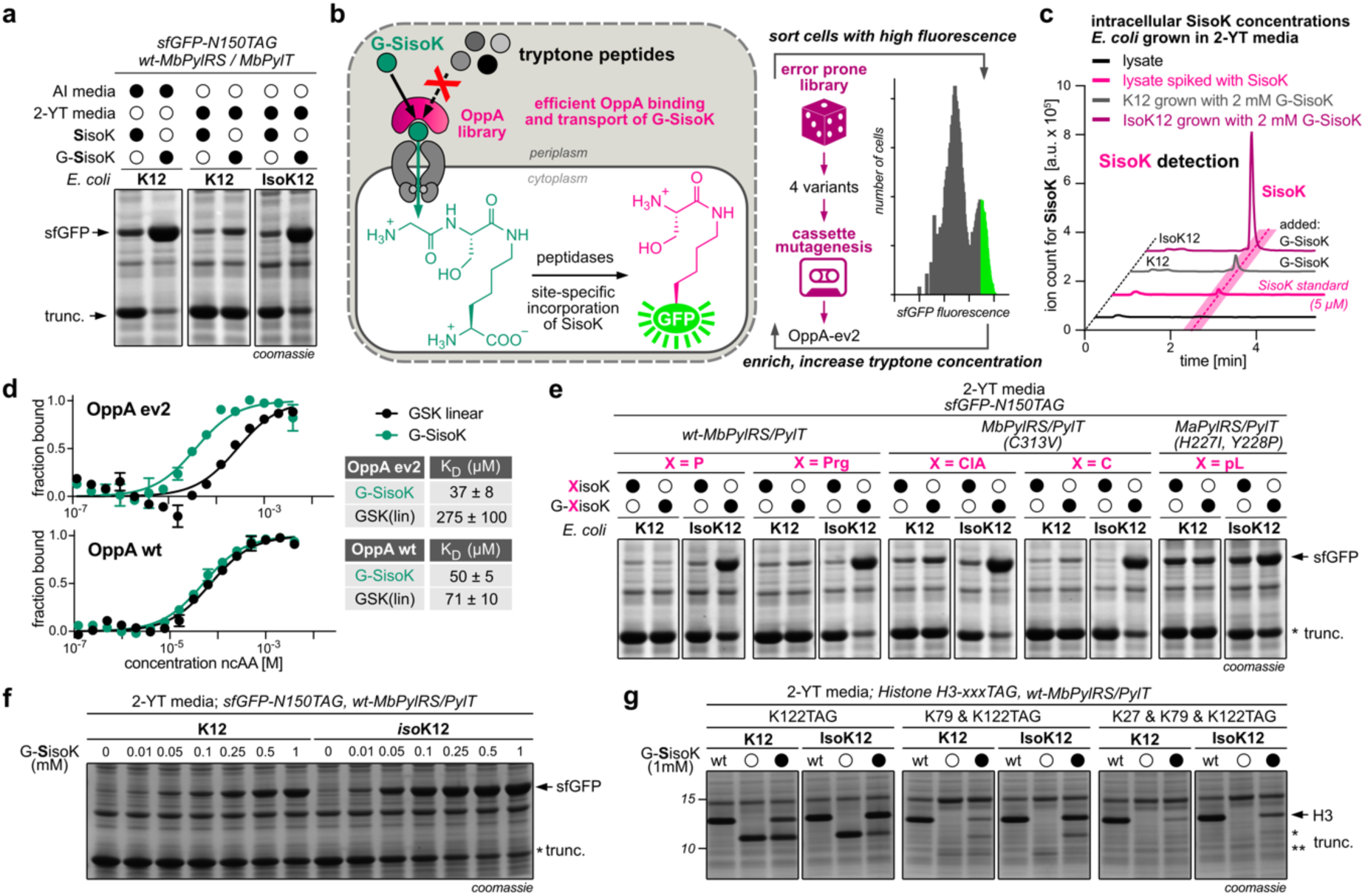
Scalable XisoK incorporation through OppA evolution. **a.** SDS-PAGE analysis of sfGFP-N150TAG expression in the presence of SisoK or G-SisoK in AI media (left) or 2-YT (middle, right). In K12 cells full-length sfGFP expression yields are significantly reduced when using tryptone containing 2-YT (middle) media due to competition between tryptic peptides and G-SisoK for OppA binding. sfGFP expression in 2-YT is recovered when using the engineered IsoK12 strain (right). Consistent results were obtained over three distinct replicate experiments. **b.** Scheme for OppA evolution for better G-SisoK uptake in tryptone containing media. OppA libraries were screened for increased G-SisoK uptake under increasing tryptone concentrations via amber suppression of sfGFP-N150TAG by sorting for fluorescent cells. Initial screening of an error-prone library yielded 4 variants which were used as basis for creating a cassette mutagenesis library, whose screening yielded OppA-ev2. **c.** Extracted ion chromatograms to determine intracellular SisoK concentrations. IsoK12 cells (purple) grown in 2-YT show 7-10-fold higher intracellular SisoK concentrations in comparison to K12 cells (grey) when adding 2 mM G-SisoK to 2-YT media. Consistent results were obtained over three distinct replicate experiments. **d.** Affinity measurements of G-SisoK and a linear GSK peptide (GSK(lin)) towards OppA using microscale thermophoresis. K_D_ values and errors (±s.e.m.) were calculated from three biologically independent experiments (*n* = 3). All data processing was performed using GraphPad Prism 10 (GraphPad software) and MO.affinity Analysis (v3.0.5, NanoTemper Technologies). **e.** SDS-PAGE analysis of sfGFP-N150TAG expression in wt-K12 cells versus IsoK12 cells grown in 2-YT media in presence of G-XisoK derivatives with X = P, Prg, ClA, C, pL. Full-length sfGFP expression is severely increased in IsoK12 cells for all X functionalities. Full gels as well as expression gels for other G-XisoK derivatives can be found in Fig. S12. **f.** SDS-PAGE analysis of sfGFP-N150TAG expression in 2xYT, comparing K12 with IsoK12 cells at different G-SisoK concentrations. **g.** SDS-PAGE analysis of Histone H3 with single (K122TAG), double (K79TAG, K122TAG) and triple (K27TAG, K79TAG, K122TAG) amber suppression in presence of 1 mM G-SisoK comparing K12 with IsoK12 cells grown in 2-YT media. **e-g**) Consistent results were obtained over three distinct replicate experiments.

Furthermore, CisoK-modified proteins are ideally suited for native chemical ligation approaches, as has been shown for generating Ub-conjugates^46,47^. Comparing CisoK-incorporation efficiencies obtained via our propeptide strategy with published CisoK-incorporation yields using specifically evolved PylRS-variants, impressively shows the advantage of actively pumping G-CisoK into bacterial cells^19^ (Fig. S7a). This indicates that intracellular ncAA concentration may be more crucial for efficient ncAA incorporation than finely tuning and evolving PylRS-variants.

Site-specific incorporation of XisoK derivatives, in which X is a natural amino acid, offers also the unique possibility to introduce a specific natural amino acid with an N-terminal α-amine within a POI sequence, equipping a protein with a second, artificial N-terminus. This allows internal protein labeling by applying various bioconjugation strategies that have been developed for targeting specific N-termini.^48^

By efficient encoding of PisoK (via uptake of G-PisoK, Fig. 3b), we introduce proline bearing a free α-amine at an internal site of a target protein and show that this allows tyrosinase-mediated labeling with phenol derivatives.^49^ Tyrosinase oxidizes p-cresol to the corresponding *o*-quinone intermediate that can oxidatively couple to the free α-amine of proline in PisoK. We show specific and quantitative labeling of PisoK-modified Ub, establishing a solid chemoenzymatic labeling approach that is applicable to any user-chosen site in various POIs (Fig. 3c, Fig. S8).

When screening for PylRS-variants for incorporation of XisoK derivatives bearing aromatic side chains at position X, such as in HisoK (Fig. 3a), we could not identify any positive hits upon supplementing cells with G-HisoK. To verify if this stems from poor G-HisoK recognition by OppA or lack of appropriate PylRS-variants for HisoK, we created a custom-designed *Mb*PylRS library with five randomized positions in the active site and subjected this library to alternating rounds of positive and negative selection in K12 cells combined with fluorescence readout in the presence of G-HisoK. Gratifyingly, a novel *Mb*PylRS-variant supported HisoK incorporation in the presence of G-HisoK, but not when supplementing cells with HisoK (Fig. 3b), indicating that OppA also binds and delivers G-XisoK amino acids with bulky and aromatic X-side chains. Efficient incorporation of synthetically easily accessible histidine-containing ncAAs may prove useful for expanding the range of genetically encoded metal coordination environments and may advance the generation and engineering of metalloenzymes with optimized properties and novel activities.^50,51^

Importantly, also non-canonical side chains can be incorporated at position X to further expand the G-XisoK-toolbox for efficient incorporation of diverse functionalities. We explored the uptake of G-XisoK derivatives, where X contained functionalities for bioorthogonal labeling^52^, as well as for light-induced and proximity-induced chemical crosslinking^53,54^. As a considerable advantage of our propeptide strategy, G-XisoK derivatives can be easily synthesized at large scales via SPPS with many non-canonical side chains commercially available as amino acid building blocks. In standard GCE approaches, apart from limiting incorporation yields, the synthetic accessibility of ncAAs often presents a considerable challenge. We first explored incorporation of functionalities that would be amenable for bioorthogonal protein labeling via Cu(I)-catalyzed azide alkyne cycloaddition (CuAAC). A G-XisoK derivative, bearing propargyl-glycine at position X, allowed very efficient installation of PrgisoK into diverse target proteins (Fig. 3b, Fig. S9), including an eGFP-specific nanobody (eGFPNb-R75TAG), which was successfully labeled with a fluorophore-conjugated azide moiety via CuAAC (Fig. 3d, Fig. S10a). As seen for all other tested derivatives, PrgisoK incorporation was successful only when cells were supplemented with G-PrgisoK, but not when supplemented with PrgisoK (Fig. 3b). Similarly, PrgisoK incorporation yields using the corresponding G-bearing tripeptide, compare very favorably to recently reported PrgisoK incorporation efficiencies using a specific PylRS-variant identified for PrgisoK incorporation^41^ (Fig S7b).

To expand the G-XisoK-toolbox to ncAAs useful for mapping and trapping PPIs, we explored efficient incorporation of different crosslinker moieties using our propeptide strategy. We integrated commercially available photoleucine (pL) as residue X in the G-XisoK scaffold. Supplementing K12 cells with this tripeptide, enabled efficient encoding of a diazirine moiety into a POI using an *Methanomethylophilus alvus (Ma)* PylRS-derived variant (Fig. 3b, Fig. S9) and UV-induced crosslinking of PPIs, as exemplified by crosslinking of glutathione-S-transferase (GST) dimer, as well as sfGFP dimer (Fig. 3e, Figure S10b). As pL is commercially available, synthesis of the needed tripeptide via SPPS is much easier, more efficient, and scalable than synthetic access to other diazirine-bearing ncAAs reported for GCE and photo-crosslinking^53–55^, highlighting an important advantage of our strategy.

For trapping transient PPIs in a proximity-induced manner, we designed a G-XisoK scaffold, bearing the finely tuned electrophile chloroalanine (ClA) at position X. Incubation of K12 cells with G-ClAisoK led to efficient incorporation of ClAisoK into different POIs (Fig. 3b, Fig S9). When ClAisoK is placed in proximity to a nucleophilic amino acid in an interacting protein, an S_N_2 nucleophilic reaction can take place, covalently stabilizing the protein complex by a stable thioether bridge (Fig. 3f). By pairwise incorporation of ClAisoK and cysteine residues at protein-protein interfaces, we covalently stabilized a variety of low-affinity protein complexes with K_D_s in the micromolar to low millimolar range in living cells, such as a sfGFP homodimer (Fig. S11a), the interaction between affibody and protein Z^56^ (Fig. S11b) as well as the ternary GDP-bound complex between small G-protein Rab1b and the guanine nucleotide exchange factor (GEF) domain of DrrA^57^ (Fig. 3f, Fig. S11c). Distances of 8-12 Å between the corresponding Cα atoms of cysteine and the lysine of ClAisoK could be efficiently crosslinked (Fig. S11). Apart from cysteine, also histidine and glutamate showed successful covalent homodimer formation with ClAisoK-modified sfGFP (Fig S11a). In vitro crosslinking using purified affibody- and protein-Z-variants confirmed that crosslinking is specific for ClAisoK, while no crosslinking was observed for AisoK-bearing affibody, as expected (Fig. S11b).

### Scalable and cheap XisoK incorporation through OppA evolution

For all tested G-XisoK tripeptides, we observed highly efficient protein production in chemically-defined auto induction (AI) media^58^. Amber suppression in nutrient-rich media, such as lysogeny broth (LB) or 2-YT, did, however, not give the desired yields of protein expression (Fig. 4a, Fig. S12a-b). We hypothesized that G-XisoK tripeptides may compete for OppA-binding with short linear peptides that are abundant in nutrient-rich expression media based on peptone or tryptone. This was confirmed by uptake assays determining intracellular SisoK concentrations, which upon G-SisoK supplementation, were sixfold lower in 2-YT compared to AI media, indicating that OppA-mediated active transport is impaired in such conditions (Fig. S13). This drastically impacts the implementation of our versatile G-XisoK toolbox for cheap and scalable protein production, as chemically-defined AI media is much more expensive and cumbersome to prepare than regular nutrient-rich growth media and does not support as high cell-density cultures leading to decreased biomass and lower overall expression yields of ncAA-modified proteins.

We therefore aimed at evolving an engineered OppA-variant that preferentially binds G-SisoK over short linear peptides as present in peptone or tryptone-based media. We developed a fluorescence-activated cell sorting (FACS)-based screening platform to alter the binding preference and selectivity of OppA using directed evolution. Our screening system couples uptake of the G-SisoK tripeptide to sfGFP fluorescence by suppressing the amber codon in sfGFP-N150TAG (Fig. 4b). We co-transformed an error-prone OppA-library together with the wt-*Mb*PylRS/PylT pair and sfGFP-N150TAG into τ1*oppA*-K12 cells in AI media and subjected these cells to subsequent rounds of enrichment from low (1g/L) to high (16g/L) tryptone concentrations. We identified four converging OppA-variants each harboring 4-5 mutations distributed all over the OppA-fold (Fig. S14). Mutations that were present in more than one variant or were structurally close to each other, were identified as hotspot positions and selected for saturation mutagenesis on all four OppA-variants and wt-OppA. The obtained library was again subjected to multiple FACS-based enrichment steps in LB and 2-YT media. The final selected OppA-variant (OppA-ev2) contains a total of seven mutations, distributed over the entire OppA-fold, with only one mutation (R439Q) being closer than 4 Å to the binding site (Fig. S14). We introduced these mutations into the K12 genome via lambda red-mediated homologous recombination to create a K12-derived engineered *E. coli* strain for scalable and cheap production of XisoK-bearing proteins in peptide containing media, which we dubbed IsoK12. Doubling times for IsoK12 and its parent strain were comparable both in AI and 2-YT media (Fig. S15a). With the IsoK12 strain in hand, we first measured intracellular SisoK concentrations in presence of G-SisoK in 2-YT media. Gratifyingly, intracellular SisoK concentrations substantially increased (7-10-fold) when compared to K12 cells (Fig. 4c), while BocK concentrations did not differ between the two *E. coli* strains, indicating that OppA-ev2 indeed preferentially binds and transports G-SisoK over linear peptides present in 2-YT (Fig. 4c, Fig. S13). Importantly, the observed elevated SisoK concentrations in IsoK12 cells led to efficient amber suppression of TAG-containing constructs in 2-YT media, exhibiting similar efficiencies as observed for K12 cells grown in defined AI media (Fig. 4a, Fig. S12c).

To confirm OppA-ev2 binding preference, we measured K_D_s of wt-OppA and OppA-ev2 towards G-SisoK, a linear GSK tripeptide (mimicking linear tryptone peptides) and SisoK via microscale thermophoresis (Fig. 4d, Fig. S15b). While both wt-OppA and OppA-ev2 showed similarly low binding affinities towards SisoK (∼300 µM), affinity of G-SisoK towards OppA-ev2 is slightly increased compared to wt-OppA (37 µM versus 50 µM). Interestingly, binding of linear GSK was four-fold decreased for OppA-ev2 versus wt-OppA (275 µM versus 71 µM), corroborating the *in cellulo* data that G-SisoK is able to compete with linear tryptone-derived peptides in the presence of OppA-ev2. In an attempt to structurally rationalize this binding behavior, we mapped the seven mutations found in OppA-ev2 onto a published OppA structure liganded to a linear tripeptide (Fig.S3c, Fig. S14, PDB ID: 3TCF)^38^. Most of the mutations are not directly contacting the bound tripeptide, but interestingly residue R439 that is important for keeping the C-terminus of the linear tripeptide in place is mutated to glutamine in OppA-ev2. The R439Q mutation may therefore be responsible for the observed lowered affinity for linear GSK, while the binding affinity of G-SisoK, which does not display a carboxy group at this position, is not affected by this mutation.

As the OppA-ev2-variant was evolved in the presence of G-SisoK, we wondered if preferential binding and uptake was extendable to other G-XisoK derivatives. Indeed, all of the tested G-XisoK derivatives (X=A, S, T, C, V, L, P, H, Prg, pL and ClA) led to very efficient amber suppression of sfGFP-N150TAG, with hardly any amounts of truncated side-product when expressed in IsoK12 cells grown in 2-YT media, resembling protein yields obtained in K12 cells grown in chemically-defined and peptide-free AI media (Fig. 4e, Fig. S12c). In fact, tripeptide uptake was so efficient in IsoK12 cells that G-SisoK concentrations as low as 50-100 µM led to similar incorporation yields as observed in K12 cells, supplemented with 1 mM G-SisoK, thereby decreasing necessary ncAA concentrations by 10-fold (Fig. 4f). We show that our propeptide strategy combined with engineered IsoK12 enables high-yielding XisoK-incorporation at various positions in diverse target proteins spanning sizes from 7 to 85 kDa. Amber suppression of PCNA, ß-lactamase, SUMO2, Calmodulin, eGFPNb, and Hsp82 attest to the wide applicability of the G-XisoK/IsoK12 combination resulting in highly efficient production of XisoK-modified proteins matching or even exceeding wt expression yields. (Fig. S16-17). Interestingly, in many cases G-XisoK uptake was also improved when IsoK12 cells were grown in AI media (Fig. S13b, S16-17). Preparative large-scale production and purification of a GFP nanobody with an ncAA bearing a propargyl moiety showed that the G-PrgisoK/IsoK12 combination resulted in similar purified protein yields (44 mg/L) as obtained for wt expression (41 mg/L) and exceeded yields from a previously reported optimized alkyne-bearing ncAA/PylRS combination (Fig. S17b). Increasing intracellular ncAA concentrations via efficient tripeptide uptake facilitated also multi-site amber suppression within one target protein. We introduced TAG-codons at up to three positions into Histone H3 (K27, K79, K122) and expressed the corresponding variants in the presence of G-SisoK. IsoK12 outperformed K12 in single, double and triple amber suppression with expression yields rivaling wt-H3 production, while only minute amounts of doubly- and triply-suppressed protein were obtained with the gold-standard ncAA BocK (Fig. 4g, Fig. S18).

### Concomitant uptake strategy for efficient encoding of two different ncAAs

Efficient G-XisoK uptake and its processing leads to high intracellular concentrations not only of dipeptide XisoK, but also of the N-terminal glycine that is cleaved off. As OppA is quite promiscuous in regard to bound peptide sequences, we hypothesized that also other side chains, including those of ncAAs, might be accepted at this position and corresponding non-canonical tripeptides might be efficiently transported into the *E. coli* cytosol. If ncAAs at this N-terminal position were also efficiently cleaved by peptidases, this could present a general tool to increase the intracellular concentration of two encodable ncAAs with one single, easily synthesizable tripeptide. We first synthesized a K-SisoK tripeptide, with lysine as N-terminal amino acid instead of glycine (Fig. S19a). Interestingly, supplementation of K12 or IsoK12 cells with K-SisoK led to similarly efficient amber suppression of sfGFP-N150TAG in the presence of wt-*Mb*PylRS/PylT as observed for G-SisoK (Fig. S19b). LC-MS analysis of purified sfGFP confirmed incorporation of SisoK (Fig. S19c), proving that indeed K-SisoK must be efficiently transported into cells and N-terminally processed within the *E. coli* cytosol to K and SisoK. In order to leverage such a concomitant uptake strategy for the site-specific incorporation of two different ncAAs into a single protein, we designed a tripeptide that contains a PTM-bearing ncAA and a photocrosslinker ncAA. Dual site-specific incorporation of such moieties into the same POI represents an ideal tool to investigate PTM-specific protein interactors or to chemically stabilize transient POI-reader complexes^59^. pLisoK was shown to be incorporated successfully into target proteins using a *Ma*PylRS-variant and supplementing cells with G-pLisoK (Fig. 3b). Acetyl-lysine (AcK), an abundant PTM of both prokaryotic and eukaryotic proteins can be specifically incorporated into target proteins by using the previously published *Mb*PylRS-variant AcKRS3^60^. Importantly, the two PylRS-variants are orthogonal to each other^61^ with AcK not being a substrate for the *Ma*PylRS-variant and vice-versa (Fig. S19d). We synthesized the tripeptide AcK-pLisoK (Fig. 5a) and showed its efficient uptake and cleavage leading to high intracellular concentrations of AcK and pLisoK, which allowed efficient dual suppression of a TAA codon by AcK and a TAG codon by pLisoK within the same target protein (sfGFP-N40Ack-N150pLisoK, Fig. 5b) using the corresponding orthogonal PylRS/PylT pairs. Importantly, significantly higher protein yields were achieved by supplementing cells with 0.5 mM of AcK-pLisoK, rather than adding 0.5 mM of both AcK and G-pLisoK to the medium, attesting to efficient uptake and cleavage (Fig 5b). LC-MS analysis of full-length sfGFP confirmed dual incorporation of AcK and pLisoK (Fig. 5c). To confirm true mutual orthogonality and absence of misincorporation of the two ncAAs and their respective PylRS-variants, we designed an Ub-SUMO2 fusion construct with a TEV protease recognition sequence linking the two target proteins (Ub-K48TAG-TEV-SUMO-K11TAA, Fig. 5d). Expression, purification and subsequent TEV cleavage of the construct, followed by LC-MS analysis identified specific incorporation of pLisoK according to the TAG codon at position 48 of Ub and encoding of AcK in response to the TAA codon at position 11 of SUMO2 (Fig. 5d).

**Figure 5.**
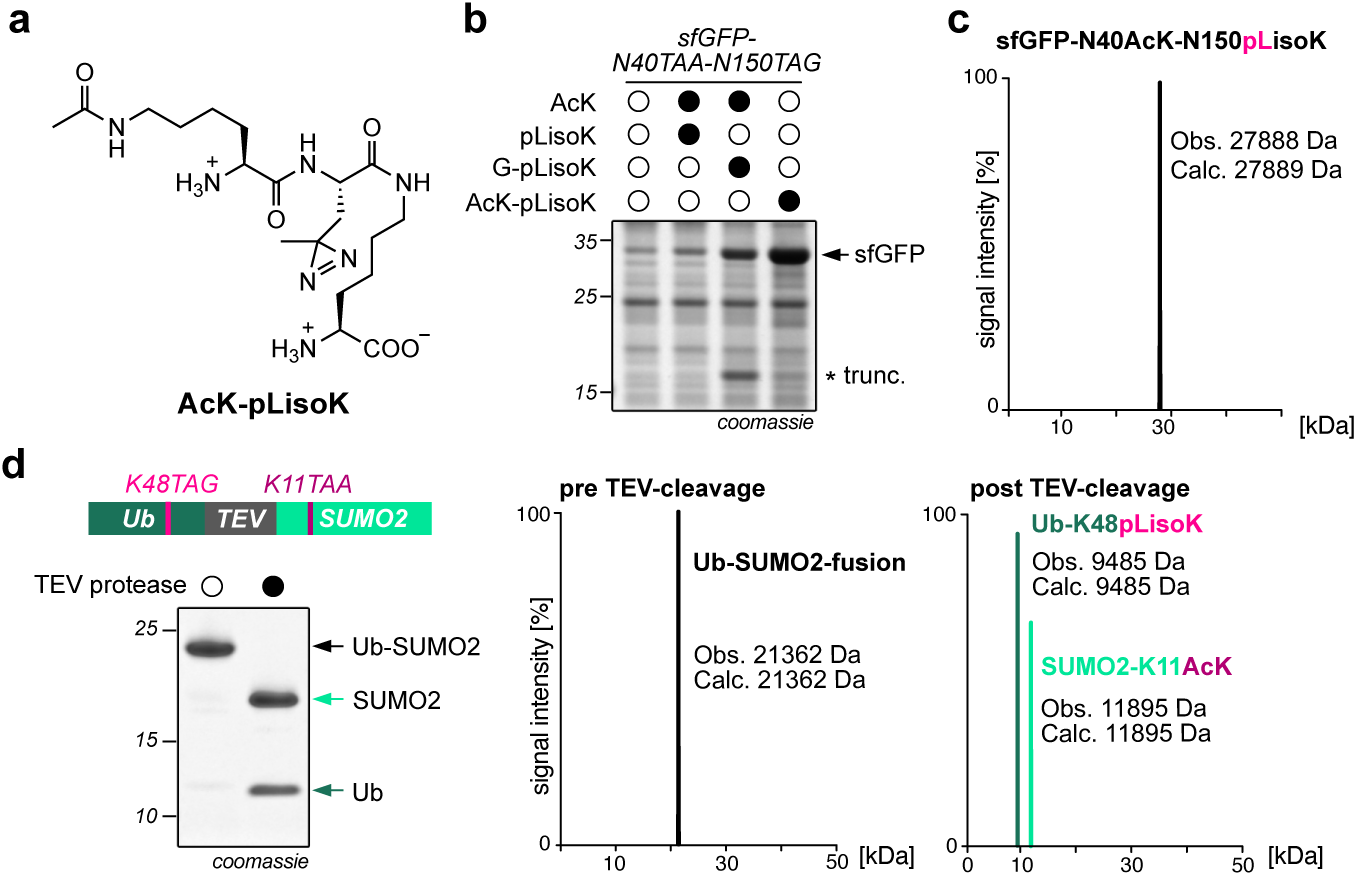
Dual stop codon suppression using a single isopeptide-linked tripeptide. **a.** Chemical structure of the tripeptide AcK-pLisoK. **b.** SDS-PAGE analysis of sfGFP-N40TAA-N150TAG expression in the presence of either AcK and G-pLisoK (or pLisoK) separately added to media or in the presence of tripeptide AcK-pLisoK in IsoK12 cells grown in AI media. Consistent results were obtained over three distinct replicate experiments. **c.** LC-MS analysis of purified sfGFP confirms dual ncAA incorporation (AcK and pLisoK) upon addition of tripeptide AcK-pLisoK. **d.** Left: Scheme of Ub-SUMO fusion construct used to confirm dual and orthogonal incorporation of pLisoK and AcK. SDS-PAGE analysis shows purified full-length construct and products of TEV protease cleavage. Right: LC-MS analysis of full-length construct and cleaved products confirming pLisoK and AcK incorporation at the intended positions.

## Conclusion and discussion

Low protein production yields combined with difficult chemical access to ncAAs are long-standing and critical challenges in genetic code expansion that have hampered widespread and straightforward implementation of such approaches for generating proteins with potential therapeutic and biotechnological significance. In this study we successfully demonstrate the potential of leveraging a propeptide strategy together with the engineering of a bacterial membrane transporter to enhance intracellular concentrations of ncAAs and thereby boost expression yields of proteins bearing site-specifically incorporated ncAAs to rival those of wt-proteins. Isopeptide-linked G-XisoK tripeptides serve as trojan horses that are easily synthesized from commercially available building blocks via SPPS, and are privileged ligands for the periplasmic binding protein OppA, enabling active, ATP-driven transport into the cytosol. Once inside the cell, the N-terminal glycine is enzymatically processed, leading to intracellular accumulation of isopeptide-linked XisoK ncAAs that serve as excellent substrates for orthogonal PylRS/PylT pairs. We show very efficient encoding of eleven different XisoK ncAAs, seven of which have not been incorporated into proteins before, greatly enhancing the chemical space of recombinantly generated proteins via GCE. Among these functionalities are ncAAs bearing bioorthogonal handles, novel PTMs, chemical crosslinker and photocrosslinker moieties, as well as functionalities for chemoenzymatic ligations and we show proof-of-principle applications for all examples. XisoK ncAAs have positively charged side chains under physiological conditions, making them ideal moieties for applications like protein labeling, where site-specific incorporation at surface-exposed positions is required. This avoids aggregation and misfolding associated with large hydrophobic ncAAs, as typically used for labeling purposes^62,63^. With these advantages, we foresee that the XisoK-toolbox can be further developed through aaRS engineering to accommodate functionalities not addressed here such as moieties for inverse-electron demand Diels-Alder cycloadditions or spectroscopic probes. To make our approach amenable to protein production in nutrient-rich, peptide-containing expression media and to evolve an OppA variant that preferentially shuffles isopeptide-linked tripeptides over linear tripeptides into the *E. coli* cytosol, we have developed a FACS-based evolution platform for screening OppA libraries. The high-throughput evolution of periplasmic binding proteins holds untapped potential for expanding the scope of ncAA-peptide substrates for which transport could be engineered. We envision that this may lead to the development of a generalizable system for ncAA transport, allowing for the design and incorporation of ncAAs without the need to design for passive membrane permeability. Such advancements will further expand the chemical space available for genetic code expansion.

Engineering transport systems could facilitate the discovery and evolution of orthogonal aaRS enzymes for charging and encoding ncAAs. This is exemplified by XisoK ncAAs, where previous attempts at incorporation required extensive screening or evolution for moderate efficiencies. In contrast, with G-XisoK-based uptake, efficient incorporation for a variety of new ncAAs was achieved with minimal aaRS engineering. We anticipate that screening and evolving orthogonal aaRS enzymes under increased ncAA regimes as obtainable via active uptake may yield even more successful outcomes.

Beyond increasing incorporation efficiencies, active uptake offers an economical strategy for producing ncAA-bearing proteins. With active transporters such as Opp, significantly lower concentrations of ncAA can be used in media without compromising on protein yields. We envision that Opp-based active uptake can work synergistically with other efforts to improve GCE, including the use of release factor deficient strains^13,14^, or genomically recoded *E. coli* for incorporating multiple ncAAs^16,17^. Excitingly, we have shown that Opp-based uptake and enzymatic processing of corresponding isopeptide-linked tripeptides can be adapted for dual ncAA incorporation and could be expanded further for multi ncAA incorporation in combination with discovery and evolution of corresponding uptake systems and tailoring enzymes.

Adapting our propeptide strategy combined with discovery and engineering of transport systems in other model organisms/cell lines such as yeast or mammalian cells may prove a game changer for scalable expression of ncAA-containing proteins that cannot be expressed in *E. coli*, allowing efficient generation of homogenously modified antibody-drug conjugates.

## Acknowledgments

This work was supported by funding from ETH Zurich and the European Research Council (ERC under the European Union’s Horizon 2020 research and innovation program, grant agreement no. 101003289─Ubl-tool to K.L.). We also thank Lang group members for useful discussions and input.

## References

1 Icking, L. S. et al. iNClusive: a database collecting useful information on non-canonical amino acids and their incorporation into proteins for easier genetic code expansion implementation. Nucleic Acids Res 52, D476–D482 (2024). 10.1093/nar/gkad1090

2 Yi, H. B. et al. Cellular and Biophysical Applications of Genetic Code Expansion. Chemical Reviews 124, 7465–7530 (2024). 10.1021/acs.chemrev.4c00112

3 Koch, N. G. & Budisa, N. Evolution of Pyrrolysyl-tRNA Synthetase: From Methanogenesis to Genetic Code Expansion. Chemical Reviews (2024). 10.1021/acs.chemrev.4c00031

4 Umehara, T. et al. N-acetyl lysyl-tRNA synthetases evolved by a CcdB-based selection possess N-acetyl lysine specificity in vitro and in vivo. FEBS Lett 586, 729–733 (2012). 10.1016/j.febslet.2012.01.029

5 O’Donoghue, P., Ling, J., Wang, Y. S. & Soll, D. Upgrading protein synthesis for synthetic biology. Nat Chem Biol 9, 594–598 (2013). 10.1038/nchembio.1339

6 Young, T. S., Ahmad, I., Yin, J. A. & Schultz, P. G. An enhanced system for unnatural amino acid mutagenesis in E. coli. J Mol Biol 395, 361–374 (2010). 10.1016/j.jmb.2009.10.030

7 Ryu, Y. & Schultz, P. G. Efficient incorporation of unnatural amino acids into proteins in Escherichia coli. Nat Methods 3, 263–265 (2006). 10.1038/nmeth864

8 Jewel, D. et al. Virus-assisted directed evolution of enhanced suppressor tRNAs in mammalian cells. Nat Methods 20, 95–103 (2023). 10.1038/s41592-022-01706-w

9 Guo, J., Melancon, C. E., 3rd, Lee, H. S., Groff, D. & Schultz, P. G. Evolution of amber suppressor tRNAs for efficient bacterial production of proteins containing nonnatural amino acids. Angew Chem Int Ed Engl 48, 9148–9151 (2009). 10.1002/anie.200904035

10 Bryson, D. I. et al. Continuous directed evolution of aminoacyl-tRNA synthetases. Nat Chem Biol 13, 1253–1260 (2017). 10.1038/nchembio.2474

11 Wang, K., Neumann, H., Peak-Chew, S. Y. & Chin, J. W. Evolved orthogonal ribosomes enhance the efficiency of synthetic genetic code expansion. Nat Biotechnol 25, 770–777 (2007). 10.1038/nbt1314

12 Neumann, H., Wang, K., Davis, L., Garcia-Alai, M. & Chin, J. W. Encoding multiple unnatural amino acids via evolution of a quadruplet-decoding ribosome. Nature 464, 441–444 (2010). 10.1038/nature08817

13 Mukai, T. et al. Highly reproductive Escherichia coli cells with no specific assignment to the UAG codon. Sci Rep 5, 9699 (2015). 10.1038/srep09699

14 Johnson, D. B. et al. RF1 knockout allows ribosomal incorporation of unnatural amino acids at multiple sites. Nat Chem Biol 7, 779–786 (2011). 10.1038/nchembio.657

15 Lajoie, M. J. et al. Genomically recoded organisms expand biological functions. Science 342, 357–360 (2013). 10.1126/science.1241459

16 Fredens, J. et al. Total synthesis of Escherichia coli with a recoded genome. Nature 569, 514–518 (2019). 10.1038/s41586-019-1192-5

17 Robertson, W. E. et al. Sense codon reassignment enables viral resistance and encoded polymer synthesis. Science 372, 1057–1062 (2021). 10.1126/science.abg3029

18 Zhang, M. S. et al. Biosynthesis and genetic encoding of phosphothreonine through parallel selection and deep sequencing. Nat Methods 14, 729–736 (2017). 10.1038/nmeth.4302

19 Tai, J. et al. Pyrrolysine-Inspired in Cellulo Synthesis of an Unnatural Amino Acid for Facile Macrocyclization of Proteins. J Am Chem Soc 145, 10249–10258 (2023). 10.1021/jacs.3c01291

20 Marchand, J. A. et al. Discovery of a pathway for terminal-alkyne amino acid biosynthesis. Nature 567, 420–424 (2019). 10.1038/s41586-019-1020-y

21 Chen, Y. et al. Unleashing the potential of noncanonical amino acid biosynthesis to create cells with precision tyrosine sulfation. Nat Commun 13, 5434 (2022). 10.1038/s41467-022-33111-4

22 Mehl, R. A. et al. Generation of a bacterium with a 21 amino acid genetic code. J Am Chem Soc 125, 935–939 (2003). 10.1021/ja0284153

23 Zhu, P. et al. PermaPhos (Ser) : autonomous synthesis of functional, permanently phosphorylated proteins. bioRxiv (2021). 10.1101/2021.10.22.465468

24 Ko, W., Kumar, R., Kim, S. & Lee, H. S. Construction of Bacterial Cells with an Active Transport System for Unnatural Amino Acids. ACS Synth Biol 8, 1195–1203 (2019). 10.1021/acssynbio.9b00076

25 Rodríguez-Robles, E. et al. Rational design of a bacterial import system for new-to-nature molecules. Metabolic Engineering 85, 26–34 (2024). 10.1016/j.ymben.2024.05.005

26 Fickel, T. E. & Gilvarg, C. Transport of impermeant substances in E. coli by way of oligopeptide permease. Nat New Biol 241, 161–163 (1973). 10.1038/newbio241161a0

27 Ames, B. N., Ames, G. F., Young, J. D., Tsuchiya, D. & Lecocq, J. Illicit transport: the oligopeptide permease. Proc Natl Acad Sci U S A 70, 456–458 (1973). 10.1073/pnas.70.2.456

28 Luo, X. et al. Genetically encoding phosphotyrosine and its nonhydrolyzable analog in bacteria. Nat Chem Biol 13, 845–849 (2017). 10.1038/nchembio.2405

29 Moussatova, A., Kandt, C., O’Mara, M. L. & Tieleman, D. P. ATP-binding cassette transporters in Escherichia coli. Biochim Biophys Acta 1778, 1757–1771 (2008). 10.1016/j.bbamem.2008.06.009

30 Fottner, M. et al. Site-specific ubiquitylation and SUMOylation using genetic-code expansion and sortase. Nat Chem Biol 15, 276–284 (2019). 10.1038/s41589-019-0227-4

31 Fottner, M. et al. Site-Specific Protein Labeling and Generation of Defined Ubiquitin-Protein Conjugates Using an Asparaginyl Endopeptidase. Journal of the American Chemical Society 144, 13118–13126 (2022). 10.1021/jacs.2c02191

32 Fottner, M. et al. A modular toolbox to generate complex polymeric ubiquitin architectures using orthogonal sortase enzymes. Nat Commun 12, 6515 (2021). 10.1038/s41467-021-26812-9

33 Huguenin-Dezot, N. et al. Trapping biosynthetic acyl-enzyme intermediates with encoded 2,3-diaminopropionic acid. Nature 565, 112–117 (2019). 10.1038/s41586-018-0781-z

34 Vergalli, J. et al. Porins and small-molecule translocation across the outer membrane of Gram-negative bacteria. Nat Rev Microbiol 18, 164–176 (2020). 10.1038/s41579-019-0294-2

35 Davidson, A. L. & Chen, J. ATP-binding cassette transporters in bacteria. Annu Rev Biochem 73, 241–268 (2004). 10.1146/annurev.biochem.73.011303.073626

36 Baba, T. et al. Construction of Escherichia coli K-12 in-frame, single-gene knockout mutants: the Keio collection. Mol Syst Biol 2, 20060008 (2006). 10.1038/msb4100050

37 Thomas, C. & Tampe, R. Structural and Mechanistic Principles of ABC Transporters. Annu Rev Biochem 89, 605–636 (2020). 10.1146/annurev-biochem-011520-105201

38 Klepsch, M. M. et al. Escherichia coli peptide binding protein OppA has a preference for positively charged peptides. J Mol Biol 414, 75–85 (2011). 10.1016/j.jmb.2011.09.043

39 Miller, C. G. Peptidases and proteases of Escherichia coli and Salmonella typhimurium. Annu Rev Microbiol 29, 485–504 (1975). 10.1146/annurev.mi.29.100175.002413

40 Jumper, J. et al. Highly accurate protein structure prediction with AlphaFold. Nature 596, 583–589 (2021). 10.1038/s41586-021-03819-2

41 Zang, J. et al. Genetic code expansion reveals aminoacylated lysine ubiquitination mediated by UBE2W. Nat Struct Mol Biol 30, 62–71 (2023). 10.1038/s41594-022-00866-9

42 He, X. D. et al. Sensing and Transmitting Intracellular Amino Acid Signals through Reversible Lysine Aminoacylations. Cell Metab 27, 151–166 e156 (2018). 10.1016/j.cmet.2017.10.015

43 Gran-Scheuch, A., Bonandi, E. & Drienovská, I. Expanding the Genetic Code: Incorporation of Functional Secondary Amines via Stop Codon Suppression. ChemCatChem 16, e202301004 (2024). 10.1002/cctc.202301004

44 Brabham, R. L. et al. Rapid sodium periodate cleavage of an unnatural amino acid enables unmasking of a highly reactive alpha-oxo aldehyde for protein bioconjugation. Org Biomol Chem 18, 4000–4003 (2020). 10.1039/d0ob00972e

45 Lee, M. M. et al. Pyrrolysine-inspired protein cyclization. Chembiochem 15, 1769–1772 (2014). 10.1002/cbic.201402129

46 Li, X., Fekner, T., Ottesen, J. J. & Chan, M. K. A pyrrolysine analogue for site-specific protein ubiquitination. Angew Chem Int Ed Engl 48, 9184–9187 (2009). 10.1002/anie.200904472

47 Zhou, H. et al. Linkage-Specific Synthesis of Di-ubiquitin Probes Enabled by the Incorporation of Unnatural Amino Acid ThzK. Chembiochem 23, e202200133 (2022). 10.1002/cbic.202200133

48 Rosen, C. B. & Francis, M. B. Targeting the N terminus for site-selective protein modification. Nat Chem Biol 13, 697–705 (2017). 10.1038/nchembio.2416

49 Maza, J. C. et al. Enzymatic Modification of N-Terminal Proline Residues Using Phenol Derivatives. J Am Chem Soc 141, 3885–3892 (2019). 10.1021/jacs.8b10845

50 Green, A. P., Hayashi, T., Mittl, P. R. E. & Hilvert, D. A Chemically Programmed Proximal Ligand Enhances the Catalytic Properties of a Heme Enzyme. Journal of the American Chemical Society 138, 11344–11352 (2016). 10.1021/jacs.6b07029

51 Huang, H. et al. Genetically encoded Ndelta-vinyl histidine for the evolution of enzyme catalytic center. Nat Commun 15, 5714 (2024). 10.1038/s41467-024-50005-9

52 Lang, K. & Chin, J. W. Cellular incorporation of unnatural amino acids and bioorthogonal labeling of proteins. Chem Rev 114, 4764–4806 (2014). 10.1021/cr400355w

53 Chin, J. W., Martin, A. B., King, D. S., Wang, L. & Schultz, P. G. Addition of a photocrosslinking amino acid to the genetic code of Escherichiacoli. Proc Natl Acad Sci U S A 99, 11020–11024 (2002). 10.1073/pnas.172226299

54 Nguyen, T. A., Cigler, M. & Lang, K. Expanding the Genetic Code to Study Protein-Protein Interactions. Angew Chem Int Ed Engl 57, 14350–14361 (2018). 10.1002/anie.201805869

55 Cigler, M. et al. Proximity-Triggered Covalent Stabilization of Low-Affinity Protein Complexes In Vitro and In Vivo. Angew Chem Int Ed Engl 56, 15737–15741 (2017). 10.1002/anie.201706927

56 Hogbom, M., Eklund, M., Nygren, P. A. & Nordlund, P. Structural basis for recognition by an in vitro evolved affibody. Proc Natl Acad Sci U S A 100, 3191–3196 (2003). 10.1073/pnas.0436100100

57 Du, J. et al. Rab1-AMPylation by Legionella DrrA is allosterically activated by Rab1. Nat Commun 12, 460 (2021). 10.1038/s41467-020-20702-2

58 Muzika, M. et al. Chemically-defined lactose-based autoinduction medium for site-specific incorporation of non-canonical amino acids into proteins. RSC Adv 8, 25558–25567 (2018). 10.1039/c8ra04359k

59 Zheng, Y., Gilgenast, M. J., Hauc, S. & Chatterjee, A. Capturing Post-Translational Modification-Triggered Protein-Protein Interactions Using Dual Noncanonical Amino Acid Mutagenesis. ACS Chem Biol 13, 1137–1141 (2018). 10.1021/acschembio.8b00021

60 Neumann, H. et al. A method for genetically installing site-specific acetylation in recombinant histones defines the effects of H3 K56 acetylation. Mol Cell 36, 153–163 (2009). 10.1016/j.molcel.2009.07.027

61 Willis, J. C. W. & Chin, J. W. Mutually orthogonal pyrrolysyl-tRNA synthetase/tRNA pairs. Nat Chem 10, 831–837 (2018). 10.1038/s41557-018-0052-5

62 Lang, K. et al. Genetic Encoding of bicyclononynes and trans-cyclooctenes for site-specific protein labeling in vitro and in live mammalian cells via rapid fluorogenic Diels-Alder reactions. J Am Chem Soc 134, 10317–10320 (2012). 10.1021/ja302832g

63 Jang, H. S., Jana, S., Blizzard, R. J., Meeuwsen, J. C. & Mehl, R. A. Access to Faster Eukaryotic Cell Labeling with Encoded Tetrazine Amino Acids. J Am Chem Soc 142, 7245–7249 (2020). 10.1021/jacs.9b11520

