## Supplementary Information for "Hijacking a bacterial membrane transporter for efficient genetic code expansion"

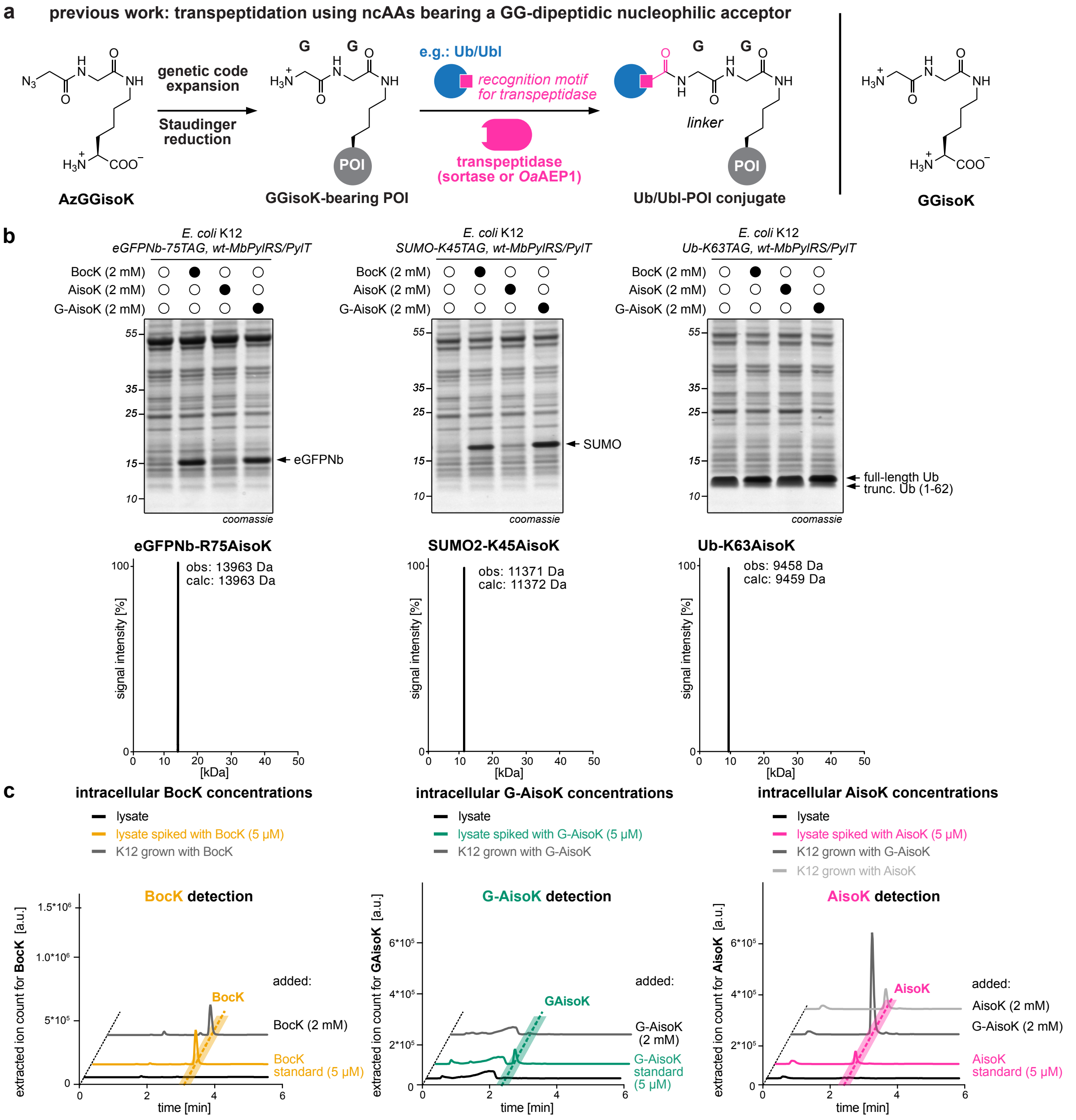

**Figure S1:** Previous work and genetic encoding of AisoK in the presence of G-AisoK into different POIs. **a.** Previously reported ncAA AzGGisoK (left) is site-specifically incorporated into proteins via GCE and decaged via Staudinger reduction to yield GGisoK-modified POIs. The dipeptidic GG acceptor nucleophile can subsequently engage in transpeptidation with proteins bearing a recognition motif at their C-termini using sortase or *Oa*AEP1 to create protein-protein conjugates. Direct incorporation of GGisoK was not successful (right). **b**. Top: SDS-PAGE analysis of eGFP nanobody (eGFPNb-R75TAG), SUMO2-K45TAG and Ubiquitin (Ub-K63TAG) expression using wt-*Mb*PylRS/PylT in the presence of BocK, AisoK and G-AisoK. Very efficient full length protein expression when cells were supplemented with G-AisoK (comparable to BocK-incorporation). Bottom: LC-MS analysis of all three proteins purified from cells supplemented with G-AisoK revealed specific AisoK incorporation. **c**. LC-MS traces of *E. coli* lysates in selected-ion mode to measure intracellular concentrations of BocK (left), G-AisoK (middle) and AisoK (right). Cells grown in the presence of 2 mM BocK showed low (~0.3 mM) intracellular BocK concentrations. Cells grown in presence of 2 mM G-AisoK showed no detectable intracellular G-AisoK (middle). In contrast, intracellular AisoK concentrations were up to 10-fold higher (right, dark grey) when cells were supplemented with G-AisoK as compared to cells grown in 2 mM AisoK (right, light grey).

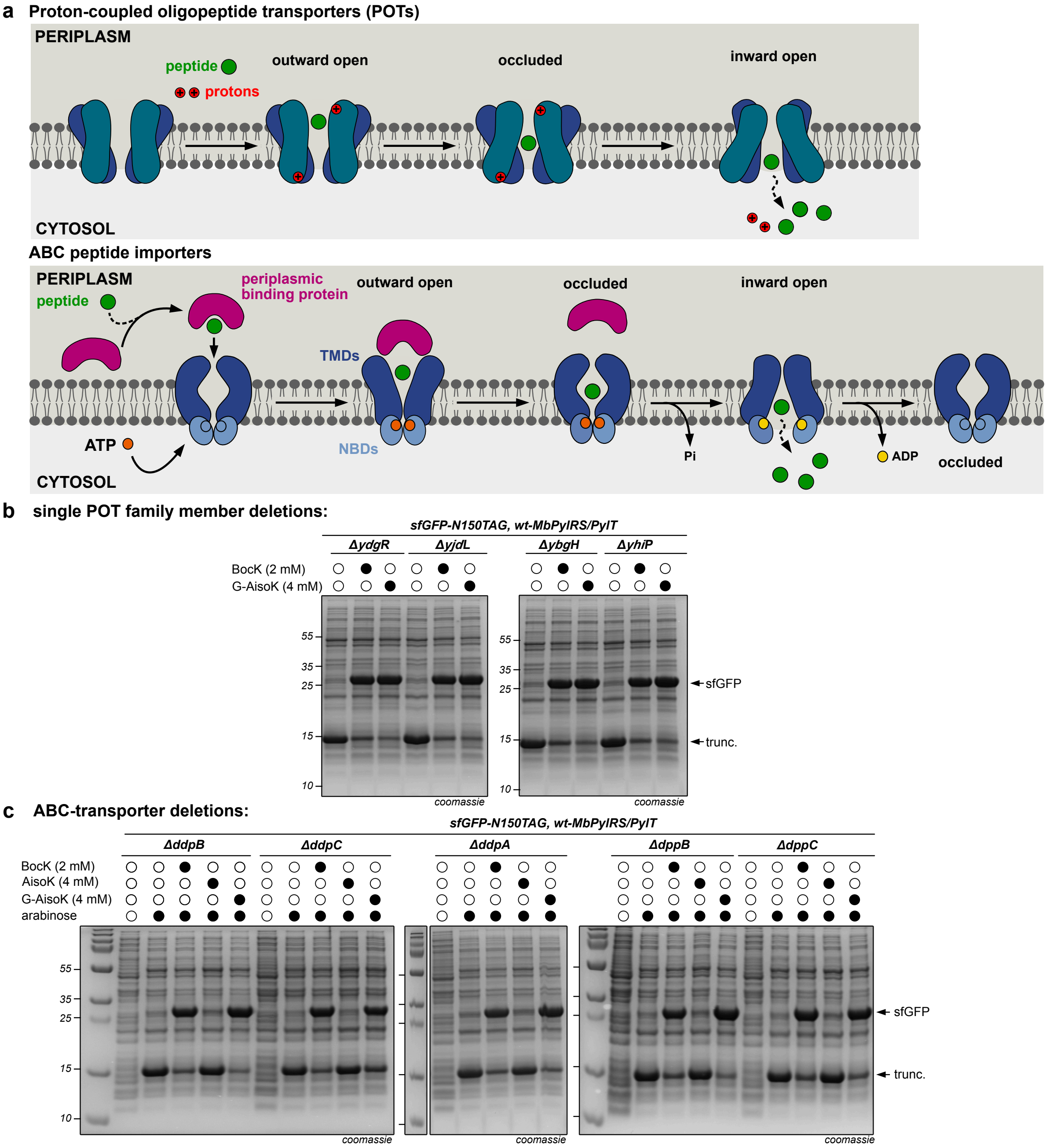

**Figure S2:** Identifying bacterial transporter systems for G-AisoK uptake **a.** Top: scheme illustrating the transport mechanism of Proton-coupled oligopeptide transporters (POTs). Peptides are co-transported along a proton gradient into the cell. Bottom: scheme illustrating the mechanism of ATP-binding cassette (ABC) peptide importers. A periplasmic binding protein binds to and shuttles the peptide to the periplasm-facing site of the transmembrane domains (TMDs) (outward open conformation), which triggers ATP binding and dimerization of the two cytoplasmic nucleotide-binding domains (NBDs) that bind and hydrolyze ATP, leading to uptake of the peptide into the translocation channel (occluded conformation). Hydrolysis of ATP leads to conformational changes in the NBDs and dissociation of their dimerization interface and flips the TMDs to trigger release of the substrate peptide into the cytoplasm (inward open state). Once the peptidic ligand is shuffled into the cytosol, the apo form of the periplasmic binding protein and bind to another peptide. **b.** Single gene knockouts (KOs) of POT family members *ydg*R, *yjd*L, *ybg*H and *yhi*P. None of these KOs abolished the expression of full length sfGFP-N150TAG in the presence of G-AisoK. **c.** Single gene KOs of other oligopeptide ABC transporters *ddp*A,B,C and *dpp*B,C. None of these KOs abolished expression of full length sfGFP-N150TAG in the presence of G-AisoK.

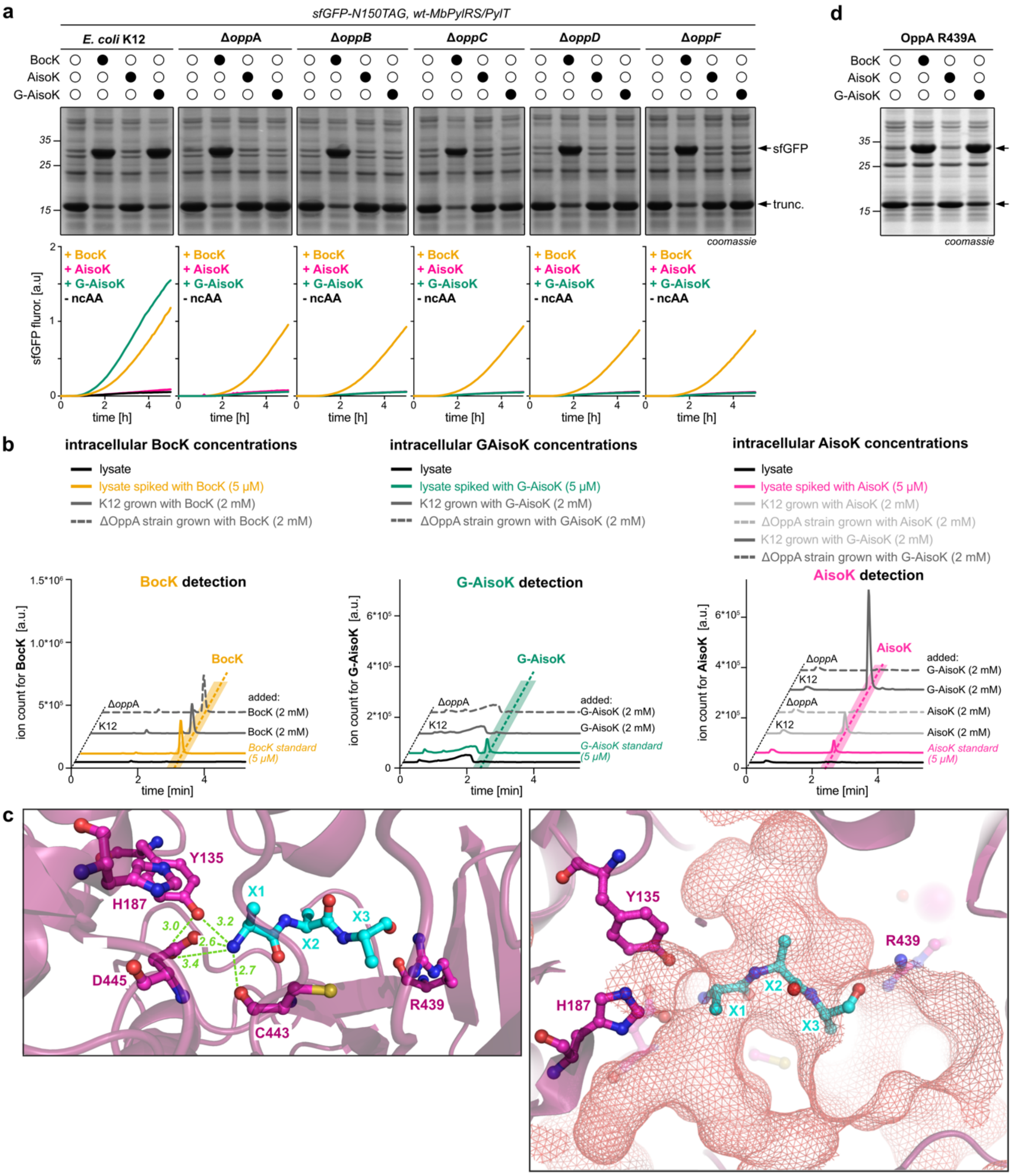

**Figure S3:** The Opp transporter is responsible for G-AisoK uptake. **a.** SDS-PAGE analysis (top) and time course fluorescence measurements (bottom) of K12 cells and individual knockout cell lines that have individual components of the Opp transporter genomically deleted, expressing sfGFP-N150TAG in the presence of BocK, AisoK or G-AisoK using wt-*Mb*PylRS/PylT. Full length sfGFP expression in presence of G-AisoK is abolished in all KOs (Δ*opp*A-F), while sfGFP expression in presence of BocK remains comparable between wt K12 cells and KO cell lines, indicating that the Opp transporter is responsible for G-AisoK uptake and associated amber codon suppression. **b.** LC-MS traces of extracted ion chromatograms to determine intracellular concentrations of BocK (left), G-AisoK (middle) and AisoK (right) in wt-K12 and Δ*opp*A cells. Intracellular BocK concentrations for cells grown with 2 mM BocK are similar between wt-K12 and Δ*opp*A-K12 (left: dashed, dark grey). Intracellular G-AisoK concentrations were negligible for both cell types (middle). In Δ*opp*A cells intracellular AisoK concentrations were negligible for cells grown in 2 mM AisoK (dashed, light grey) as well as 2 mM G-AisoK (right, dashed, dark grey). **c.** X-ray crystal structure of OppA bound to a linear bound tripeptide XXX (PDB ID: 3TCF[1], side chains are depicted as alanines, but are not resolved in the crystal structure). Left: peptide binding site shows that the peptide N-terminus is held in place by an extensive H-bonding network, interacting with the sidechains of D445 and Y135 and the backbone of C443. Hydrogen-bonding of D445 to Y135 and H187 lowers its pKa and keeps it in its deprotonated form, facilitating salt bridge formation with the positively charged N-terminus amine. Right: peptide binding site of OppA shown with binding pocket cavities denoted as a red mesh. Cavities highlight the space available in positions X1, X2 that can potentially accommodate larger sidechains. **d.** SDS-PAGE analysis of sfGFP-N150TAG expression in Δ*opp*A cells, constitutively expressing the OppA-R439A variant. sfGFP expression in presence G-AisoK remains unchanged in cells expressing the OppA-R439A mutant, indicating R439 is not involved in G-AisoK binding and uptake.

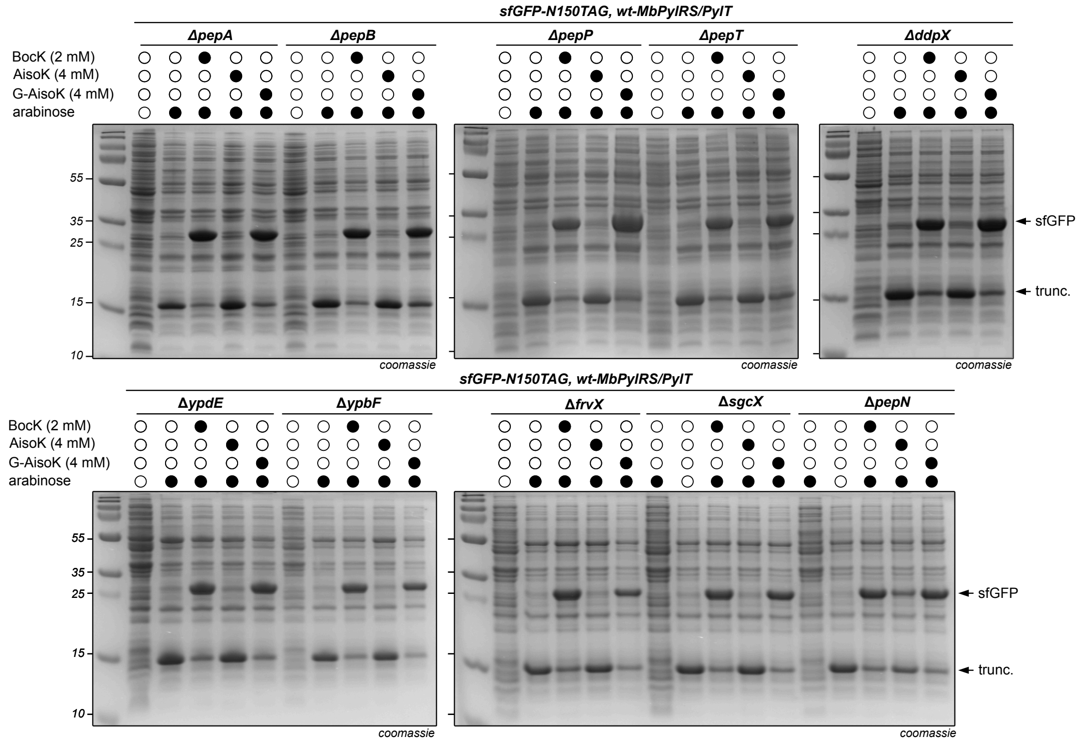

**Figure S4:** SDS-PAGE analysis of sfGFP-N150TAG expression in K12 knockout cell lines that had individually single amino peptidase enzymes *pep*A, *pep*B, *pep*P, *pep*T, *ddp*X, *ypd*E, *ypb*F, *frv*X, *sgc*X and *pep*N genomically deleted. None of these KOs abolished expression of full length sfGFP-N150TAG in the presence of G-AisoK and wt-*Mb*PylRS/PylT, suggesting there is no single peptidase that is responsible for the cleavage of the N-terminal glycine of G-AisoK.

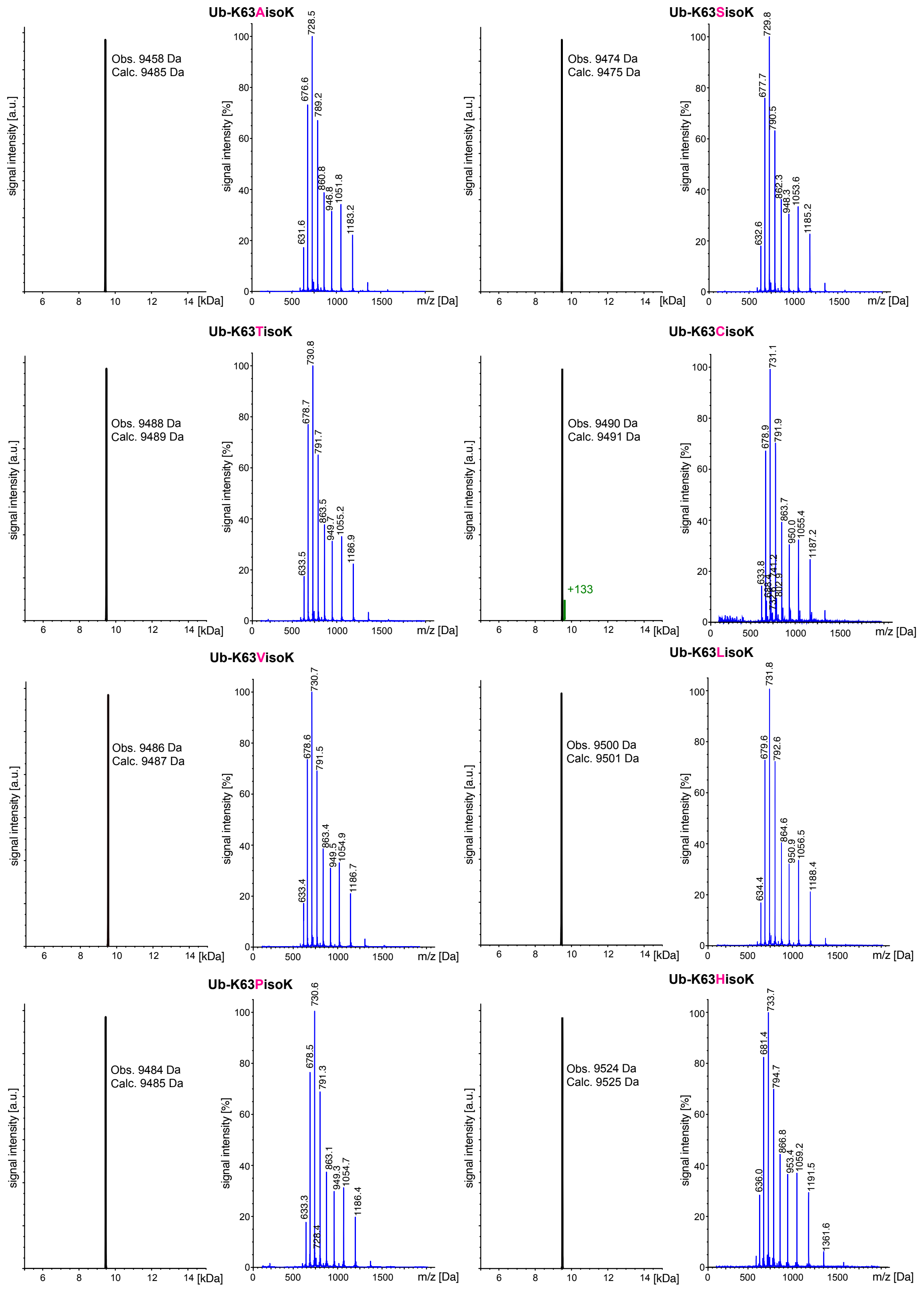

**Figure S5.** LC-MS analysis of purified Ub-K63XisoK expressed in the presence of G-XisoK derivatives, in which X stands for a natural amino acid. All observed masses confirm incorporation of the corresponding XisoK derivatives. Ub-K63CisoK was treated with methoxyamine to remove metabolic adducts.

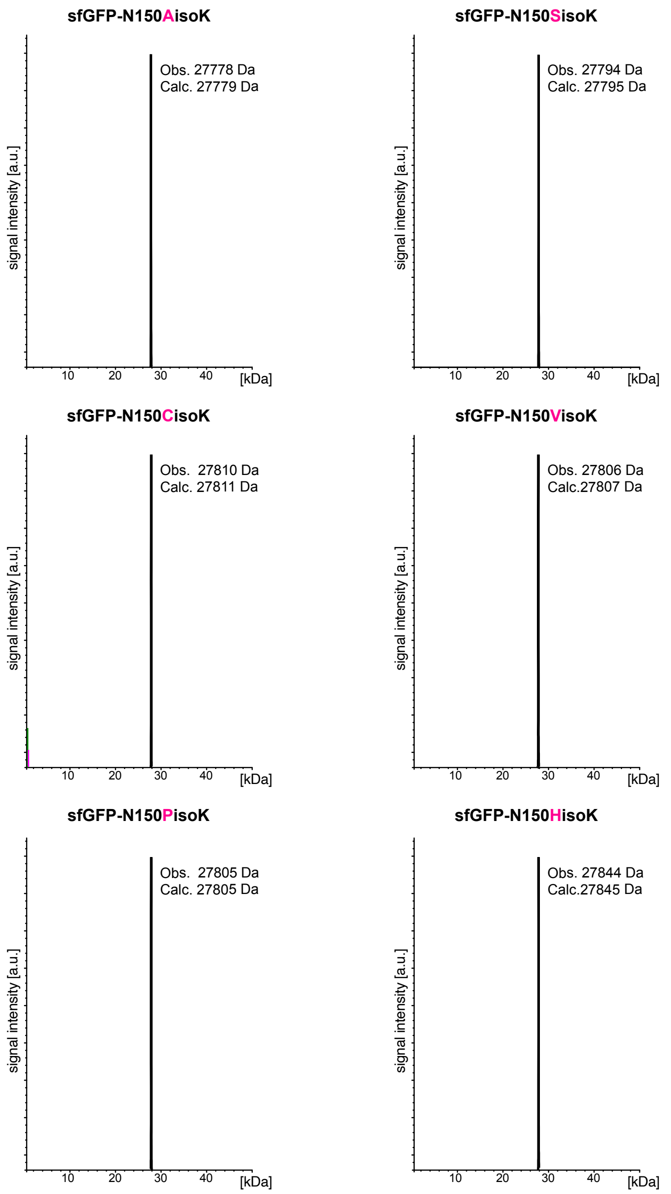

**Figure S6:** LC-MS analysis of purified sfGFP-N150XisoK expressed in the presence of G-XisoK derivatives, where X is a natural amino acid. All observed masses confirm incorporation of the corresponding XisoK derivatives into sfGFP. sfGFP-N150CisoK was treated with methoxyamine to remove metabolic adducts.

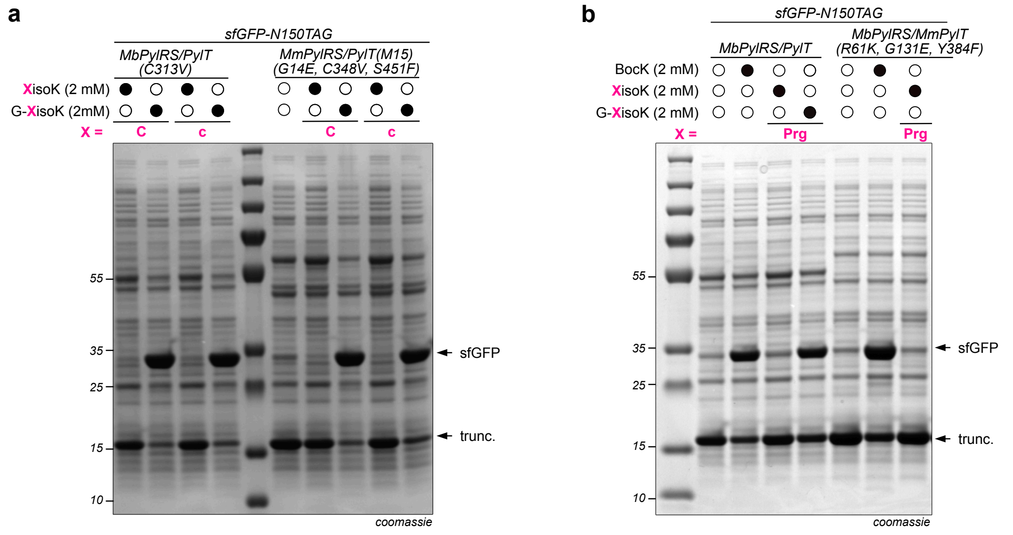

**Figure S7:** Comparison of incorporation efficiencies using the G-XisoK scaffold versus previously reported evolved PylRS variants for CisoK and PrgisoK. (C = L-cysteine, c = D-cysteine) **a**. SDS-PAGE analysis of sfGFP-N150TAG expression with G-CisoK, CisoK, G-cisoK and cisoK using *Mb*PylRS(C313V)/PylT used in this study or previously reported aaRS/tRNA pair for cisoK incorporation (*Mm*PylRS(G14E,C348V,S451F)/*Mm*PylT(M15[2]))[3]. Similar expression yields for full-length sfGFP are obtained for G-CisoK and G-cisoK with both aaRS/tRNA pairs, while nearly no full-length sfGFP expression is observed using the dipeptides CisoK/cisoK and either of the aaRS/tRNA pairs, indicating the superiority of our approach. **b.** SDS-PAGE analysis of sfGFP-N150TAG expression with BocK, PrgisoK and G-PrgisoK comparing wt-*Mb*PylRS/PylT used in this study with a previously reported PylRS/tRNA pair for PrgisoK incorporation (*Mb*PylRS(R61K,G131E,Y348F) /*Mm*PylT)[4]. Full-length sfGFP expression is significantly higher using G-PrgisoK and wt-*Mb*PylRS/PylT in comparison to the previously published synthetases with dipeptide PrgisoK.

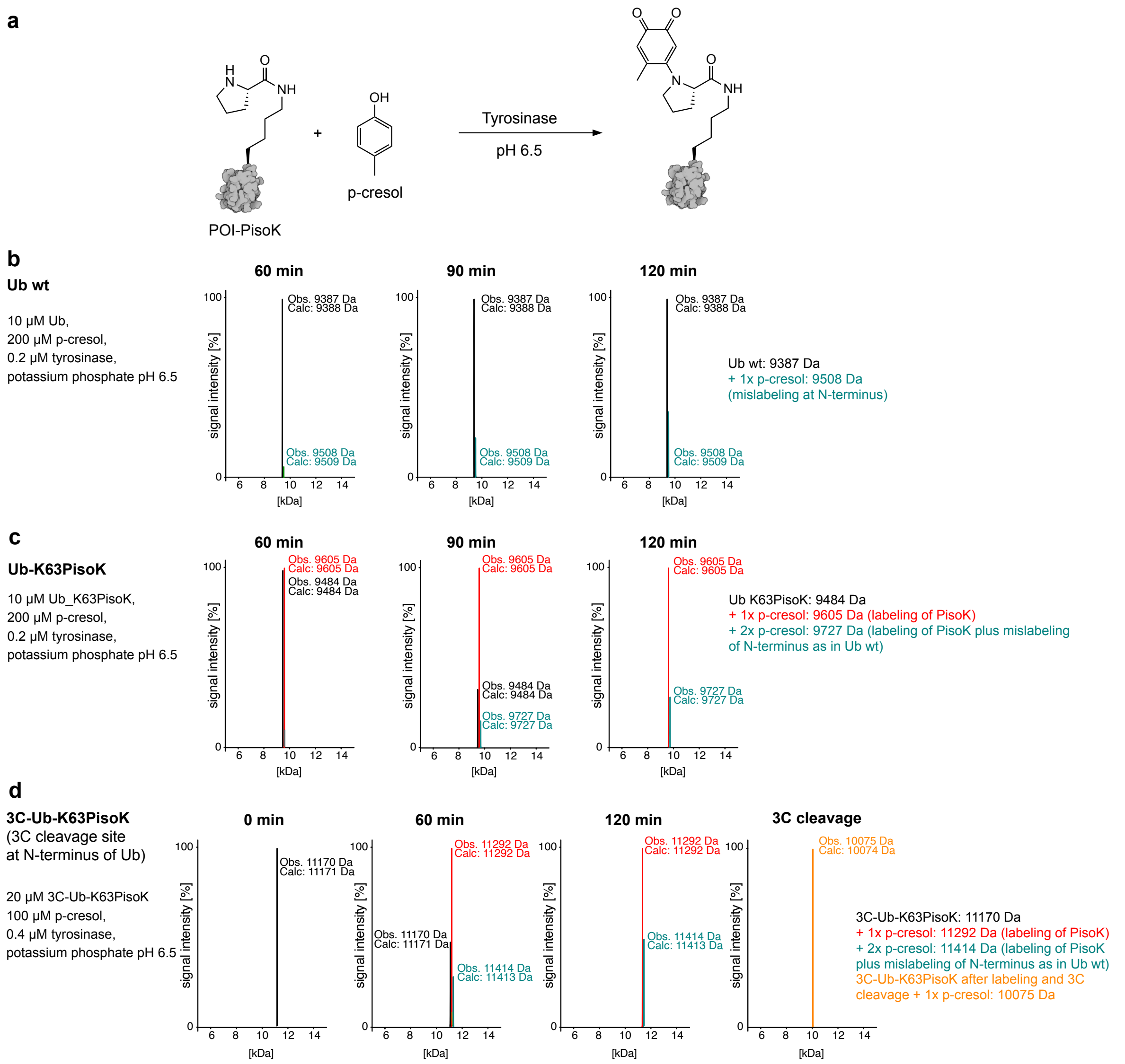

**Figure S8:** Tyrosinase-mediated labeling of PisoK-bearing proteins. **a.** Scheme of labeling proteins of interest (POI) bearing PisoK with model substrate p-cresol mediated by tyrosinase. **b.** Incubation of Ub wt (10 µM) with p-cresol (200 µM) in the presence of 0.2 µM tyrosinase at pH 6.5, r.t. shows mislabeling at Ub N-terminus (green peak) after 120 min, as observed by LC-MS analysis. **c.** Incubation of 10 µM Ub-K63PisoK with 200 µM p-cresol with 0.2 µM tyrosinase after 120 minutes gives complete labeling of PisoK and some mislabeling of the N-terminus as observed for wt-Ub. **d.** To confirm quantitative Ub-K63PisoK labeling, Ub bearing an N-terminal 3C protease cleavage site was incubated under the same conditions with 0.4 µM tyrosinase for 120 min, after which the labeled protein was subjected to 3C cleavage to reveal quantitative internal PisoK labeling.

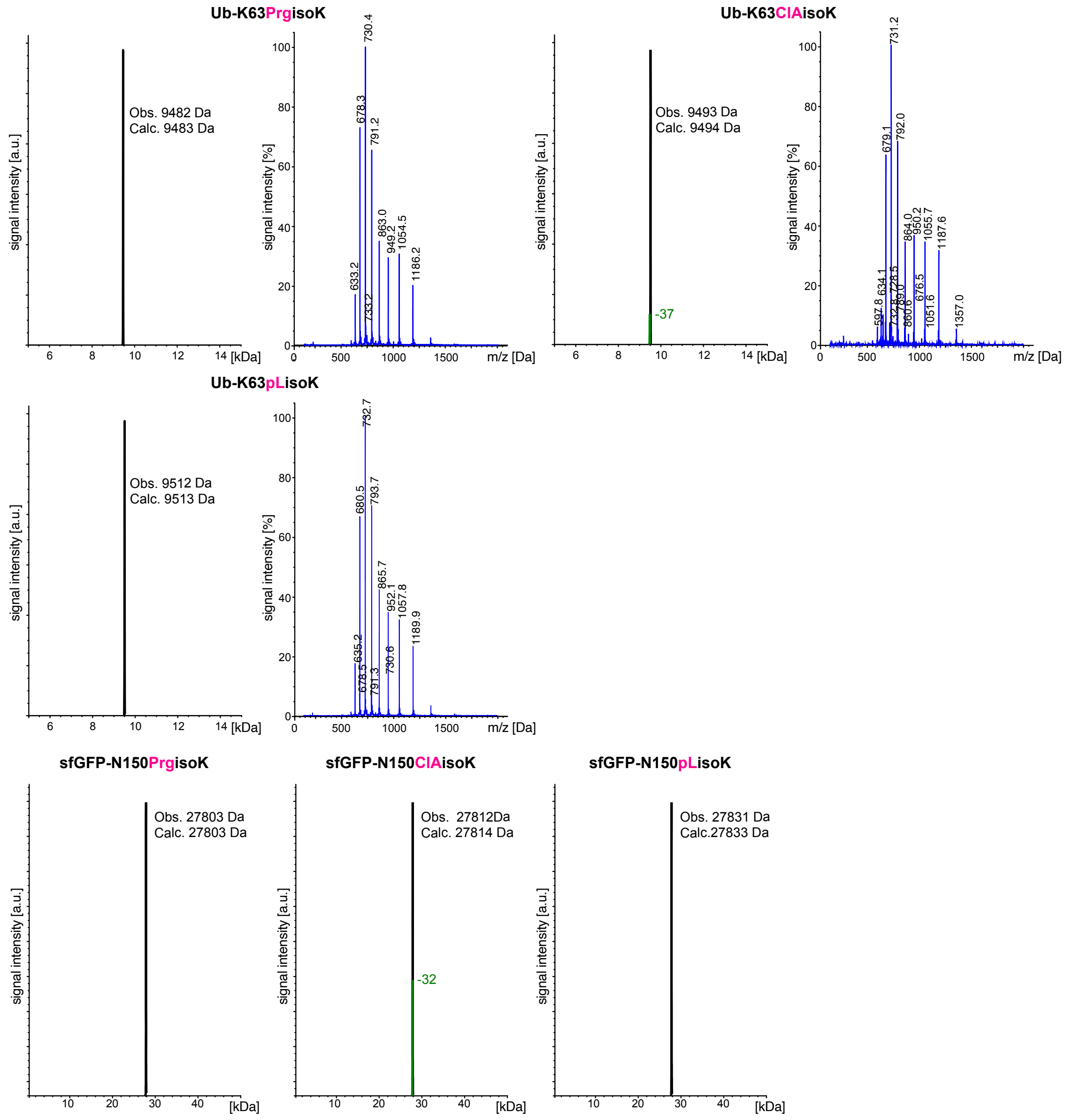

**Figure S9:** LC-MS analysis of purified Ub-K63XisoK and sfGFP-N150XisoK expressed in the presence of G-XisoK derivatives, where X is an ncAA. Observed masses confirm incorporation of the corresponding XisoK derivatives. Peaks denoted in green Ub-K63ClAisoK and sfGFP-N150ClAisoK correspond to the elimination of HCl.

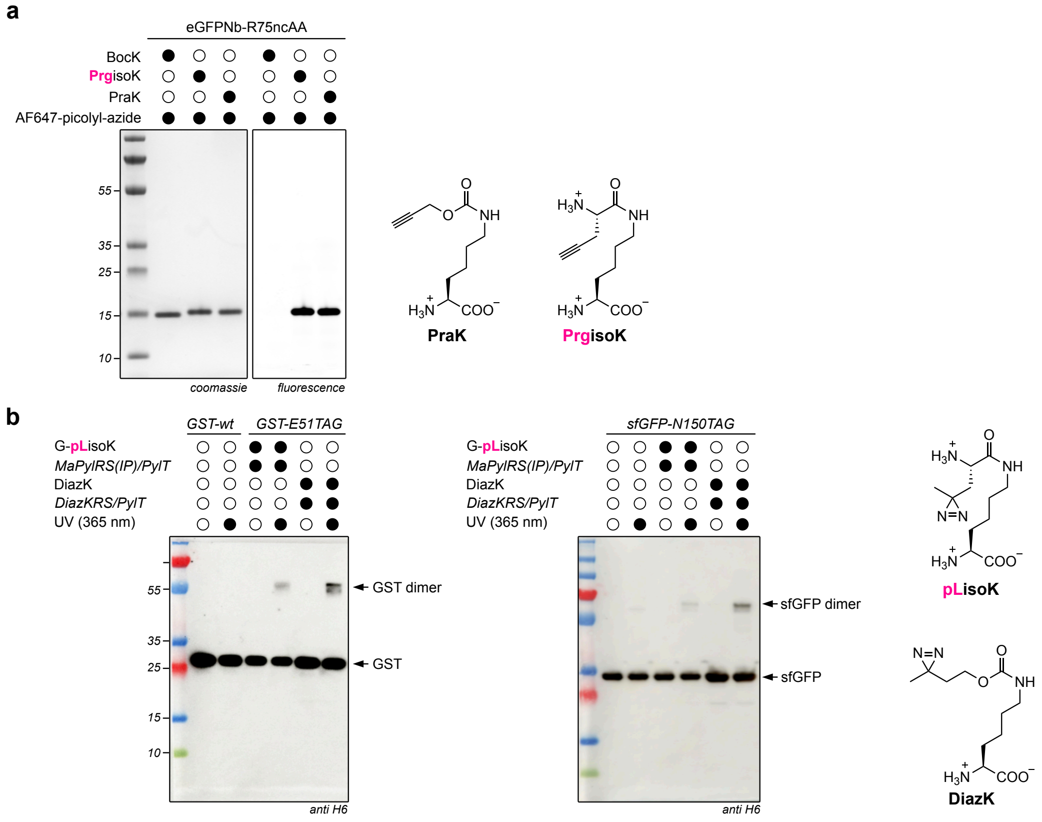

**Figure S10:** XisoK ncAAs for bioorthogonal labeling and photocrosslinking. **a.** CuAAC-based labeling of proteins bearing a terminal alkyne moiety. Purified eGFPNb bearing either PrgisoK or the previously reported ncAA PraK[5] at position 75 are labeled with comparable efficiencies with an azide-bearing fluorophore (conditions: 100 µM AF647-Picolyl-Azide, 50 µM $CuSO_4$, 250 µM THPTA and 2.5 mM ascorbic acid in PBS for 30 minutes at room temperature). BocK-bearing eGFPNb is not labeled under the same conditions, as expected. **b.** Photo-crosslinking of diazirine-bearing glutathione-transferase (GST) and sfGFP. left: Western blot analysis of cells expressing GST-E51TAG in the presence of G-pLisoK and the previously reported DiazK[6]. Upon $UV_{365\ nm}$ illumination both G-pLisoK and DiazK samples show cross-linked GST dimers. Right: Western blot analysis of cells expressing sfGFP-N150TAG in the presence of G-pLisoK and DiazK. Illumination with $UV_{365\ nm}$ for G-pLisoK and DiazK samples show cross-linked sfGFP dimers. (conditions: cells expressing POIs were diluted to $OD_{600}$ = 1 and irradiated for 15 minutes at 365 nm, 15 W)

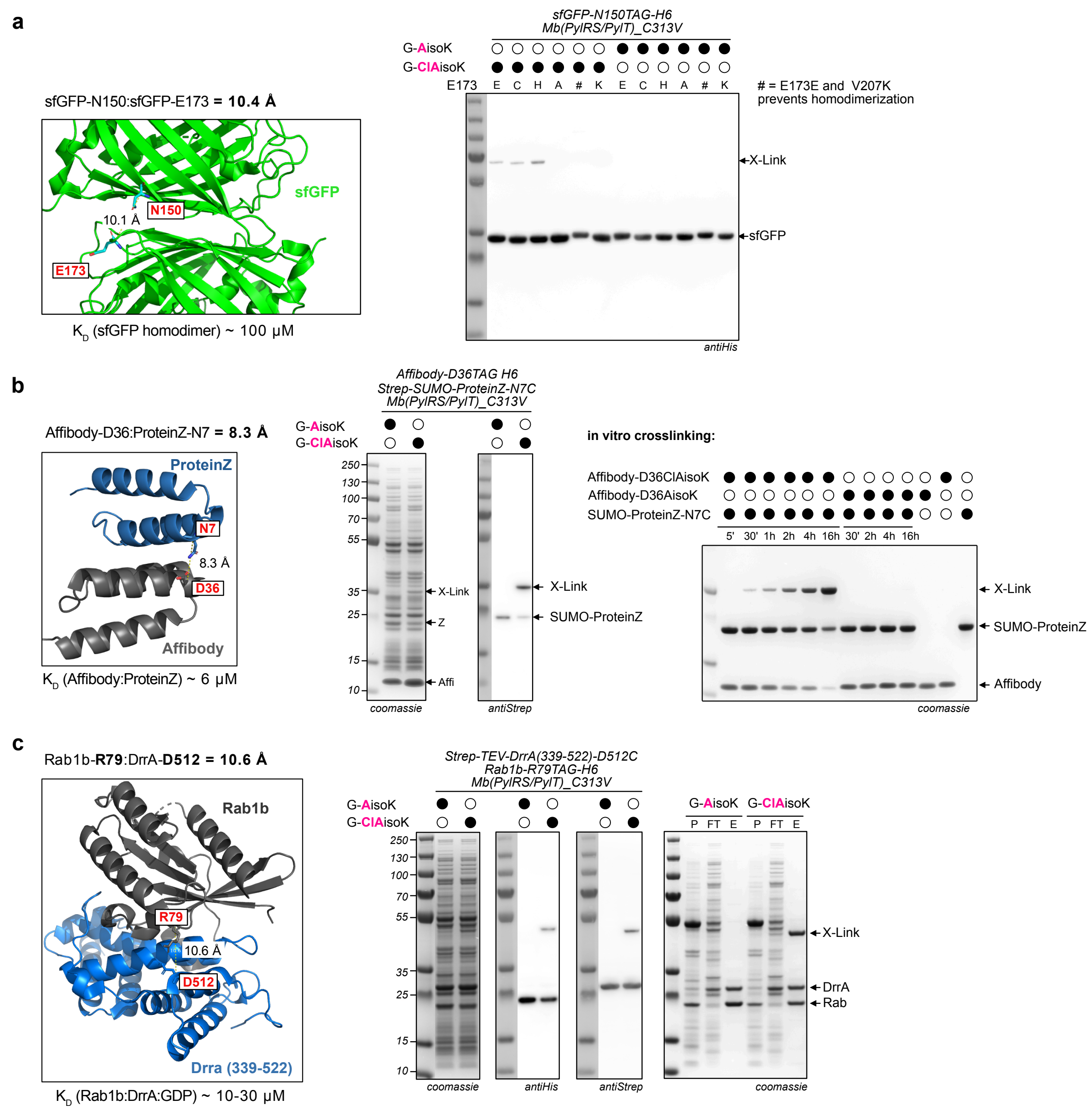

**Figure S11:** Proximity-induced chemical crosslinking using ClAisoK-bearing proteins. **a.** Chemical crosslinking of ClAisoK-bearing sfGFP homodimer. Left: X-ray crystal structure of the sfGFP dimeric interface (PDB ID :1GFL[7]) denoting position N150 (where ncAAs are incorporated) and E173. The distance between the two Cα atoms of these two amino acids is 10.4 Å; the $K_D$ of sfGFP dimerization has been determined with 100 µM[8]. Right: Western blot analysis of cells expressing sfGFPN150TAG in the presence of G-AisoK or G-ClAisoK. Cross-linked sfGFP homodimer is observed for samples grown in the presence of GClAisoK and with residue 173 mutated to E, C or H. # denotes a sfGFP variant, in which a mutation (V207K) that disrupts the dimerization interface has been introduced. **b.** Chemical crosslinking of protein Z with selected affibody[9]. Left: X-ray crystal structure of affibody-proteinZ complex (PDB ID: 1LP1[9]) highlighting residues D36 in affibody (position for ncAA incorporation) and N7 in protein Z, which is mutated to cysteine to allow for crosslinking. The distance between the two Cα atoms of these two residues is 8.3 Å; the $K_D$ has been determined to be 6 µM[9]. Middle: In cellulo crosslinking: SDS-PAGE and Western blot analysis of cells expressing Strep-tagged SUMO2-proteinZ-N7C and affibody-D36TAG-H6 in the presence of either G-AisoK or G-ClAisoK. Cross-linked complex is only observed for the G-ClAisoK sample. Right: SDS-PAGE analysis of *in-vitro* crosslinking with purified binding partners. Cross-linked complex is observed for ClAisoK-bearing affibody after 30 minutes and near quantitative covalent crosslinking between affibody and protein Z is observed after 16 h. No crosslinking is observed for AisoK-bearing affibody. **c.** Complete data from Fig. 3f. Chemical crosslinking between ClAisoK-bearing Rab1b GTPase and Legionella effector protein $DrrA_{339-522}$[10]. Left: X-ray crystal structure of Rab1b-$DrrA_{339-522}$ complex (PDB ID: 3JZA[11]), highlighting R79 in Rab1b (changed to ncAA) and D512 in DrrA (mutated to cysteine). The distance between the two Cα atoms of these two residues is 11.4 Å; the $K_D$ of ternary complex formation in presence of GDP has been determined to be ca. 10-30 μM[11]. Western blot analysis of cells expressing both binding partners in the presence of either G-AisoK or G-ClAisoK shows crosslinked complex formation only in the presence of G-ClAisoK (middle). Purification via Ni-NTA beads enriches both binding partners for both G-AisoK and G-ClAisoK, but covalently crosslinked complex is only present in G-ClAisoK sample (right). P = pellet, FT = flow through and E = elution.

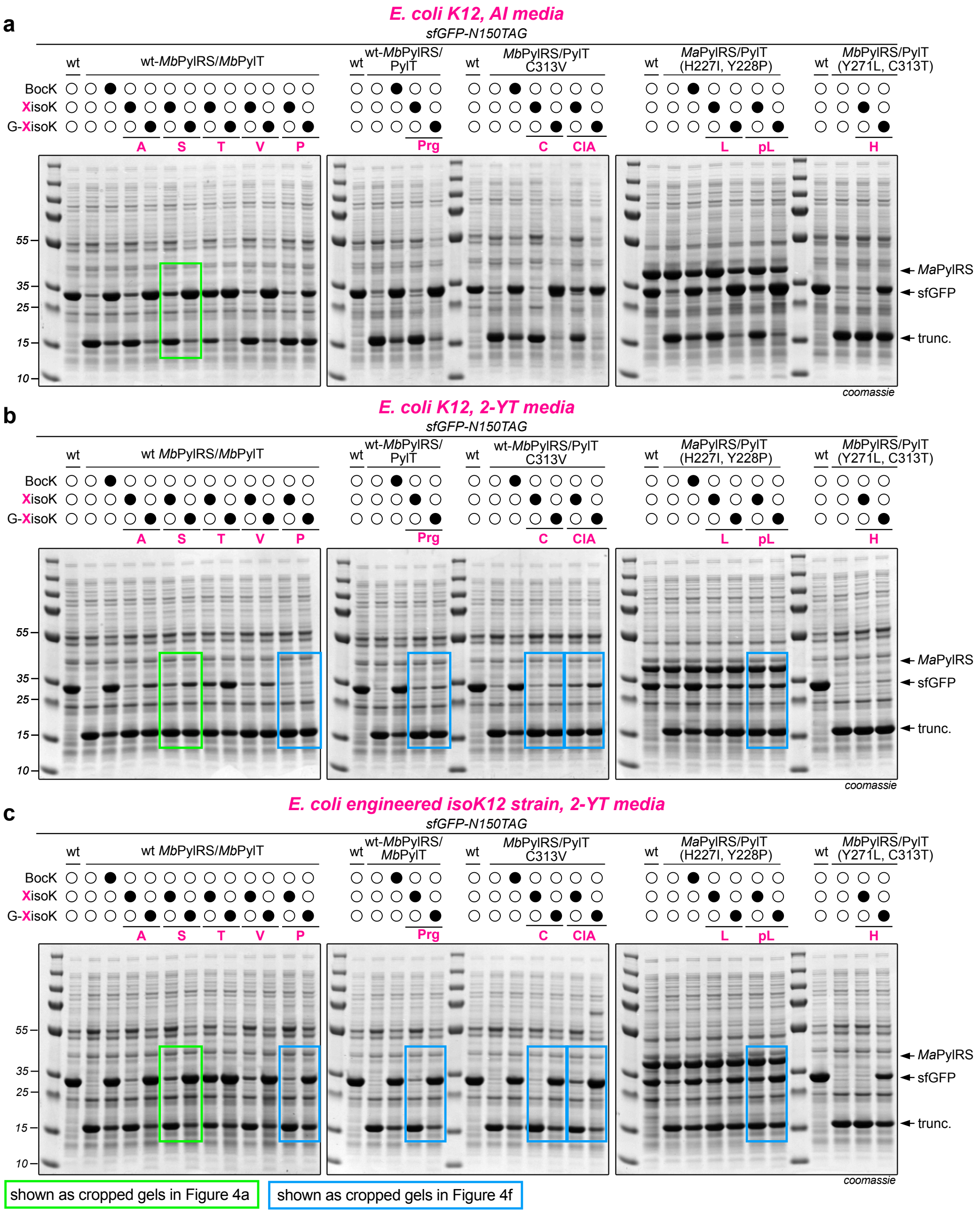

**Figure S12:** SDS-PAGE analysis of sfGFP-N150TAG expression in the presence of all G-XisoK and XisoK derivatives used in this study, comparing amber suppression levels in K12 cells versus engineered IsoK12 cells grown in different media. **a.** Expression in K12 cells using autoinducing (AI) media. **b.** Expression in K12 cells using 2-YT media. **c.** Expression in engineered IsoK12 cells using 2-YT media.

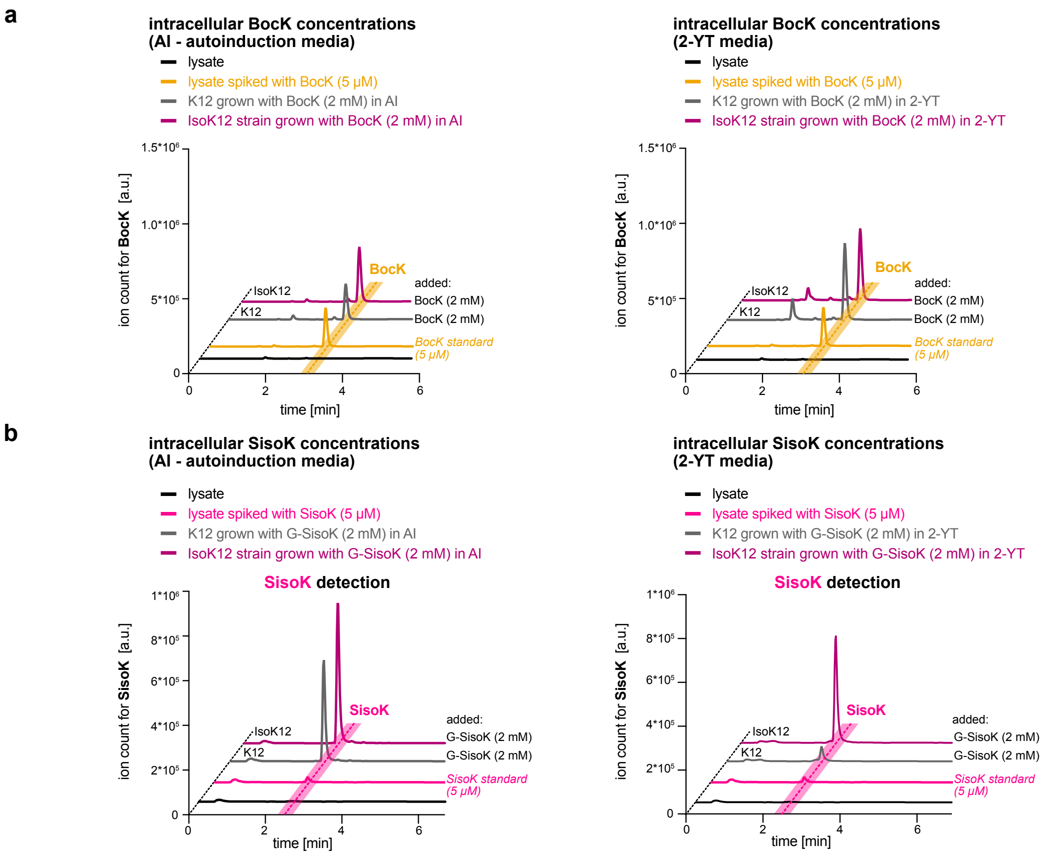

**Figure S13:** LC-MS traces of cell lysates in selected-ion mode to determine intracellular concentrations of ncAAs. **a.** Extracted ion chromatograms to determine intracellular BocK concentrations. Wt-K12 cells or IsoK12 cells are grown in presence of 2 mM BocK using AI media (left) or 2-YT media (right). BocK concentrations in lysate are comparable for both cell types in both media. **b.** Extracted ion chromatograms to determine intracellular SisoK concentrations. Wt-K12 cells or IsoK12 cells grown in presence of 2 mM G--SisoK using AI media (left) or 2-YT media (right). SisoK concentrations in AI media are ca. 1.4-fold higher in IsoK12 cells compared to wt-K12 cells (left). In 2-YT media, SisoK concentrations are 7-10-fold higher in IsoK12 cells (purple) compared to wt-K12 cells (grey) (right).

**a**

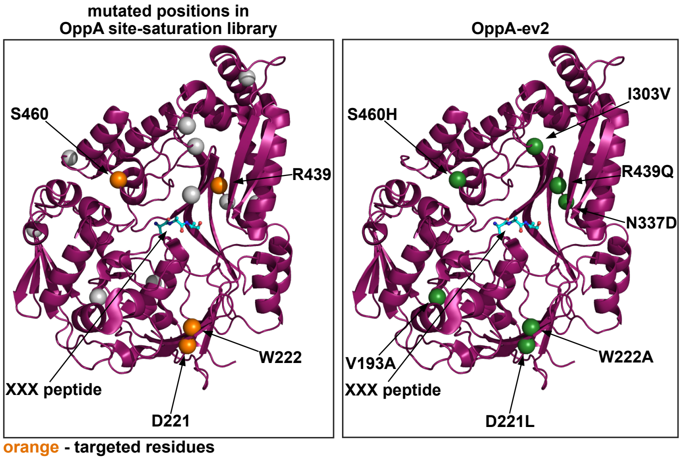

**b**

| OppA variant | mutations<br>**bold** - targeted residues |
|---|---|
| error-prone variant 1 | T173N, **D221**G, K371E, R439H |
| error-prone variant 2 | L78I, **W222**R, K307R, K333N |
| error-prone variant 3 | T429A, **R439**L, **S460**C, V482A |
| error-prone variant 4 | V193A, I303V, N337D, **S460**N |
| site-saturation variant OppA-ev2 | V193A, **D221**L, **W222**A, I303V, N337D, **R439**Q, **S460**H |

**Figure S14:** Evolved OppA-variant OppA-ev2. **a.** X-ray crystal structure of OppA (PDB ID: 3TCF[1]) with positions that were mutated denoted as spheres. Left: positions that were mutated in error prone OppA variants 1-4. Orange spheres: positions that were targeted for a site-saturation OppA library. Grey spheres: other positions that were mutated in the error prone OppA variants. Right: green spheres show positions that were mutated in the final variant OppA-ev2. **b.** Table indicating all the mutations in error prone OppA variants 1-4 and mutations in the final variant OppA-ev2. Bold residues were targeted for site saturation.

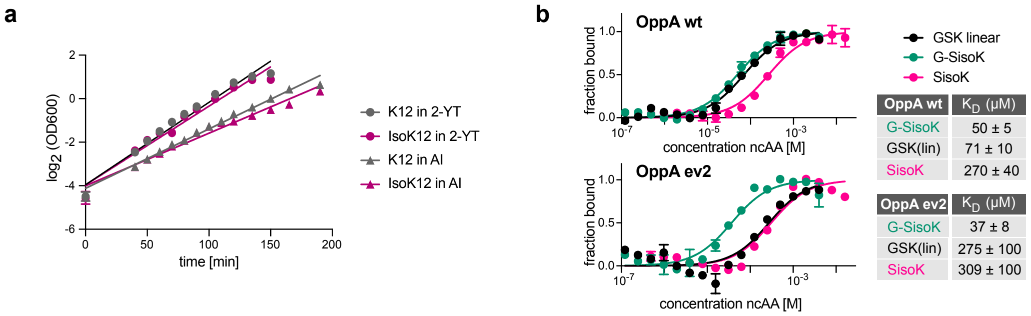

**Figure S15:** Characterization of evolved IsoK12 strain. **a.** Doubling times of wt-K12 cells (grey) and IsoK12 cells (purple) grown in AI media and 2-YT. Doubling times for IsoK12 in 2-YT was 19 minutes compared to K12, which was 18 minutes. In AI media, IsoK12 had a doubling time of 29 minutes in comparison to K12, which was 25 minutes. **b.** Microscale thermophoresis affinity measurements for GSK linear (GSK(lin)), G-SisoK and SisoK towards wt-OppA and evolved OppA ev2. Affinity towards SisoK and G-SisoK remained similar for both wt-OppA and OppA-ev2 (50 µM vs 27 µM). In contrast affinity towards a linear GSK tripeptide is reduced ~ 4-fold for OppA-ev2.

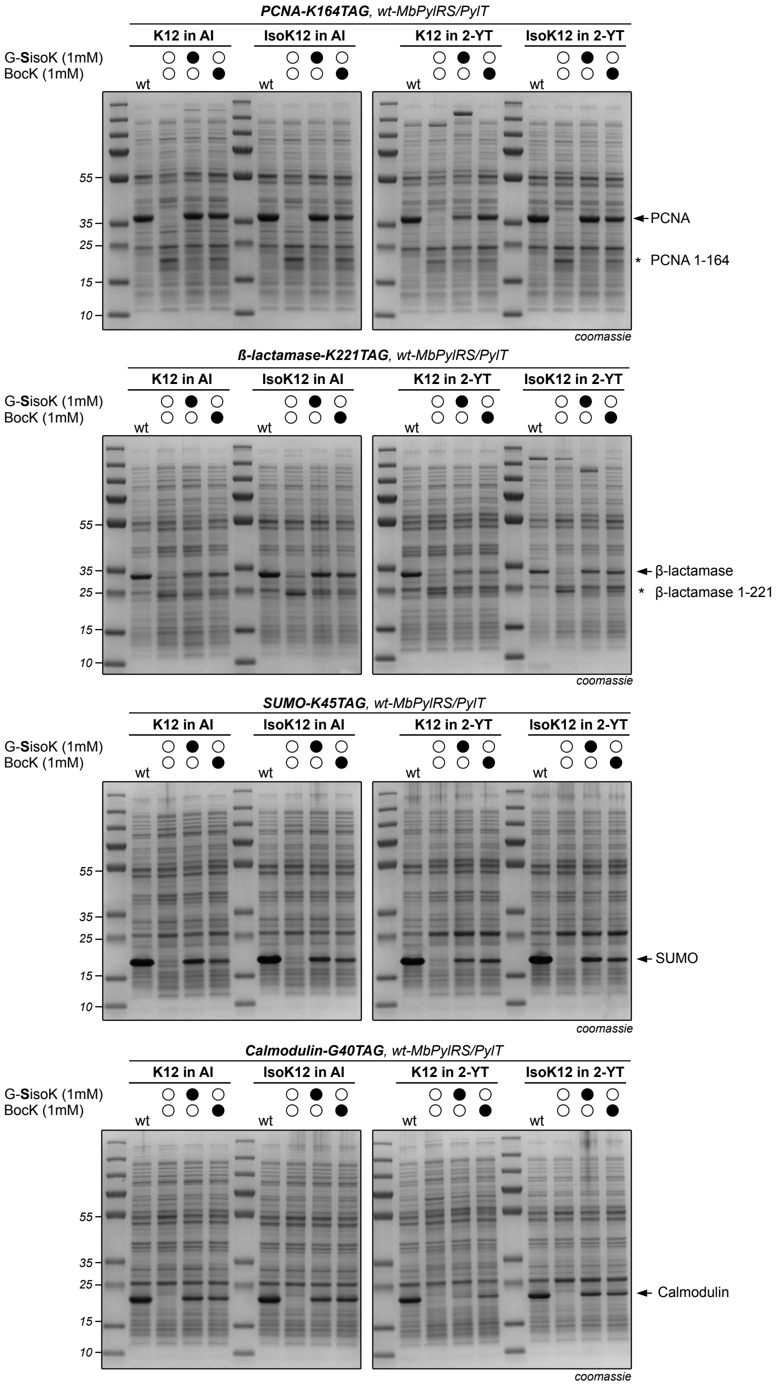

**Figure S16:** SDS-PAGE analysis of amber suppression efficiencies within different POIs in the presence of 1 mM BocK or G-SisoK in wt-K12 and IsoK12 cells comparing AI media with 2-YT. From the top: PCNA-K164TAG, β-lactamase-K122TAG, SUMO-K45TAG and Calmodulin-G40TAG.
In all cases, amber suppression efficiency for G-SisoK is considerably higher in IsoK12 cells than in wt-K12 cells grown in 2-YT. Arrows indicate full-length POIs, asterisks indicate truncated proteins.

a

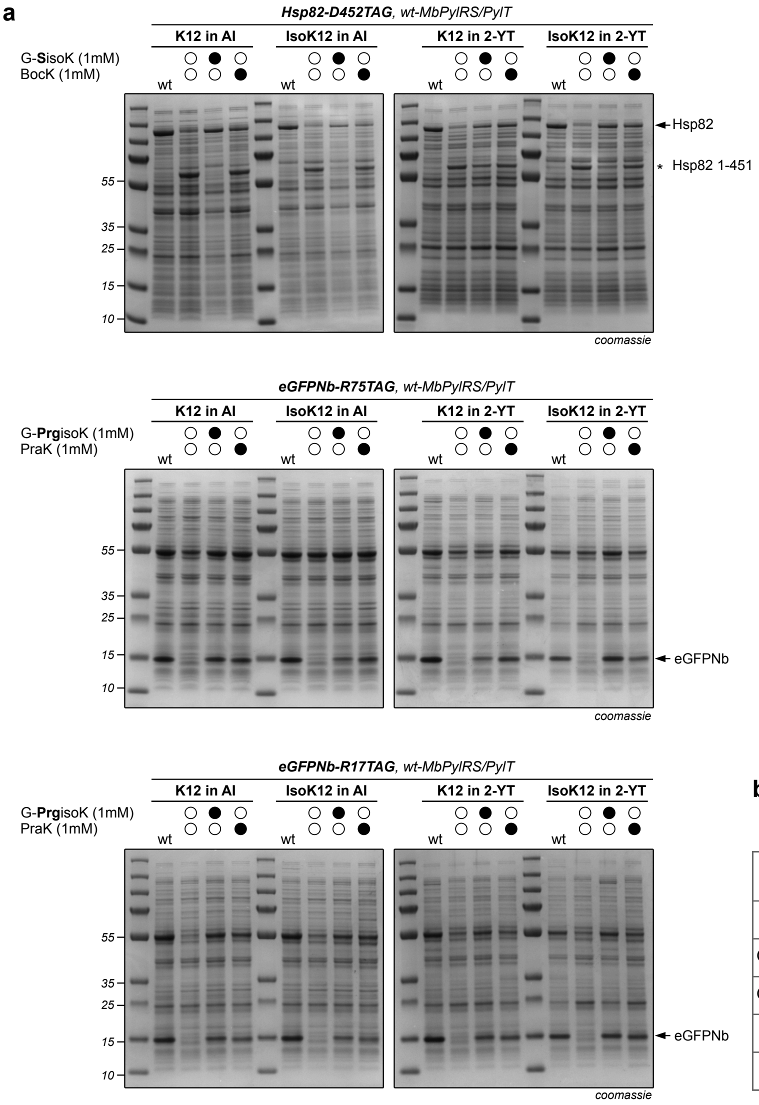

b

| added ncAA | expression strain | eGFPNb-H6 variant | yield: [mg/L] |
|---|---|---|---|
| - | wt-K12 | wt | 41 |
| G-PrgisoK | wt-K12 | R17PrgK | 22 |
| G-PrgisoK | IsoK12 | R17PrgK | 44 |
| PraK | IsoK12 | R17PraK | 29 |
| PraK | wt-K12 | R17PraK | 27 |

**Figure S17: a.** SDS-PAGE analysis of amber suppression efficiencies within different POIs in the presence of 1 mM BocK, G-SisoK, PraK[5] or G-PrgisoK in wt-K12 and IsoK12 cells comparing AI media and 2-YT media. Top: Hsp82-D452TAG, middle: eGFPNb-R75TAG, bottom: eGFPNb-R17TAG. Full-length protein expression is higher in IsoK12 cells compared to wt-K12 cells in 2-YT. **b.** Preparative protein yields of wt-eGFPNb, eGFPNb-R17PrgisoK or eGFPNb-R17PraK. For PraK structure see Fig. S10a. wt-*Mb*PylRS/PylT pair used for PraK and G-PrgisoK incorporation.[5] PrgisoK-bearing eGFPNb yields exceed yields for PraK-bearing eGFPNb, matching wt eGFPNb yields. Arrows indicate full-length POIs, asterisks indicate truncated proteins.

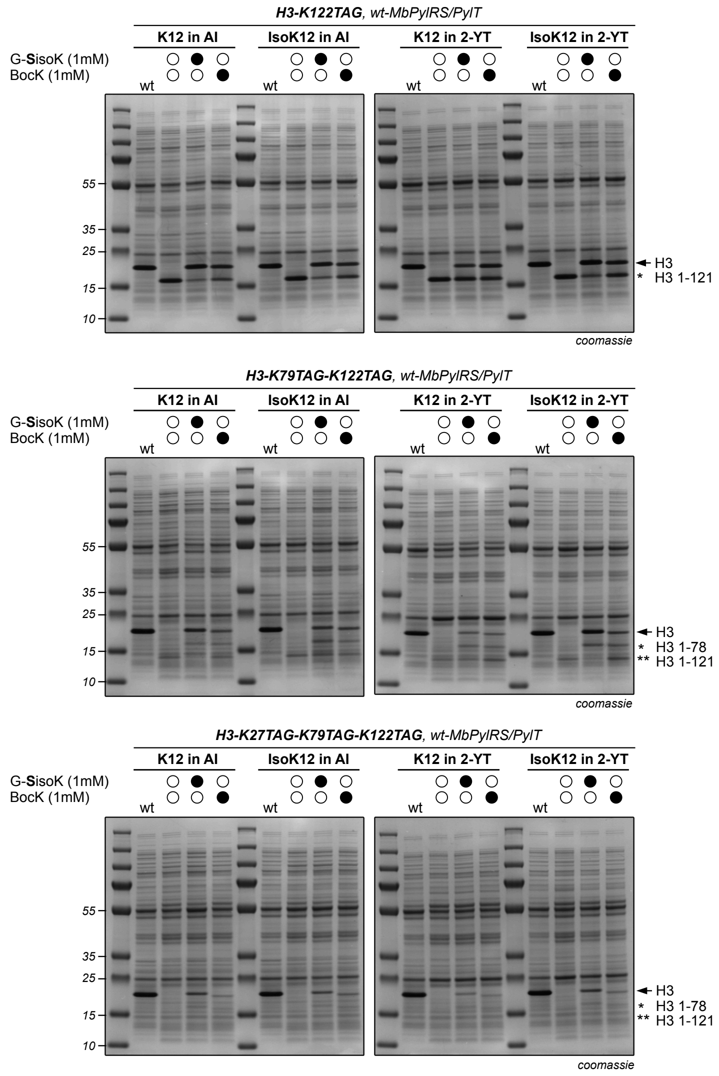

**Figure S18:** Full gels of Figure 4g. SDS-PAGE analysis of amber suppression efficiencies for Histone H3 variants bearing either 1 TAG codon (K122TAG, top), 2 TAG codons (K79TAG, K122TAG, middle) or 3 TAG codons (K27TAG, K79TAG, K122TAG, bottom) in wt-K12 or IsoK12 cells, comparing expression in AI media or 2-YT media. Expression of all variants are higher in IsoK12 in 2-YT. Arrows indicate full-length POIs, asterisks indicate truncated proteins.

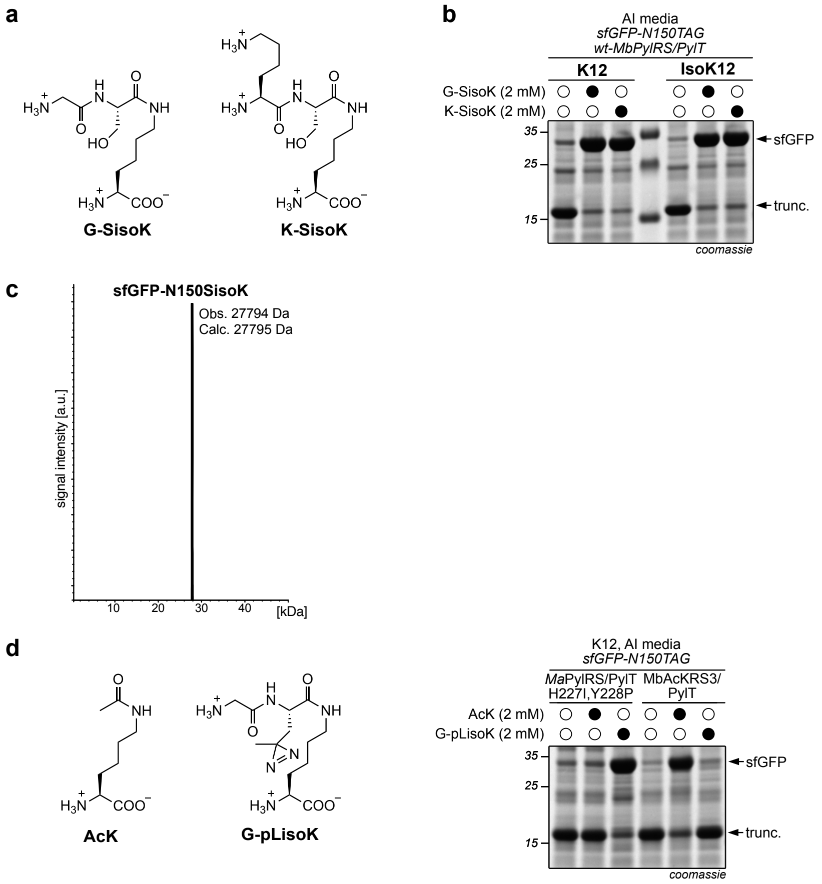

**Figure S19:** Dual incorporation of two different ncAAs using a single isopeptide linked tripeptide. **a.** Chemical structures of G-SisoK and K-SisoK. **b.** SDS-PAGE analysis of sfGFP-N150TAG expression in the presence of either G-SisoK or K-SisoK and wt-MbPylRS/PylT. Expression in presence of K-SisoK leads to similar sfGFP levels as in presence of G-SisoK, suggesting that the glycine can be replaced with lysine, maintaining uptake and intracellular cleavage. **c.** LC-MS analysis confirming the incorporation of SisoK at position 150 of sfGFP when expressed in presence of K-SisoK. **d.** Left: Chemical structures of ncAAs for dual stop codon suppression to allow for orthogonal incorporation of AcK and pLisoK. Right: SDS-PAGE analysis of sfGFP-N150TAG expression in the presence of 2 mM AcK or 2 mM G-pLisoK and their corresponding PylRS/PylT pairs to confirm substrate orthogonality between the pairs. Full-length sfGFP expression in presence of AcK is only observed with AcKRS3/PylT and in presence of G-pLisoK only observed with *Ma*PylRS(H227I,Y228P)/PylT, hence confirming orthogonality.

**General methods: Plasmids and Reagents**

Genes encoding GST-E51TAG, tyrosinase and Histone H3 mutants were ordered as DNA string from Twist biosciences and cloned into the pBAD vector via standard restriction cloning with enzymes from New England Biolabs. Single point mutants of OppA as well as insertion of a 3C site on UbK63 were done via Site-directed, Ligase-Independent Mutagenesis (SLIM)[12]. Recombination plasmid pSIJ8 was purchased from Addgene (Addgene ID: 68122). Oligonucleotide primers were ordered from Microsynth AG unless otherwise specified. General solvents and chemical reagents were purchased from Sigma Aldrich, cslabs, Fisher Scientific, Carbolution or Acros Organics. Fmoc building blocks of photoleucine, propagylglycine and chloroalanine were purchased from Iris Biotech. All reagents were used without further purification.

SDS-PAGE gels (Bolt™ Bis-Tris Plus Mini Protein Gels, 4-12%, Invitrogen) were run on a Bolt™ Mini Gel Tank (Invitrogen) system (165 V for 40 minutes) and stained with Quick Coomassie Stain (Generon). As a marker, PageRuler Prestained Plus Protein Ladder 10-250 kDa (ThermoFisher) was used. Proteins were transferred onto a nitrocellulose membrane using a Bio-Rad Trans-blot Turbo Transfer System. After transfer, the membrane was blocked with 5% skim milk in TBS-T buffer for 1 h at RT and incubated with the appropriate HRP-coupled antibody. Imaging of western blots was performed on an Amersham ImageQuant™ 800 using Immobilon® Forte Western HRP Substrate (Millipore).

Peptide and protein LC-MS was done using an Agilent Technologies 1260 Infinity LC-MS system with a 6310 Quadrupole spectrometer. Proteins were measured on a Phenomenex Jupiter C4 300 A LC Column (150 x 2 mm, 5 µm) and peptides, on a Luna Omega PS C18 (100 x 2.1 mm, 3 µm). The solvent system consisted of 0.1% formic acid in water (solvent A) and 0.1% formic acid in acetonitrile (solvent B). Both proteins and peptides were analyzed in positive mode.

Table 1: Plasmids used in this study

| **Plasmid** | **Resistance** | **Description** |
|---|---|---|
| pBAD_sfGFP_N150TAG_H6 | Ampicillin | sfGFP-N150TAG with a C-terminal His6 tag under an arabinose promoter |
| pBAD_sfGFP_wt_H6 | Ampicillin | sfGFP wt with a C-terminal His6 tag under an arabinose promoter |
| pBAD_Ub_K63TAG_H6 | Ampicillin | Ubiquitin-K63TAG with a C-terminal His6 tag under an |

| | | |
|---|---|---|
| | | arabinose promoter |
| pEVOL_*Mb*PylRS | Chloramphenicol | Two copies of *Mb*PylRS wt under an arabinose promotor and a glnS promotor and *Mb* $tRNA_{CUA}$ under a proK promoter. |
| pEVOL_*Mb*PylRS_C313V | Chloramphenicol | Two copies of *Mb*PylRS C313V under an arabinose promotor and a glnS promotor and *Mb* $tRNA_{CUA}$ under a proK promoter |
| pEVOL_*Ma*PylRS_IP | Chloramphenicol | Two copies of *Ma*PylRS H227I, Y228P under an arabinose promotor and a glnS promotor and *Ma* $tRNA_{CUA}$ under a proK promoter |
| pEVOL_HisoKRS | Chloramphenicol | Two copies of *Mb*PylRS Y271L C313T under an arabinose promotor and a glnS promotor and *Mb* $tRNA_{CUA}$ under a proK promoter |
| pEVOL_DiazKRS | Chloramphenicol | Two copies of DiazKRS under an arabinose promotor and a glnS promotor and *Mm* $tRNA_{CUA}$ under a proK promoter |
| pBAD_eGFP-NB_wt_H6 | Ampicillin | eGFP nanobody wt with a C-terminal His6 tag under an |

| | | |
|---|---|---|
| | | arabinose promoter |
| pBAD_eGFP-NB_R17TAG_H6 | Ampicillin | eGFP nanobody-R17TAG with a C-terminal His6 tag under an arabinose promoter |
| pBAD_eGFP-NB_R75TAG_H6 | Ampicillin | eGFP nanobody-R75TAG with a C-terminal His6 tag under an arabinose promoter |
| pBAD_GST_E51TAG_H6 | Ampicillin | GST-E51TAG with a C-terminal His6 tag under an arabinose promoter |
| pBAD_3C_Ub_K63TAG_H6 | Ampicillin | Ub-K63TAG with a N-terminal 3C cleavage site and a C-terminal His6 tag under an arabinose promoter |
| pPylT_CAT_111TAG | Tetracycline | Chloramphenicol acetyl transferase with a TAG stop codon at position 111 |
| pBK_*Mb*PylRS_lib | Ampicillin | *Mb*PylRS library with positions A276, Y271, L274, C313 and M315 randomized with NNK primers under a *gln*S promoter |
| pYOBB_Barnase_3TAG | Chloramphenicol | Barnase with a TAG stop codon at position 3 |
| pPylT_sfGFP150TAG_CAT_111TAG | Tetracycline | sfGFP with a TAG stop codon at position 150 under an arabinose promoter and Chloramphenicol |

| | | |
|---|---|---|
| | | acetyl transferase with a TAG stop codon at position 111 |
| pEVOL_*Mb*PylRS_*opp*A_wt | Chloramphenicol | *Mb*PylRS under an arabinose promotor and *opp*A wt under a *gln*S promoter |
| pEVOL_*Mb*PylRS_*opp*A_D446A | Chloramphenicol | *Mb*PylRS under an arabinose promotor and *opp*A D446A under a *gln*S promoter |
| pEVOL_*Mb*PylRS_*opp*A_R439A | Chloramphenicol | *Mb*PylRS under an arabinose promotor and *opp*A R439A under a *gln*S promoter |
| pEVOL_*Mb*PylRS_*opp*A_EP_lib | Chloramphenicol | *Mb*PylRS under an arabinose promotor and a error prone library of oppA under a glnS promoter |
| pEVOL_*Mb*PylRS_*opp*A_trimer_lib | Chloramphenicol | *Mb*PylRS under an arabinose promotor and a site saturation library of oppA under a glnS promotor |
| pEVOL_*Mb*PylRS_*opp*A_ev2 | Chloramphenicol | *Mb*PylRS under an arabinose promotor and a site saturation library of oppA ev2 under a glnS promoter |
| pBAD_oppA_wt_H6 | Ampicillin | *opp*A wt with a C-terminal His6 tag under an arabinose promoter |
| pBAD_oppA_ev2_H6 | Ampicillin | *opp*A ev2 with a C-terminal His6 tag under an |

| | | |
|---|---|---|
| | | arabinose promoter |
| pBAD_PCNA_wt_H6 | Ampicillin | PCNA wt with a C-terminal His6 tag under an arabinose promoter |
| pBAD_PCNA_K164TAG_H6 | Ampicillin | PCNA-K164TAG with a C-terminal His6 tag under an arabinose promoter |
| pBAD_β-lactamase_wt_H6 | Ampicillin | β-lactamase wt with a C-terminal His6 tag under an arabinose promoter |
| pBAD_β-lactamase_K221TAG_H6 | Ampicillin | β-lactamase-K221TAG with a C-terminal His6 tag under an arabinose promoter |
| pBAD_SUMO2_wt_H6 | Ampicillin | SUMO2 wt with a C-terminal His6 tag under an arabinose promoter |
| pBAD_SUMO_K45TAG_H6 | Ampicillin | SUMO2-K45TAG with a C-terminal His6 tag under an arabinose promoter |
| pBAD_Calmodulin_wt_strep | Ampicillin | Calmodulin wt with a C-terminal Strep tag under an arabinose promoter |
| pBAD_Calmodulin_G40TAG_strep | Ampicillin | Calmodulin-G40TAG with a C-terminal Strep tag under an arabinose promoter |
| pBAD_H3_wt_H6 | Ampicillin | Histone H3 wt with a C-terminal His6 tag under an |

| | | |
|---|---|---|
| | | arabinose promoter |
| pBAD_H3_K122TAG_H6 | Ampicillin | Histone H3-K122TAG with a C-terminal His6 tag under an arabinose promoter |
| pBAD_H3_K79TAG_K122TAG_H6 | Ampicillin | Histone H3K79TAG K122TAG with a C-terminal His6 tag under an arabinose promoter |
| pBAD_H3_K27TAG_ K79TAG_K122TAG_H6 | Ampicillin | Histone H3 K27TAG K79TAG K122TAG with a C-terminal His6 tag under an arabinose promoter |
| pBAD_Hsp82_wt_H7 | Ampicillin | Hsp82 wt with a C-terminal His7 tag under an arabinose promoter |
| pBAD_Hsp82_D452TAG_H7 | Ampicillin | Hsp82-D452TAG with a C-terminal His7 tag under an arabinose promoter |
| pET28_tyrosinase_H6 | Kanamycin | Tyrosinase with a C-terminal His6 tag under a T7 promoter |
| pSIJ8 | Ampicillin | Addgene ID: 68122 |
| pEVOL_AcKRS3(TAA)_RBS_MaPylRS_IP(TAG) | Chloramphenicol | AcKRS3 and MaPylRS_IP under a arabinose promoter polycistronically and *Mb* $tRNA_{UUA}$, *Ma* $tRNA_{CUA}$ under separate proK promoters |

| | | |
|---|---|---|
| pBAD_sfGFP_N40TAA_N150TAG_H6 | Ampicillin | sfGFP with N40TAA, N150TAG mutations and a C-terminal His6 tag under an arabinose promoter. |
| pBAD_UbK48TAA_TEV_SUMO2K11TAG_H6 | Ampicillin | Ubiquitin with K48TAA mutation followed by a TEV protease cleavage sequenced and SUMO2 with mutation K11TAG and a C-terminal His6 tag under an arabinose promoter. |

Table 2: Primers used in this study

| **Primers** | **Sequence** |
|---|---|
| OppA_EP_fwd | CGCTTTGAGGAATCCCATATGGGG |
| OppA_EP_rev | GAAACTGCAGTTAATGGTGATGATGATGGTGC |
| OppA_D221X_W222X_fwd | CGTAAGAAGACCCTTAAAA**X01X01**GTCGTAAACGAACGAATC |
| OppA_D221X_W222X_rev | CGTTAGAAGACTTTTAAGGTATAGGCACCG |
| OppA_R439X_fwd | CGAATGAAGACATGTGGCC**X01**GCAGGCTGGTGTGCTGAC |
| OppA_R439X_rev | GCATAGAAGACGGCCACATCAAAAGTACCCTGGTGAC |
| OppA_S460X_fwd | GATATGAAGACTTTCGAAC**X01**TCGATGAATACCGCGC |
| OppA_S460X_rev | GCTTAGAAGACGTTCGAAAGCATGGTGTTC |
| OppA_genomic_fwd | TACACATGCTGGTTAATACCAGTAATTATAATGAGGGAGTCCAAAAAACAATGACCAACATCACCAAGAGAAG |
| OppA_genomic_rev | AAAATCAGACACCGTGGAGCAGGACACTCCTGCCCCACGTATTGCCATTACTTCACAATGTACATATTCCGGG |

Table 3: Primers used for SLIM cloning

| **Construct** | **Primer** | **Sequence** |
|---|---|---|
| | Short fwd<br><br>Short rev | TACAACGAACCAACTTCC<br><br>ACGGGCCACATCAAAAGTAC |

| | | |
|---|---|---|
| pEVOL_*Mb* PylRS_*opp* A_D446A | Tail fwd<br>Tail rev | GCAGGCTGGTGTGCTGCTTACAACGAACCAACTTCC<br>AGCAGCACACCAGCCTGCACG |
| pEVOL_*Mb* PylRS_*opp* A_R439A | Short fwd<br>Short rev<br>Tail fwd<br>Tail rev | GCAGGCTGGTGTGCTGAC<br>CTGGTGACGGGTGTCGAG<br>GGTACTTTTGATGTGGCCGCGGCAGGCTGGTGTGCTG<br>CGCGGCCACATCAAAAGTACCCTGG |

Protein Sequences

**sfGFP N150TAG H6**
MPSKGEELFTGVVPILVELDGDVNGHKFSVRGEGEGDATNGKLTLKFICTTGKLPVP
WPTLVTTLTYGVQCFSRYPDHMKRHDFFKSAMPEGYVQERTISFKDDGTYKTRAEV
KFEGDTLVNRIELKGIDFKEDGNILGHKLEYNFNSH*VYITADKQKNGIKANFKIRHN
VEDGSVQLADHYQQNTPIGDGPVLLPDNHYLSTQSVLSKDPNEKRDHMVLLEFVTA
AGITHGMDELYKGSHHHHHH

**UbK63TAG H6**
MQIFVKTLTGKTITLEVEPSDTIENVKAKIQDKEGIPPDQQRLIFAGKQLEDGRTLSDY
NIQ*ESTLHLVLRLRGGHHHHHH

***Mb*PylRS**
MDKKPLDVLISATGLWMSRTGTLHKIKHHEVSRSKIYIEMACGDHLVVNNSRSCRTA
RAFRHHKYRKTCKRCRVSDEDINNFLTRSTESKNSVKVRVVSAPKVKKAMPKSVSR
APKPLENSVSAKASTNTSRSVPSPAKSTPNSSVPASAPAPSLTRSQLDRVEALLSPEDKI
SLNMAKPFRELEPELVTRRKNDFQRLYTNDREDYLGKLERDITKFFVDRGFLEIKSPIL
IPAEYVERMGINNDTELSKQIFRVDKNLCLRPMLAPTLYNYLRKLDRILPGPIKIFEVG
PCYRKESDGKEHLEEFTMVNFCQMGSGCTRENLEALIKEFLDYLEIDFEIVGDSCMV
YGDTLDIMHGDLELSSAVVGPVSLDREWGIDKPWIGAGFGLERLLKVMHGFKNIKR
ASRSESYYNGISTNL

***Ma*PylRS H227I Y228P**
MTVKYTDAQIQRLREYGNGTYEQKVFEDLASRDAAFSKEMSVASTDNEKKIKGMIA
NPSRHGLTQLMNDIADALVAEGFIEVRTPIFISKDALARMTITEDKPLFKQVFWIDEKR
ALRPMLAPNLYSVMRDLRDHTDGPVKIFEMGSCFRKESHSGMHLEEFTMLNLVDMG
PRGDATEVLKNYISVVMKAAGLPDYDLVQEESDVYKETIDVEINGQEVCSAAVGPIPL
DAAHDVHEPWSGAGFGLERLLTIREKYSTVKKGGASISYLNGAKIN

**DiazKRS**
MDKKPLNTLISATGLWMSRTGTIHKIKHHEVSRSKIYIEMACGDHLVVNNSRSSRTAR
ALRHHKYRKTCKRCRVSDEDLNKFLTKANEDQTSVKVKVVSAPTRTKKAMPKSVAR
APKPLENTEAAQAQPSGSKFSPAIPVSTQESVSVPASVSTSISSISTGATASALVKGNTN
PITSMSAPVQASAPALTKSQTDRLEVLLNPKDEISLNSGKPFRELESELLSRRKKDLQQ
IYAEERENYLGKLEREITRFFVDRGFLEIKSPILIPLEYIERMGIDNDTELSKQIFRVDKN

FCLRPMLAPNLMNYARKLDRALPDPIKIFEIGPCYRKESDGKEHLEEFTMLNFAQMGS
GCTRENLESIITDFLNHLGIDFKIVGDSCMVYGDTLDVMHGDLELSSAVVGPIPLDRE
WGIDKPWIGAGFGLERLLKVKHDFKNIKRAARSESYYNGISTNL

**eGFP-NB wt**
MKYLLPTAAAGLLLLAAQPAMAQVQLVESGGALVQPGGSLRLSCAASGFPVNRYSM
RWYRQAPGKEREWVAGMSSAGDRSSYEDSVKGRFTISRDDARNTVYLQMNSLKPE
DTAVYYCNVNVGFEYWGQGTQVTVSSKKKHHHHHH

**eGFP-NB R17TAG H6**
MKYLLPTAAAGLLLLAAQPAMAQVQLVESGGALVQPGGSL*LSCAASGFPVNRYSM
RWYRQAPGKEREWVAGMSSAGDRSSYEDSVKGRFTISRDDARNTVYLQMNSLKPE
DTAVYYCNVNVGFEYWGQGTQVTVSSKKKHHHHHH

**eGFP-NB R75TAG H6**
MKYLLPTAAAGLLLLAAQPAMAQVQLVESGGALVQPGGSLRLSCAASGFPVNRYSM
RWYRQAPGKEREWVAGMSSAGDRSSYEDSVKGRFTISRDDA*NTVYLQMNSLKPED
TAVYYCNVNVGFEYWGQGTQVTVSSKKKHHHHHH

**GST E51TAG H6**
MSPILGYWKIKGLVQPTRLLLEYLEEKYEEHLYERDEGDKWRNKKFELGL*FPNLPY
YIDGDVKLTQSMAIIRYIADKHNMLGGCPKERAEISMLEGAVLDIRYGVSRIAYSKDF
ETLKVDFLSKLPEMLKMFEDRLCHKTYLNGDHVTHPDFMLYDALDVVLYMDPMCL
DAFPKLVCFKKRIEAIPQIDKYLKSSKYIAWPLQGWQATFGGGDHPPKHHHHHH

**3C-Ub-K63TAG H6**
MENLLEVLFQGPGGGGSMQIFVKTLTGKTITLEVEPSDTIENVKAKIQDKEGIPPDQQ
RLIFAGKQLEDGRTLSDYNIQ*ESTLHLVLRLRGGHHHHHH

**OppA H6**
MGTNITKRSLVAAGVLAALMAGNVALAADVPAGVTLAEKQTLVRNNGSEVQSLDPH
KIEGVPESNISRDLFEGLLVSDLDGHPAPGVAESWDNKDAKVWTFHLRKDAKWSDG
TPVTAQDFVYSWQRSVDPNTASPYASYLQYGHIAGIDEILEGKKPITDLGVKAIDDHT
LEVTLSEPVPYFYKLLVHPSTSPVPKAAIEKFGEKWTQPGNIVTNGAYTLKDWVVNE
RIVLERSPTYWNNAKTVINQVTYLPIASEVTDVNRYRSGEIDMTNNSMPIELFQKLKK
EIPDEVHVDPYLCTYYYEINNQKPPFNDVRVRTALKLGMDRDIIVNKVKAQGNMPAY
GYTPPYTDGAKLTQPEWFGWSQEKRNEEAKKLLAEAGYTADKPLTINLLYNTSDLH
KKLAIAASSLWKKNIGVNVKLVNQEWKTFLDTRHQGTFDVARAGWCADYNEPTSFL
NTMLSNSSMNTAHYKSPAFDSIMAETLKVTDEAQRTALYTKAEQQLDKDSAIVPVYY
YVNARLVKPWVGGYTGKDPLDNTYTRNMYIVKHHHHHH

**OppA ev2 H6**
MGTNITKRSLVAAGVLAALMAGNVALAADVPAGVTLAEKQTLVRNNGSEVQSLDPH
KIEGVPESNISRDLFEGLLVSDLDGHPAPGVAESWDNKDAKVWTFHLRKDAKWSDG
TPVTAQDFVYSWQRSVDPNTASPYASYLQYGHIAGIDEILEGKKPITDLGVKAIDDHT
LEVTLSEPVPYFYKLLVHPSTSPAPKAAIEKFGEKWTQPGNIVTNGAYTLKLAVVNERI
VLERSPTYWNNAKTVINQVTYLPIASEVTDVNRYRSGEIDMTNNSMPIELFQKLKKEI
PDEVHVDPYLCTYYYEVNNQKPPFNDVRVRTALKLGMDRDIIVNKVKAQGDMPAYG
YTPPYTDGAKLTQPEWFGWSQEKRNEEAKKLLAEAGYTADKPLTINLLYNTSDLHK
KLAIAASSLWKKNIGVNVKLVNQEWKTFLDTRHQGTFDVAQAGWCADYNEPTSFLN

TMLSNHSMNTAHYKSPAFDSIMAETLKVTDEAQRTALYTKAEQQLDKDSAIVPVYY
YVNARLVKPWVGGYTGKDPLDNTYTRNMYIVKHHHHHH

**β-lactamase wt H6**
MGADLADRFAELERRYDARLGVYVPATGTTAAIEYRADERFAFCSTFKAPLVAAVLH
QNPLTHLDKLITYTSDDIRSISPVAQQHVQTGMTIGQLCDAAIRYSDGTAANLLLADL
GGPGGGTAAFTGYLRSLGDTVSRLDAEEPELNRDPPGDERDTTTPHAIALVLQQLVL
GNALPPDKRALLTDWMARNTTGAKRIRAGFPADWKVIDKTGTGDYGRANDIAVVW
SPTGVPYVVAVMSDRAGGGYDAEPREALLAEAATCVAGVLAGSGGSGHHHHHH

**β-lactamase K221TAG H6**
MGADLADRFAELERRYDARLGVYVPATGTTAAIEYRADERFAFCSTFKAPLVAAVLH
QNPLTHLDKLITYTSDDIRSISPVAQQHVQTGMTIGQLCDAAIRYSDGTAANLLLADL
GGPGGGTAAFTGYLRSLGDTVSRLDAEEPELNRDPPGDERDTTTPHAIALVLQQLVL
GNALPPDKRALLTDWMARNTTGA*RIRAGFPADWKVIDKTGTGDYGRANDIAVVWS
PTGVPYVVAVMSDRAGGGYDAEPREALLAEAATCVAGVLAGSGGSGHHHHHH

**SUMO2 wt H6**
MADEKPKEGVKTENNDHINLKVAGQDGSVVQFKIKRHTPLSKLMKAYCERQGLSM
RQIRFRFDGQPINETDTPAQLEMEDEDTIDVFQQQTGGHHHHHH

**SUMO2 K45TAG H6**
MADEKPKEGVKTENNDHINLKVAGQDGSVVQFKIKRHTPLSKLM*AYCERQGLSMR
QIRFRFDGQPINETDTPAQLEMEDEDTIDVFQQQTGGHHHHHH

**Calmodulin wt strep**
MADQLTEEQIAEFKEAFSLFDKDGDGTITTKELGTVMRSLGQNPTEAELQDMINEVD
ADGNGTIDFPEFLTMMARKMKDTDSEEEIREAFRVFDKDGNGYISAAELRHVMTNL
GEKLTDEEVDEMIREADIDGDGQVNYEEFVQMMTAKSAWSHPQFEKGGGSGGGSG
GSAWSHPQFEK

**Calmodulin G40TAG strep**
MADQLTEEQIAEFKEAFSLFDKDGDGTITTKELGTVMRSL*QNPTEAELQDMINEVD
ADGNGTIDFPEFLTMMARKMKDTDSEEEIREAFRVFDKDGNGYISAAELRHVMTNL
GEKLTDEEVDEMIREADIDGDGQVNYEEFVQMMTAKSAWSHPQFEKGGGSGGGSG
GSAWSHPQFEK

**Histone H3 wt H6**
MARTKQTARKSTGGKAPRKQLATKAARKSAPATGGVKKPHRYRPGTVALREIRRYQ
KSTELLIRKLPFQRLVREIAQDFKTDLRFQSSAVMALQEASEAYLVALFEDTNLCAIHA
KRVTIMPKDIQLARRIRGERARSHHHHHH

**Histone H3 K112TAG H6**
MARTKQTARKSTGGKAPRKQLATKAARKSAPATGGVKKPHRYRPGTVALREIRRYQ
KSTELLIRKLPFQRLVREIAQDFKTDLRFQSSAVMALQEASEAYLVALFEDTNLCAIHA
KRVTIMP*DIQLARRIRGERARSHHHHHH

**Histone H3 K79 K112TAG H6**

MARTKQTARKSTGGKAPRKQLATKAARKSAPATGGVKKPHRYRPGTVALREIRRYQ
KSTELLIRKLPFQRLVREIAQDF*TDLRFQSSAVMALQEASEAYLVALFEDTNLCAIHA
KRVTIMP*DIQLARRIRGERARSHHHHHH

**Histone H3 K27 K79 K112TAG H6**
MARTKQTARKSTGGKAPRKQLATKAAR*SAPATGGVKKPHRYRPGTVALREIRRYQK
STELLIRKLPFQRLVREIAQDF*TDLRFQSSAVMALQEASEAYLVALFEDTNLCAIHAK
RVTIMP*DIQLARRIRGERARSHHHHHH

**Tyrosinase H6**
MSNKYRVRKNVLHLTDTEKRDFVRTVLILKEKGIYDRYIAWHGAAGKFHTPPGSDRN
AAHMSSAFLPWHREYLLRFERDLQSINPEVTLPYWEWETDAQMQDPSQSQIWSADF
MGGNGNPIKDFIVDTGPFAAGRWTTIDEQGNPSGGLKRNFGATKEAPTLPTRDDVLN
ALKITQYDTPPWDMTSQNSFRNQLEGFINGPQLHNRVHRWVGGQMGVVPTAPNDPV
FFLHHANVDRIWAVWQIIHRNQNYQPMKNGPFGQNFRDPMYPWNTTPEDVMNHRK
LGYVYDIELRKSKRSSLEHHHHHH

**Hsp82 H7**
MASETFEFQAEITQLMSLIINTVYSNKEIFLRELISNASDALDKIRYKSLSDPKQLETEP
DLFIRITPKPEQKVLEIRDSGIGMTKAELINNLGTIAKSGTKAFMEALSAGADVSMIGQ
FGVGFYSLFLVADRVQVISKSNDDEQYIWESNAGGSFTVTLDEVNERIGRGTILRLFL
KDDQLEYLEEKRIKEVIKRHSEFVAYPIQLVVTKEVEKEVPIPEEEKKDEEKKDEEKK
DEDDKKPKLEEVDEEEEKKPKTKKVKEEVQEIEELNKTKPLWTRNPSDITQEEYNAF
YKSISNDWEDPLYVKHFSVEGQLEFRAILFIPKRAPFDLFESKKKKNNIKLYVRRVFIT
DEAEDLIPEWLSFVKGVVDSEDLPLNLSREMLQQNKIMKVIRKNIVKKLIEAFNEIAE
DSEQFEKFYSAFSKNIKLGVHEDTQNRAALAKLLRYNSTKSVDELTSLTDYVTRMPE
HQKNIYYITGESLKAVEKSPFLDALKAKNFEVLFLTDPIDEYAFTQLKEFEGKTLVDIT
KDFELEETDEEKAEREKEIKEYEPLTKALKEILGDQVEKVVVSYKLLDAPAAIRTGQF
GWSANMERIMKAQALRDSSMSSYMSSKKTFEISPKSPIIKELKKRVDEGGAQDKTVK
DLTKLLYETALLTSGFSLDEPTSFASRINRLISLGLNIDEDEETETAPEASTAAPVEEVPA
DTEMEEVDPGEQKCEEWKRRYEKEKEKNARLKGKVEKLEIELARWRPGSAWSHHH
HHHH

**Hsp82 D452TAG H7**
MASETFEFQAEITQLMSLIINTVYSNKEIFLRELISNASDALDKIRYKSLSDPKQLETEP
CLFIRITPKPEQKVLEIRDSGIGMTKAELINNLGTIAKSGTKAFMEALSAGADVSMIGQ
FGVGFYSLFLVADRVQVISKSNDDEQYIWESNAGGSFTVTLDEVNERIGRGTILRLFL
KDDQLEYLEEKRIKEVIKRHSEFVAYPIQLVVTKEVEKEVPIPEEEKKDEEKKDEEKK
DEDDKKPKLEEVDEEEEKKPKTKKVKEEVQEIEELNKTKPLWTRNPSDITQEEYNAF
YKSISNDWEDPLYVKHFSVEGQLEFRAILFIPKRAPFDLFESKKKKNNIKLYVRRVFIT
DEAEDLIPEWLSFVKGVVDSEDLPLNLSREMLQQNKIMKVIRKNIVKKLIEAFNEIAE
DSEQFEKFYSAFSKNIKLGVHEDTQNRAALAKLLRYNSTKSV*ELTSLTDYVTRMPEH
QKNIYYITGESLKAVEKSPFLDALKAKNFEVLFLTDPIDEYAFTQLKEFEGKTLVDITK
DFELEETDEEKAEREKEIKEYEPLTKALKEILGDQVEKVVVSYKLLDAPAAIRTGQFG
WSANMERIMKAQALRDSSMSSYMSSKKTFEISPKSPIIKELKKRVDEGGAQDKTVKD
LTKLLYETALLTSGFSLDEPTSFASRINRLISLGLNIDEDEETETAPEASTAAPVEEVPAD
TEMEEVDPGEQKCEEWKRRYEKEKEKNARLKGKVEKLEIELARWRPGSA
WSHHHHHHH

**AcKRS3**

MDKKPLDVLISATGLWMSRTGTLHKIKHHEVSRSKIYIEMACGDHLVVNNSRSCRTA
RAFRHHKYRKTCKRCRVSGEDINNFLTRSTESKNSVKVRVVSAPKVKKAMPKSVSR
APKPLENSVGAKASTNTSRSVPSPAKSTPNSSVPASAPAPSLTRSQLDRVEALLSPEDKI
SLNMAKPFRELEPELVTRRKNDFQRLYTNDREDYLGKLERDITKFFVDRGFLEIKSPIL
IPAEYVERMGINNDTELSKQIFRVDKNLCLRPMMAPTIFNYARKLDRILPGPIKIFEVG
PCYRKESDGKEHLEEFTMVNFFQMGSGCTRENLEALIKEFLDYLEIDFEIVGDSCMV
YGDTLDIMHGDLELSSAVVGPVSLDREWGIDKPWIGAGFGLERLLKVMHGFKNIKR
ASRSESYYNGISTNL

**sfGFP N40TAA N150TAG H6**
MPSKGEELFTGVVPILVELDGDVNGHKFSVRGEGEGDAT*GKLTLKFICTTGKLPVPW
PTLVTTLTYGVQCFSRYPDHMKRHDFFKSAMPEGYVQERTISFKDDGTYKTRAEVKF
EGDTLVNRIELKGIDFKEDGNILGHKLEYNFNSH*VYITADKQKNGIKANFKIRHNVE
DGSVQLADHYQQNTPIGDGPVLLPDNHYLSTQSVLSKDPNEKRDHMVLLEFVTAAGI
THGMDELYKGSHHHHHH

**Ub K48TAA TEV SUMO2K11TAG H6**
MQIFVKTLTGKTITLEVEPSDTIENVKAKIQDKEGIPPDQQRLIFAG*QLEDGRTLSDY
NIQKESTLHLVLRLRGGEDLYFQSMADEKPKEGVKTENNDHINL*VAGQDGSVVQFK
IKRHTPLSKLMKAYCERQGLSMRQIRFRFDGQPINETDTPAQLEMEDEDTIDVFQQQQ
NGLHHHHHHH

**Synthesis of peptides via solid phase peptide synthesis**

All peptides were synthesized via solid phase peptide synthesis (SPPS). Reactions were performed in plastic syringes with a frit using <1g of resin. SPPS was done using the Fmoc-strategy using a 2-Chlorotrityl chloride resin. For charging, 2 eq. of Fmoc protected amino acid and 3 eq of DIPEA were dissolved in DCM and added to 1eq of resin (1.5 mmol/g maximum capacity, 100-200 mesh) and incubated for 1 h on a roller at RT. Resin was then washed 5 times with DCM and 5 times with DMF. Fmoc deprotection was performed by adding a 20% piperidine (v/v) solution in DMF and incubated for 10 min at RT. This process was performed twice for complete deprotection followed by washing the resin 5 times with DMF. For subsequent coupling of amino acids a coupling solution was prepared by dissolving 2 eq. of Fmoc protected amino acid, 1.8 eq. HATU and 3 eq. DIPEA in DMF and mixing for 5 min at RT. Coupling solution was added to the resin and incubated on a roller for 1 h at RT. This process was repeated depending on the length of peptide being synthesized.

For cleavage of the protected peptide off the resin, the resin was washed 5 times with DMF and 5 times with DCM and cleavage was performed by adding 20% HFIP in DCM (v/v) for 10 minutes at RT. This process was repeated 2 times, each time collecting the filtrate. Solvent from the filtrate was evaporated under reduced pressure.

Deprotection was performed by adding 95% TFA with 2.5% water and 2.5% TIPS and stirred at RT till deprotection was complete, monitoring the reaction via LC-MS. Deprotected peptide was subsequently precipitated in cold ether and dissolved in water. Any remaining ether was evaporated under low pressure and the resulting peptide solution was flash frozen in liquid nitrogen and lyophilized to obtain a dry powder.

Table 4: Peptides used in this study

| **Peptide** | **Expected mass** | **[M+H]+** |
|---|---|---|
| AisoK | 217.25 | 218.2 |
| G-AisoK | 274.32 | 275.2 |
| SisoK | 233.25 | 234.2 |
| G-SisoK | 290.32 | 291.1 |
| TisoK | 247.28 | 248.2 |
| G-TisoK | 304.35 | 305.2 |
| VisoK | 245.31 | 246.2 |
| G-VisoK | 302.38 | 303.2 |
| CisoK | 249.31 | 250.2 |
| G-CisoK | 306.38 | 307.2 |
| PisoK | 243.29 | 244.2 |
| G-PisoK | 300.36 | 301.2 |
| PrgisoK | 241.27 | 242.2 |
| G-PrgisoK | 298.34 | 299.1 |
| LisoK | 259.33 | 260.2 |
| G-LisoK | 316.4 | 317.2 |
| pLisoK | 271.3 | 272.2 |

| | | |
|---|---|---|
| G-pLisoK | 328.37 | 329.2 |
| HisoK | 283.31 | 284.2 |
| G-HisoK | 340.38 | 341.2 |
| K-SisoK | 361.44 | 362.2 |
| AcK-pLisoK | 441.53 | 442.3 |

**Protein purification**

Expression and purification of OppA and variants

Chemically competent *E. coli* K12 Δ*Opp*A cells were transformed with pBAD_OppA_His6. After recovery in 1 mL SOC for 1 h at 37 °C, the cells were cultured in 50 mL of 2-YT supplemented with 100 µg/mL Ampicillin overnight at 37 °C. the overnight culture was then diluted to $OD_{600}$ 0.05 in 2-YT supplemented with 50 µg/mL Ampicillin and grown at 37 °C, 200 rpm till $OD_{600}$ 0.6. Protein expression was induced with 0.05% L-arabinose and the culture grown for 3 h at 37 °C, 200 rpm. Cells were harvested by centrifugation at 7000 x g for 20 min and pellets flash frozen in liquid nitrogen and stored at -20 °C.

Cell pellets were resuspended in lysis buffer (20 mM Tris-HCl pH 8, 300 mM NaCl, 30 mM imidazole, 1 mM PMSF) (30 ml/liter culture) and cells lysed using a cell disruptor (CF1 Constant Systems Ltd., 40 Kpsi) . Lysed cells were centrifuged at 14,000 x g, 20 minutes and cleared lysate loaded on a 5ml HisTrap HP column (Cytiva) pre-equilibrated with wash buffer (20 mM Tris-HCl pH 8, 300 mM NaCl, 30 mM imidazole). The loaded column was then washed with 5 column volumes (CV) of wash buffer. Protein was eluted with a gradient from 30 to 300 mM imidazole over 20 CV, collecting 1.4 ml fractions. Fractions containing protein were pooled and concentrated using Amicon® centrifugal filter units (Millipore, 30 kDa MWCO) and then injected on a HiLoad 16/600 Superdex 75 pg size exclusion (SE) column (Cytiva) equilibrated with SE buffer (2x PBS). SE fractions containing protein were then pooled and concentrated. Any endogenously bound peptides were removed by partially unfolding OppA with 2M Guanidine hydrochloride (GdnHCl) in SE buffer and removing the peptides via dialysis (Pur-A-Lyzer™ Maxi, 12 kDa MWCO, Sigma-Aldrich) in SE buffer containing decreasing amounts of GdnHCl from 2 M to 0 M. resulting protein was then flash frozen in liquid nitrogen and stored at -80 °C till further use.

Expression of XisoK bearing proteins

Chemically competent *E. coli* K12 cells were co-transformed with pBAD_POI (encodes the protein of interest (POI) with a C-terminal H6 tag) and pEVOL_PylRS (encodes two copies of the PylRS variant and $tRNA_{CUA}$). After recovery in 1 mL SOC media for 1 h at 37 °C, cells were cultured in 5 ml of 2-YT supplemented with Ampicillin (100 µg/mL) and Chloramphenicol (50 µg/mL) and grown overnight at 37 °C. The overnight culture was then diluted to an $OD_{600}$ of 0.05 in either 2-YT or Autoinducing media[13] supplemented with Ampicillin (100 µg/mL), Chloramphenicol (50 µg/mL) and non-canonical amino acid. Cells grown in 2-YT were grown till an $OD_{600}$ of 0.6 and induced with 0.05% L-arabinose and grown overnight at 37 °C. Cells grown in Autoinducing media were directly grown overnight at 37

°C. Cells of the Overnight culture were harvested by centrifugation at 4000 x g for 10 min and the pellets stored at -20 °C till further use.

Purification of His6 tagged proteins

Cell pellets were resuspended in lysis buffer (20 mM Tris-HCl pH 8, 300 mM NaCl, 30 mM imidazole, 1 mM PMSF) and the cells lysed via sonication in an ice water bath. Lysed cells were centrifuged at 14,000 x g for 20 minutes and cleared lysate was incubated with Ni Sepharose fast flow beads (1 mL slurry/ 1 L culture, Cytiva) pre-equilibrated with Ni-NTA wash buffer (20 mM Tris-HCl pH 8, 300 mM NaCl, 30 mM imidazole) and incubated on a roller for 1 h at 4 °C. Beads were then washed with 10 CV of wash buffer 3 times and protein eluted with elution buffer (20 mM Tris-HCl pH 8, 300 mM NaCl, 300 mM imidazole). Resulting protein was either directly used for mass determination or further purification was performed via gel filtration using a Superdex 75 increase 10/300 GL column (Cytiva) equilibrated with SE buffer (2xPBS pH 7 for eGFP-NB or 20 mM Potassium phosphate buffer pH 6.5, 100 mM NaCl for PisoK bearing proteins or 1x PBS pH 7.4 for Affibody and ProteinZ). Fractions containing protein were pooled and concentrated using Amicon® centrifugal filter units with the appropriate MWCO. Protein was then flash frozen in liquid nitrogen and stored at -80 °C till further use. Proteins bearing CisoK were treated with methoxyamine (10 µM protein, 100 mM methoxyamine at pH 4 ammonium acetate) and buffer exchanged in 1xPBS before storage.

Expression and purification of Tyrosinase

Chemically competent *E. coli* BL21 (DE3) were transformed with pET28_Tyrosinase_H6 (encoding for tyrosinase from *Priesta megaterium*). After recovery with 1 mL of SOC medium for 1 h at 37 °C, the cells were cultured overnight in 50 mL 2-YT medium containing kanamycin (50 µg/mL) at 37 °C, 200 rpm. The overnight culture was diluted to an $OD_{600}$ of 0.05 in 1 L of fresh 2-YT medium supplemented with kanamycin (25 µg/mL) and cultured at 37 °C with shaking (200 rpm) until $OD_{600}$ reached 0.5.

IPTG was added to a final concentration of 0.4 mM and protein expression was induced for 18 h at 37 °C. The cells were harvested by centrifugation (4000 × g, 20 min, 4 °C), resuspended in lysis buffer (20 mM Tris-HCl pH 8, 300 mM NaCl, 30 mM imidazole, 0.2 mM PMSF) and lysed via sonication in an ice water bath. The lysed cells were centrifuged (15,000 × g, 40 min, 4 °C) and the cleared lysate was applied to Ni Sepharose fast flow beads (1mL slurry/ 1L culture) pre-equilibrated with Ni-NTA wash buffer (20 mM Tris-HCl pH 8, 300 mM NaCl, 30 mM imidazole) and incubated on a roller for 1 h at 4 °C. After incubation, the mixture was transferred to an empty plastic column and washed with 10 CV of wash buffer (20 mM Tris-HCl pH 8, 300 mM NaCl, 30 mM imidazole). Tyrosinase was eluted in 1 mL fractions with wash buffer supplemented with 300 mM imidazole and 0.02 mM $CuSO_4$. The fractions containing the protein were pooled and buffer was exchanged to PBS pH 7.4 supplemented with 15 % glycerol and 0.02 mM CuSO4 using an Amicon® centrifugal filter units with an appropriate MWCO (Millipore). Protein concentration was calculated from the measured A280 absorption (extinction coefficients were calculated with ProtParam (https://web.expasy.org/protparam/). Tyrosinase was flash frozen using liquid nitrogen and stored at -80 °C until further use.

eGFP nanobody purification and yield determination

In order to determine yields of amber suppressed eGFPNb, expression and purification were performed as with other XisoK bearing proteins with a few modifications. Both *E. coli* K12 and isoK12 were grown in 25 mL 2xYT cultures manually inducing expression with 0.05% arabinose at an $OD_{600}$ of 0.6. 300 uL of Ni-NTA bead slurry was used for each purification and after loading the beads with lysate an additional high salt wash was done with wash buffer containing 1 M NaCl to remove non-specifically bound proteins. Protein was eluted in elution buffer till no more protein was detected in the flow-though, checking with Bradford's reagent. Eluted protein was buffer exchanged in 1xPBS, 5 times and concentrations determined using a NanoPhotometer® NP60 (Implen). To check protein purity as well as if all the protein bound to the beads, eluted protein and lysate flow-though were analyzed via SDS-PAGE.

**Evolution of HisoKRS**

Electrocompetent *E. coli* DH10β cells containing a positive selection plasmid pPylT_CAT111TAG were transformed with a pBK_*Mb*PylRS_lib (MbPylRS library with positions A276, Y271, L274, C313 and M315 randomized using degenerate NNK primers). After recovery with 5 mL SOC for 1 hour at 37 °C. cells were added to 100 mL of 2-YT supplemented with Ampicillin (100 µg/mL) and tetracycline (17 µg/ml) and grown to an $OD_{600}$ of 0.6. Cells were then plated in a dilution series on 24 cm autoinduction agar plates containing 0.5 mM G-HisoK and incubated overnight at 37 °C. Plates with distict single colonies were scraped and the plasmids were isolated from the cell mass and the pBK_*Mb*PylRS_lib was isolated on a 1% agarose gel. Electrocompetent *E. coli* DH10β cells containing a negative selection plasmid pYOBB_Barnase3TAG were then transformed with the isolated library and after recovery with 1 mL SOB for 1 h at 37 °C, cells were added to 100 ml of 2-YT supplemented with Ampicillin (100 µg/mL) and Chloramphenicol (50 µg/mL) and grown to an $OD_{600}$ of 0.6. For negative selection, cells were then plated on LB-agar plates supplemented with Ampicillin (100 µg/mL), Chloramphenicol (50 µg/mL) and 0.2% arabinose and grown overnight at 37 °C. Plates were scraped and library plasmid isolated for a final positive selection round using a pPylT_GFP_N150TAG_CAT111TAG double reporter plasmid. Single colonies were then picked into a 96 deep well plate containing 1 mL non-inducing media/well and grown overnight at 37 °C. Two 96 deep well plates containing autoinducing media with and without 0.5 mM G-HisoK were inoculated with the overnight culture and grown for 24 h. sfGFP fluorescence was measured for both plates on a Varioscan Lux plate reader (Thermo Scientific). Clones with sufficient fluorescence above negative sample were sent for sanger sequencing.

**Generation of a OppA error prone library**

An error prone library of *opp*A was generated using an error prone PCR kit (Jena Bioscience Cat no. PP-102). Primers OppA_EP_fwd and OppA_EP_rev (**Table 2**) were used to amplify *opp*A from pEVOL_*Mb*PylRS_*opp*A according to manufacturer's instructions running for 25 cycles and adding 2.5 µL error prone solution. Resultant amplicon was purified on a 1% agarose gel and digested with *Nde*1 and *Pst*1-HF. Digested insert was then ligated with T4 ligase into a backbone generated by digesting pEVOL_*Mb*PylRS_*opp*A with *Nde*1 and *Pst*1-HF followed by dephosphorylation with Antarctic Phosphatase. Electrocompetent DH10β cells were then transformed with the purified ligation. After recovery in SOC media for 1 h at 37 °C, cells were added to 50 ml of 2-YT supplemented with Chloramphenicol (50 µg/mL) and grown overnight at 37 °C. library plasmid from the overnight culture was purified and used for subsequent steps. The size of the library was determined by plating the freshly transformed cells in a dilution series on LB-agar plates supplemented with Chloramphenicol (50 µg/mL). The size was calculated to be 4 x $10^6$. Multiple single clones were sequenced to determine the average error rate and was optimized to be ~ 0.8% or an average of 5 amino acid mutations/gene.

**Generation of an OppA site saturation library**

4 hotspot positions in OppA were selected for site saturation based on sequenced variants from selection of the error prone library. Positions were chosen if mutations occurred more than once (R439, S460) or if mutations occurred close to each other in different variants (D221, W222).

Sites were randomized using degenerate trimer primers (Ella Biotech). A mix of templates based on pEVOL_*Mb*PylRS_*opp*A containing OppA wt and four variants from the error prone library were amplified with library primers (see primer table.). The linear amplicon was then purified on a 1% agarose gel and digested with *Bbs*1-HF and *Dpn*1. Digested fragment was circularized via ligation with T4 ligase and electrocompetent DH10β cells were transformed with the circularized library plasmid. After recovery in SOC media for 1 h at 37 °C, cells were added to 50 ml of 2-YT supplemented with Chloramphenicol (50 µg/mL) and grown overnight at 37 °C. Library plasmid was purified from the overnight culture and used as the template for the next round of amplification to randomize the next position(s). This process was repeated 3 times to give a library with positions D221, W222, R439 and S460 mutated to all 20 amino acids on oppA wt and 4 error prone variants. The library size was 8 x $10^5$.

**FACS based screening protocol**

Electrocompetent *E. coli* K12 Δ*Opp*A cells containing sfGFP reporter plasmid pBAD_sfGFP_N150TAG_H6 were transformed with the OppA library (pEVOL_*Mb*PylRS_*opp*A_EP_lib, pEVOL_*Mb*PylRS_*opp*A_trimer_lib or pEVOL_*Mb*PylRS_*opp*A_wt). After recovery in 1 mL SOC for 1 h at 37 °C, transformed cells were diluted in 50 mL of non-inducing media (auto-inducing media without arabinose) supplemented with Ampicillin (100 µg/mL) and Chloramphenicol (50 µg/mL) and grown overnight. The Overnight culture was then diluted in non-inducing media and grown to an $OD_{600}$ of 0.6. At this point, 0.05% arabinose was added to induce sfGFP expression and the culture was split into smaller cultures supplemented with +/- 0.5 mM G-SisoK and varying amounts of tryptone to apply a selection pressure towards OppA variants which preferably bound to G-SisoK. Cultures were then grown for 4 h at 37 °C to allow for sfGFP expression. To halt growth, the cultures were cooled on ice for 10 minutes, centrifuged (4000 x g, 5 min) and resuspended in ice cold PBS. The PBS cell suspension was sorted on a Sony cell sorter (SH800) using a 70 µM chip sorting for cells with highest sfGFP fluorescence intensity. Gating was decided based on the positive control (oppA wt, 0.5 mM G-SisoK, 0 g/L tryptone) and the negative controls (oppA wt, 0.5 mM G-SisoK, Xg/L tryptone – with X being the tryptone concentration used in that round of enrichment, and oppA wt, 0 mM G-SisoK, 0 g/L tryptone). For each round cells with the top 0.5-2% sfGFP fluorescence were sorted. Sorted cells were recovered in SOC media supplemented with Ampicillin (50 µg/mL) and Chloramphenicol (25 µg/mL) overnight at 37 °C and the process repeated for further enrichment. After multiple rounds of enrichment, cells were sorted into a 96-well plate containing SOC with supplemented with Ampicillin (50 µg/mL) and Chloramphenicol (25 µg/mL) and grown overnight at 37 °C. Cultures grown from single cells were further evaluated via the fluorescence plate reader assay and variants which showed the desired phenotype were sequenced.

The error prone library was subjected to 5 rounds of enrichment, each time doubling the tryptone concentration from 1 g/L (round 1) to 16 g/L (round 5).

The site saturation library was directly grown in LB media (10 g/L tryptone) for 2 rounds of enrichment and 2xYT (16 g/L) for 3 rounds of enrichment.

Table 5: Mutations in OppA variants

| OppA variant | Mutations |
|---|---|
| Error-prone Variant 1 | T173N, **D221**G, K371E, **R439**H |
| Error-prone Variant 2 | L78I, **W222**R, K307R, K333N |
| Error-prone Variant 3 | T429A, **R439**L, **S460**C, V482A |
| Error-prone Variant 4 | V193A, I303V, N337D, **S460**N |
| Site-saturation variant OppA-ev2 | V193A, **D221**L, **W222**A, I303V, N337D, **R439**Q, **S460**H |

In **bold**: positions targeted for site saturation mutagenesis

**Preparation of E. coli lysates for LC-MS based uptake assays**

Uptake assay protocol adapted from previously published protocol[14]. Relevant *E. coli* strains were transformed with a dummy pBAD plasmid to prevent contamination of cultures. After recovery in 1mL SOC for 1 h at 37 °C, cells were cultured in 5 ml of 2xYT or Autoinducing media supplemented with Ampicillin (100 µg/mL) and grown overnight at 37 °C. Overnight cultures were diluted to an $OD_{600}$ of 0.05 in 5 mL of either 2xYT or autoinducing media supplemented with Ampicillin (100 µg/mL) and non-canonical amino acid and grown overnight at 37 °C. The $OD_{600}$ of the overnight cultures was determined and 12 OD.mL were harvested by centrifugation at 4000 x g for 10 minutes. Cell pellets were then washed 3 times with 1 mL of cold media and resuspended in 400 µL of a methanol:water solution (60:40). Cells were lysed via 5 freeze thaw cycles in liquid nitrogen and a 42 °C water bath. Lysate was cleared by centrifugation at 17,900 x g for 20 min. 5 uL of cleared lysate was injected onto the LC-MS for analysis. For cultures incubated with BocK, samples were injected onto a Zorbax

SB-C18 (Agilent, 4.6 x 150 mm) column and a gradient of 5-95% was used. For cultures incubated with XisoK and GXisoK peptides, samples were injected on a Poroshell 120 HILIC-Z (Agilent, 2.1 x 100 mm) column and using a gradient of 95-10% . The mass spectrometer was set to single ion mode to detect the relevant m/z for each non-canonical amino acid.

To determine intracellular concentrations, calibration points of lysate spiked with known non-canonical amino acid concentrations were measured. Ion peaks were integrated, and integral values plotted against concentration to determine a linear calibration line. Integral values of unknown samples interpolated on calibration line to determine lysate concentrations. Intracellular concentrations were estimated assuming 1 $OD_{600}$ = 8 x $10^8$ cells/mL and the volume of an *E. coli* cell = 0.6 fL.

**Determination of $K_D$s using microscale thermophoresis**

Microscale thermophoresis (MST) was performed on the NanoTemper Monolith NT.115 (NanoTemper Technologies). Wt OppA and evolved variants were fluorescently labeled using the Monolith Protein Labeling Kit RED-NHS 2nd Generation (NanoTemper Technologies) and diluted to 100 nM in assay buffer (2xPBS, 0.02% tween 20). Peptides were diluted to double the highest measured concentration in assay buffer and diluted 2-fold in a dilution series to give 16 peptide concentrations. Equal volumes of protein and peptide solutions were mixed (final protein concentration of 50 nM) and incubated at room temperature for 30 minutes. Samples were loaded into capillaries (Monolith NT.115 Capillaries, NanoTemper Technologies) and measured according to manufacturer's instructions. Three independent replicates were measured for each peptide-protein combination and data analysis was performed with MO.affinity Analysis (v3.0.5, NanoTemper Technologies).

**Generating isoK12 strain via homologous recombination**

Primers with 50 bp overhangs homologous to regions upstream and downstream of the OppA locus in *E. coli* genome were used to amplify *opp*A ev2 from pEVOL_*Mb*PylRS_*opp*A_ev2. (**Table 2**)

A single clone of *E. coli* K12 Δ*Opp*A cells transformed with a pSIJ8 plasmid was cultured in 2-YT supplemented with Ampicillin (50 µg/mL) and grown at 30°C, 200 rpm until an $OD_{600}$ of 0.3 followed by induction with 15 mM arabinose for the expression of lambda red recombineering genes . After incubation for 45 min at 37 °C the culture was cooled on ice to halt growth and made electrocompetent via a standard protocol. The resulting electrocompetent *E. coli* K12 Δ*Opp*A cells with expressed recombineering genes were then transformed with the linear DNA fragment encoding *opp*A ev2. After recovery in 1mL SOC for 2 h at 37 °C, cells were diluted in 2-YT and grown overnight at 37 °C for the curing of thermosensitive plasmid pSIJ8. The overnight culture (containing a mix of Δ*opp*A and knock-in cells) was diluted and grown to an $OD_{600}$ of 0.6 and made electrocompetent. These cells were then co-transformed with aaRS plasmid pEVOL_*Mb*PylRS and sfGFP reporter pBAD_sfGFP_N150TAG_H6. Transformed cells were recovered in 1 mL SOC for 1 h at 37 °C and diluted in 2-YT supplemented with Ampicillin (100 µg/mL) and Chloramphenicol (50 µg/mL) and grown to an OD600 of 0.6 and sfGFP expression was induced with 0.05% arabinose and 0.5 mM G-

SisoK was added. The culture was grown for 4 h, cooled on ice, centrifuged (4000 x g, 5 minutes), and resuspended in ice cold PBS. Cells with the highest sfGFP fluorescence were sorted as single cells into a 96 well plate containing 2-YT and grown overnight. Clones with the correct genomic insert were confirmed with sequencing of the genomic locus and whole genome sequencing. Once confirmed, plasmids were cured from the strain via electroporation.

**On bead CuAAC labeling of eGFP-NB with Picolyl-Azide-Sulfo-Cy5**

eGFP-NB bearing either BocK, XisoK (Prg) or PraK[5] at position R75 was expressed and purified as described in '**Expression of XisoK bearing proteins**' and '**Purification of His6 tagged proteins**'. Purified proteins were buffer exchanged into PBS using Amicon® centrifugal filter units (Millipore, 3 kDa MWCO), and 20 µM of each eGFP-NB variant was bound to magnetic Ni-NTA beads (Cube biotech) . Beads were washed 2 times with PBS and then incubated with CuAAC mix containing 100 µM AF647-Picolyl-Azide (Jena Bioscience), 50 µM $CuSO_4$, 250 µM THPTA and 2.5 mM ascorbic acid in PBS for 30 minutes at room temperature. Beads were then washed 5 times with PBS and protein was eluted with 300 mM imidazole in PBS. Eluted protein and flow-through samples were run on an SDS-PAGE and labeling visualized via in-gel fluorescence (iBright™ FL1500, ThermoFisher Scientific).

**Photocrosslinking of diazirine bearing proteins in cells.**

Chemically competent *E. coli* K12 cells were co-transformed with pBAD_POI_His6 (either sfGFP or GST encoding a protein of interest with a C-terminal His6 tag and a TAG codon at the indicated position) and pEVOL_*Ma*PylRS_IP (encoding two copies of *Ma*PylRS H227I, Y228P and *Ma* $tRNA_{CUA}$) for incorporating pLisoK or pEVOL_DiazKRS (encoding two copies of DiazKRS[6] and *Mm* $tRNA_{CUA}$). After recovery in 1 mL SOC for 1 h at 37 °C, cells were added to 5 mL of 2-YT supplemented with Ampicillin (100 µg/mL) and Chloramphenicol (50 µg/mL) and grown overnight at 37 °C. Autoinducing media supplemented with Ampicillin (100 µg/mL) and Chloramphenicol (50 µg/mL) and containing either no non-canonical amino acid, 2mM BocK or 2mM G-pLisoK were inoculated with the overnight culture and grown at 37 °C overnight. The resulting culture was diluted to an $OD_{600}$ of 1 and +UV samples were irradiated with UV light (15 W, 365 nm) using a lamp (Vilber, VL-215.L) for 15 minutes. +UV and -UV samples were run on an SDS-PAGE and analyzed via western blot using an anti-His peroxidase-coupled antibody (Sigma).

**Tyrosinase-mediated labelling of PisoK bearing proteins**

20 µM of protein (either Ub wt, UbK63PisoK or 3C-UbK63PisoK) was incubated with 160 µM p-cresol and 0.4 µM tyrosinase in tyrosinase buffer (20 mM potassium phosphate buffer pH 6.5, 100 mM NaCl) and incubated at room temperature for 2 hours. Reaction was quenched with 1 mM TCEP and 1 mM BCN-OH (Synaffix) for 10 minutes. Cleavage of the 3C-UbK63PisoK construct N-terminus was done with HRV 3C-protease (ThermoFischer) to obtain single-labeled protein.

**Chemical crosslinking of protein-protein complexes using ClAisoK in living *E. coli***

Affibody-ProteinZ:

Chemically competent *E. coli* K12 cells were co-transformed with pBAD_Affibody_D36TAG_His6, pBAD_RSF1035_Strep_SUMO_ProteinZ_N7C and pEVOL_*Mb*PylRS_C313V (encoding two copies of *Mb* PylRS with a C313V mutation and *Mm* $tRNA_{CUA}$) or pEVOL_wt_*Mb*PylRS. After recovery in 1 mL SOC for 1 h at 37 °C, cells were added to 5 mL of 2-YT supplemented with Ampicillin (100 µg/mL), Chloramphenicol (50 µg/mL) and Kanamycin (50 µg/mL) and grown overnight at 37 °C, 200 rpm. Autoinducing media (2 mL cultures in a 24-well plate) [13] supplemented with Ampicillin (100 µg/mL), Chloramphenicol (50 µg/mL) and Kanamycin (50 µg/mL) and containing either no non-canonical amino acid, 2 mM G-ClAisoK or 2 mM G-AisoK were inoculated with the overnight culture and grown at 37 °C, 200 rpm overnight.
After overnight incubation, $OD_{600}$ was measured, samples were normalized and subjected to SDS-PAGE followed by either Coomassie staining or western blot analysis using an anti-His peroxidase-coupled antibody (Sigma) or antiStrep-HRP (StrepMAB-Classic HRP, IBA, Cat.No. 2-1509-001).

sfGFP dimer:

Chemically competent *E. coli* K12 cells were co-transformed with pBAD_sfGFP_N150TAG_His6 (and mutants E173C, E173H, E173A or V207K) and pEVOL_*Mb*PylRS_C313V (encoding two copies of *Mb* PylRS with a C313V mutation and *Mm* $tRNA_{CUA}$) or pEVOL_wt_*Mb*PylRS. Expression and downstream analysis were performed analogously to the above-described procedure for Affibody-ProteinZ.

### Rab1b-DrrA

Chemically competent *E. coli* K12 cells were co-transformed with pBAD_Rab1b_R79TAG_His6_RBS_Strep_TEV_DrrA(339-522)_D512C (encoding both target genes in a polycistronical manner) and pEVOL_*Mb*PylRS_C313V (encoding two copies of *Mb* PylRS with a C313V mutation and *Mm* $tRNA_{CUA}$) or pEVOL_wt_*Mb*PylRS. Expression and downstream analysis were performed analogously to the above-described procedure for Affibody-ProteinZ. Ni-NTA purification was performed as described in “Purification of His6 tagged proteins”.

**Chemical crosslinking of Affibody and ProteinZ in vitro**

Affibody bearing either AisoK or ClAisoK at position 36 and a SUMO-ProteinZ N7C mutant were expressed and purified as described in sections “Expression of XisoK bearing proteins” and “Purification of His6 tagged proteins”. For in vitro crosslinking assays Affibody variants and the SUMO-ProteinZ N7C mutant were diluted to 50 µM each into PBS pH 7.4 supplemented with 1 mM TCEP and incubated at 37 °C. At denoted time points, 2 µL samples were taken and quenched by the addition of 8 µL 4× SDS loading buffer. After boiling at 95 °C for 10 min, the samples were loaded on SDS–PAGE and visualized by Coomassie staining.

**Determination of doubling times of isoK12 and K12 in AI and 2-YT media**

Single colonies of *E. coli* K12 and GXK were cultured in 20 ml of 2-YT and grown to an $OD_{600}$ of 0.6 at 37 °C, 200 rpm. These culture were then diluted to an $OD_{600}$ of 0.05 in pre-warmed 2-YT or autoinduction media and cultured at 37 °C, 200 rpm while monitoring $OD_{600}$ at regular intervals. To determine doubling times the slope of time (minutes) vs log2($OD_{600}$) at the exponential phase was calculated. Analysis was done using GraphPad Prism Version 10).

**Platereader based sfGFP fluorescence measurements**

sfGFP fluorescence time courses of *E. coli* cultures were measured on a Tecan Spark® Multimode Microplate Reader with a humidity cassette in order to prevent evaporation. *E. coli* cultures containing the appropriate plasmids were diluted to an $OD_{600}$ of 0.05 in the relevant media and grown to an $OD_{600}$ of 0.6 and sfGFP expression was induced with 0.05% arabinose and non-natural amino acid added to the media. 200 uL of this culture was added to the well of a 96-well microplate with a clear bottom (Greiner Bio-One µClear™ Cat.no. 655096) and the plate sealed with a breathe-EASIERä (Diversified Biotech) membrane. sfGFP fluorescence was measured from the bottom every 10 minutes while shaking in-between reads at 37 °C.

**Dual stop codon suppression for incorporation fo AcK and pLisoK into proteins**

Chemically competent *E. Coli* K12 cells were co-transformed with pBAD_POI (either pBAD_sfGFP_N40TAA_N150TAG_H6 or pBAD_Ub_K48TAA_TEV_SUMO2_K11TAG_H6) with a C-terminal His6 tag) and pEVOL_AcKRS3(TAA)_RBS_MaPylRS_IP(TAG) (encodes AcKRS3 and MaPylRS_IP polycistronically and their respective tRNAs). After recovery in 1 mL SOC media for 1 h at 37 °C, cells were cultured in 5 ml of 2-YT supplemented with Ampicillin (100 µg/mL) and Chloramphenicol (50 µg/mL) and incubated overnight at 37 °C, 200 rpm. The overnight culture was then diluted to an $OD_{600}$ of 0.05 in autoinducing media supplemented with Ampicillin (100 µg/mL), Chloramphenicol (50 µg/mL) and respective ncAA and/or peptide. Cells were grown overnight at 37 °C and the overnight culture was harvested by centrifugation at 4000 x g for 10 minutes and the pellets were stored at -20 °C till further use.

Ub-K48pLisoK-TEV-SUMO2-K11AcK-H6 was purified as described in '**Purification of His6 tagged proteins**'. For cleavage, Ub-K48pLisoK-TEV-SUMO2-K11AcK-H was diluted into buffer (PBS with 3 mM DTT) and incubated with TEV protease (0.1 mg/ml) for 30 minutes at RT.

### References

1 Klepsch, M. M. *et al.* Escherichia coli peptide binding protein OppA has a preference for positively charged peptides. *J Mol Biol* **414**, 75-85 (2011). https://doi.org/10.1016/j.jmb.2011.09.043

2 Serfling, R. *et al.* Designer tRNAs for efficient incorporation of non-canonical amino acids by the pyrrolysine system in mammalian cells. *Nucleic Acids Res* **46**, 1-10 (2018). https://doi.org/10.1093/nar/gkx1156

3 Tai, J. *et al.* Pyrrolysine-Inspired in Cellulo Synthesis of an Unnatural Amino Acid for Facile Macrocyclization of Proteins. *J Am Chem Soc* **145**, 10249-10258 (2023). https://doi.org/10.1021/jacs.3c01291

4 Zang, J. *et al.* Genetic code expansion reveals aminoacylated lysine ubiquitination mediated by UBE2W. *Nat Struct Mol Biol* **30**, 62-71 (2023). https://doi.org/10.1038/s41594-022-00866-9

5 Nguyen, D. P. *et al.* Genetic Encoding and Labeling of Aliphatic Azides and Alkynes in Recombinant Proteins via a Pyrrolysyl-tRNA Synthetase/tRNACUA Pair and Click Chemistry. *Journal of the American Chemical Society* **131**, 8720-8721 (2009). https://doi.org/10.1021/ja900553w

6 Nguyen, T. A., Gronauer, T. F., Nast-Kolb, T., Sieber, S. A. & Lang, K. Substrate Profiling of Mitochondrial Caseinolytic Protease P via a Site-Specific Photocrosslinking Approach. *Angew Chem Int Ed Engl* **61**, e202111085 (2022). https://doi.org/10.1002/anie.202111085

7 Yang, F., Moss, L. G. & Phillips, G. N. The molecular structure of green fluorescent protein. *Nature Biotechnology* **14**, 1246-1251 (1996). https://doi.org/10.1038/nbt1096-1246

8 Zacharias, D. A., Violin, J. D., Newton, A. C. & Tsien, R. Y. Partitioning of lipid-modified monomeric GFPs into membrane microdomains of live cells. *Science* **296**, 913-916 (2002). https://doi.org/10.1126/science.1068539

9 Hogbom, M., Eklund, M., Nygren, P. A. & Nordlund, P. Structural basis for recognition by an in vitro evolved affibody. *Proc Natl Acad Sci U S A* **100**, 3191-3196 (2003). https://doi.org/10.1073/pnas.0436100100

10 Cigler, M. *et al.* Proximity-Triggered Covalent Stabilization of Low-Affinity Protein Complexes In Vitro and In Vivo. *Angew Chem Int Ed Engl* **56**, 15737-15741 (2017). https://doi.org/10.1002/anie.201706927

11 Schoebel, S., Oesterlin, L. K., Blankenfeldt, W., Goody, R. S. & Itzen, A. RabGDI Displacement by DrrA from Legionella Is a Consequence of Its Guanine Nucleotide Exchange Activity. *Molecular Cell* **36**, 1060-1072 (2009). https://doi.org/https://doi.org/10.1016/j.molcel.2009.11.014

12 Chiu, J., March, P. E., Lee, R. & Tillett, D. Site-directed, Ligase-Independent Mutagenesis (SLIM): a single-tube methodology approaching 100% efficiency in 4 h. *Nucleic Acids Res* **32**, e174 (2004). https://doi.org/10.1093/nar/gnh172

13 Muzika, M. *et al.* Chemically-defined lactose-based autoinduction medium for site-specific incorporation of non-canonical amino acids into proteins. *RSC Adv* **8**, 25558-25567 (2018). https://doi.org/10.1039/c8ra04359k

14 Huguenin-Dezot, N. *et al.* Trapping biosynthetic acyl-enzyme intermediates with encoded 2,3-diaminopropionic acid. *Nature* **565**, 112-117 (2019). https://doi.org/10.1038/s41586-018-0781-z
